# Synthetic hybrids of six yeast species

**DOI:** 10.1101/597633

**Authors:** David Peris, William G. Alexander, Kaitlin J. Fisher, Ryan V. Moriarty, Mira G. Basuino, Emily J. Ubbelohde, Russell L. Wrobel, Chris Todd Hittinger

**Author notes:** Correspondence to (CTH), (DP).

## Abstract

Allopolyploidy generates diversity by increasing the number of copies and sources of chromosomes. Many of the best-known evolutionary radiations, crops, and industrial organisms are ancient or recent allopolyploids. Allopolyploidy promotes differentiation and facilitates adaptation to new environments, but the tools to test its limits are lacking. Here we develop an iterative method to combine the genomes of multiple budding yeast species, generating *Saccharomyces* allopolyploids of an unprecedented scale. Chromosomal instability and cell size increased dramatically as additional copies of the genome were added, but we were able to construct synthetic hybrids of up to six species. The six-species hybrids initially grew slowly, but they rapidly adapted when selection to a novel environment was applied, even as they retained traits from multiple species. These new synthetic yeast hybrids have potential applications for the study of polyploidy, genome stability, chromosome segregation, cancer, and bioenergy.

**One sentence summary:** We constructed six-species synthetic hybrids and showed that they were chromosomally unstable but able to adapt rapidly.

## Introduction

Polyploidy generates diversity by increasing the number of copies of each chromosome ^1^. Allopolyploidy instantly adds chromosomal variation from multiple species through hybridization, while autopolyploidy leads to variation as gene copies from a single species diverge during evolution. Allopolyploidy facilitates differentiation and adaptation to new environments ^2^. Many plants, animals, fungi, and other eukaryotes are ancient or recent allopolyploids, including some of the best-known industrial organisms, crops, and evolutionary radiations ^3, 4^.

Phylogenomic analyses support an allopolyploid origin for the baker’s yeast *Saccharomyces cerevisiae* ^5^. *S. cerevisiae* has been one of the most important model organisms to study polyploidy in the context of evolution ^6^, its effects on mutation rate ^7^, and as a model of how cancer progresses as clonal populations adapt through driver mutations ^8^. Despite the decreased fitness of newly generated polyploids ^9^, experimental evolution assays in *S. cerevisiae* and comparisons of the genomes of industrial *Saccharomyces* interspecies hybrids have shown that they return to high fitness through many of the same genetic mechanisms that occur during the clonal expansion of tumorigenic cells, such as aneuploidy, chromosomal rearrangements, and loss-of-heterozygosity ^10, 11^.

Genome rearrangements are common in *Saccharomyces* allopolyploids used to make fermented beverages ^12, 13^, but experimental tools to test the limits of polyploidy and genome rearrangements are lacking. Random chromosomal aberrations can be easily generated by using molecular techniques, such as SCRaMbLE ^14^. However, SCRaMbLE is currently only available in single, partly synthetic *S. cerevisiae* strain, limiting the genomic diversity that can be explored.

*Saccharomyces* species have similar genome content, identical numbers of chromosomes (n = 16), and genomes that are mostly syntenic ^15^. Since they have limited pre-zygotic barriers, interspecies hybrids can be generated easily when haploid strains of opposite mating types encounter each other. Much more rarely, diploid yeast cells can become competent to mate by inactivating or losing one *MAT* idiomorph or undergoing gene conversion at the *MAT* locus ^16^. To facilitate the generation of allododecaploid (base ploidy of 12n) hybrids of six species, we developed an iterative <u>Hy</u>brid <u>Pr</u>oduction (iHyPr) method. iHyPr combines traits from multiple species, such as temperature tolerance, and through adaptive laboratory evolution, facilitates rapid adaptation to new environments. This new method will enable basic research on polyploidy, cancer, and chromosome biology. iHyPr can further be applied to research on bioenergy and synthetic biology, as genomic diversity can be harnessed to generate more efficient strains that produce new bioproducts ^17^ or to combine industrially useful traits from multiple species ^18, 19^.

## Results

### Synthetic hybrids of six yeast species can be generated with iHyPr

iHyPr allowed us to experimentally test the limits of chromosome biology and allopolyploidy by constructing a series of higher-order interspecies hybrids (Supplementary Fig. 1). First, we used two differentially marked HyPr plasmids, which each encode a drug-inducible *HO* gene (*<u>ho</u>*mothallic switching endonuclease) that promotes mating-type switching, to efficiently generate and select for two-species hybrids as done previously ^20^. Next, using two newly created, differentially marked HyPr plasmids, we crossed these two-species hybrids to construct three-species and four-species hybrids. The construction of higher-order synthetic hybrids has not been reported previously. Finally, we constructed six-species hybrids using three different crossing schemes (Figure 1, Supplementary Fig. 2).

**Figure 1.**
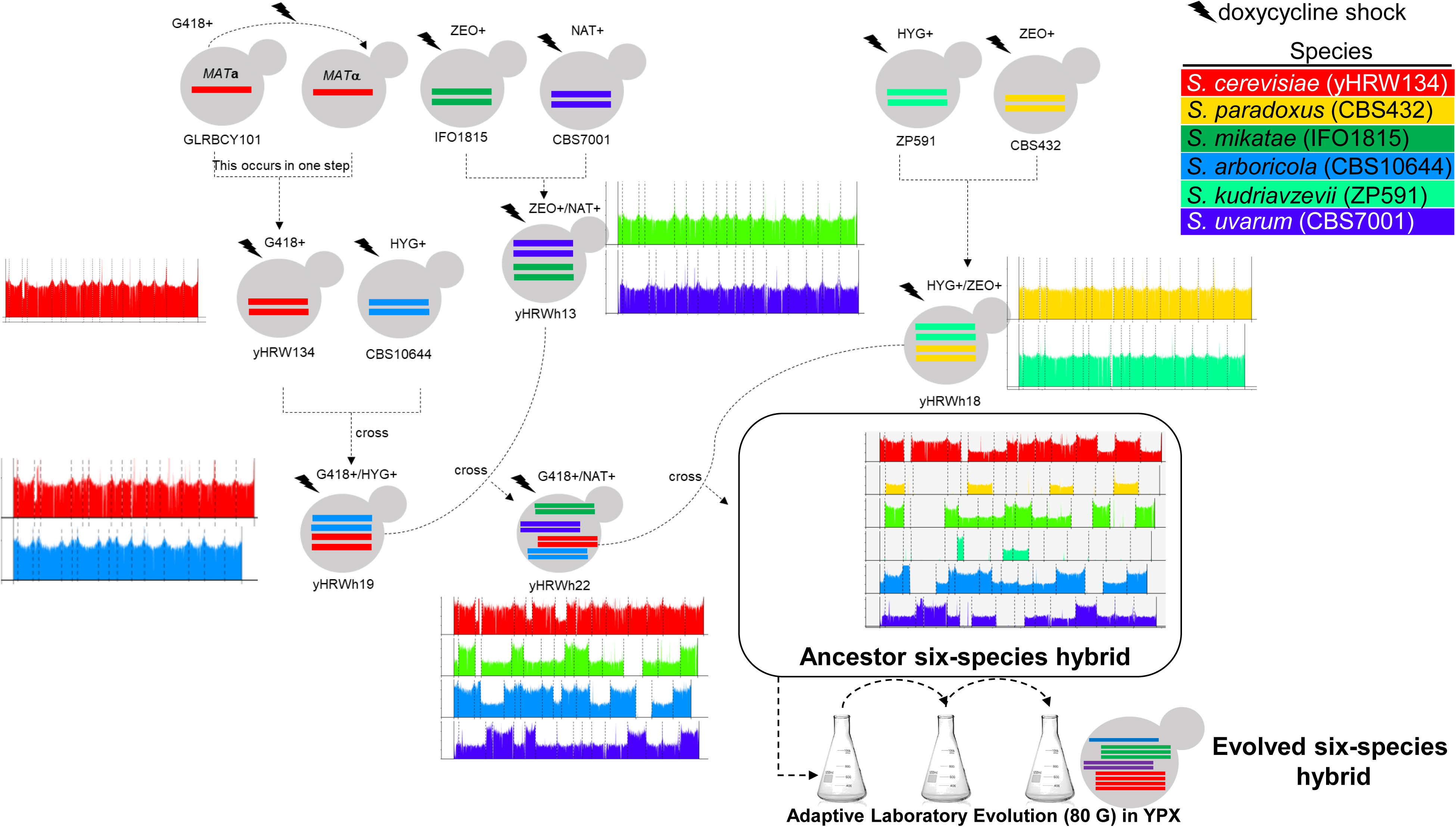
The generation of ancestor and evolved six-species hybrids. Synthetic hybrid generation scheme using the iHyPr method. The example shown is the six-species hybrid yHRWh39. Chromosomes were colored according to their species designation, with height representing copy number, using the `sppIDer` pipeline ^50^. For an extended explanation of iHyPr, including the other two crossing schemes, see Supplementary Fig. 1, 2. Arrows mark hybridization steps. For additional intermediate and six-species hybrid nuclear and mitochondrial genomes with higher resolution, see Supplementary Fig. 3, 4. Ancestor six-species hybrids underwent ALE for 80 <u>G</u>enerations.

In all three schemes, diploid genomes were successfully introduced from each of the six parent species (Figure 2B, Supplementary Fig. 3). During hybrid construction, as more and more genomes were introduced, the frequency of successful matings decreased (Spearman rank sum test R = −0.89, *p*-value = 1.1*10^-5^, Figure 3A, Supplementary Table 3), and the fitness of synthetic hybrids declined (Spearman rank sum test = −0.77, *p*-value = 7.6*10^-4^, Figure 3C). The fitness decrease may be due to the increased cell area, which is correlated with the increased genome size (Spearman rank sum test R = 0.97, *p*-value = 1.4*10^-5^, Figure 3B), as well as interspecies genetic incompatibilities.

**Figure 2.**
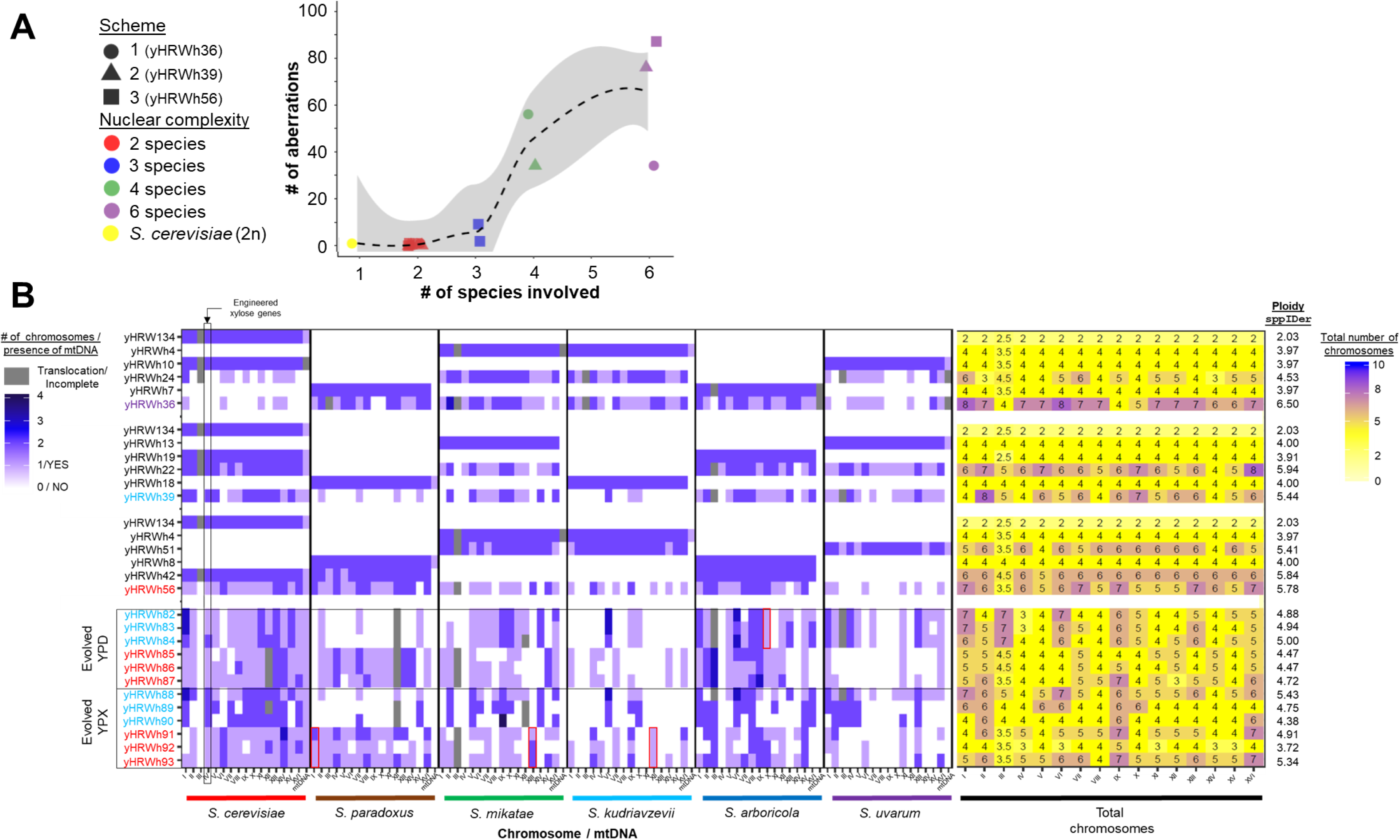
Genome contributions to synthetic hybrids. The numbers and sources of chromosomes for each synthetic hybrid were inferred from `sppIDer` plots (Supplementary Fig. 3), which were corrected based on flow cytometry ploidy estimations. A) The number of chromosomal aberrations were inferred for each synthetic hybrid as new translocation, gain, and loss events not seen in the preceding hybrid (Supplementary Fig. 3). Chromosomal aberrations involving parts of chromosomes were conservatively counted only in cases of clear fusion of entire arms, whereas smaller loss-of-heterozygosity events were not counted. The synthetic hybrids generated from each independent scheme are represented with different shapes. Color points are colored according to the number of species genomes contributing to the strain. A LOESS regression line and the 95% confidence interval of the fit are represented with a discontinuous black color and gray shadow, respectively. B) Chromosome content was colored according to the species donor. Mitochondrial inheritance was inferred using `mitosppIDer` (Supplementary Fig. 4). The numbers of chromosomes for each species are colored according to the left heatmap legend. Incomplete and recombinant mtDNA are colored in gray. Total number of chromosomes is shown in the right part of the figure, which is colored according to the right legend. Ploidy estimates based on de novo genome assemblies, which correlates with flow cytometry (Spearman rank sum test R = 0.88, *p*-value = 7.5*10^-8^, Supplementary Fig. 5C), are indicated at the right side of the figure. Synthetic hybrids are reported in the order constructed (Supplementary Fig. 2). Diploidized GLRBCY101 (yHRW134) and yHRWh4 are shown multiple times because of their use in multiple crossing schemes. Evolved hybrids are grouped based on the conditions in which they were evolved, and they are colored according to their ancestor hybrid. Red squares highlight chromosomes that were retained or lost in all hybrids evolved in the same condition when compared to their siblings evolved in the other condition. *S. cerevisiae* chromosome IV, where the xylose utilization genes were inserted, is indicated by the black square. Note that considerable karyotypic diversity continued to be generated during 80 generations of ALE (Figure 6), but each evolved strain is easily recognized as more similar to its ancestor six-species hybrid.

**Figure 3.**
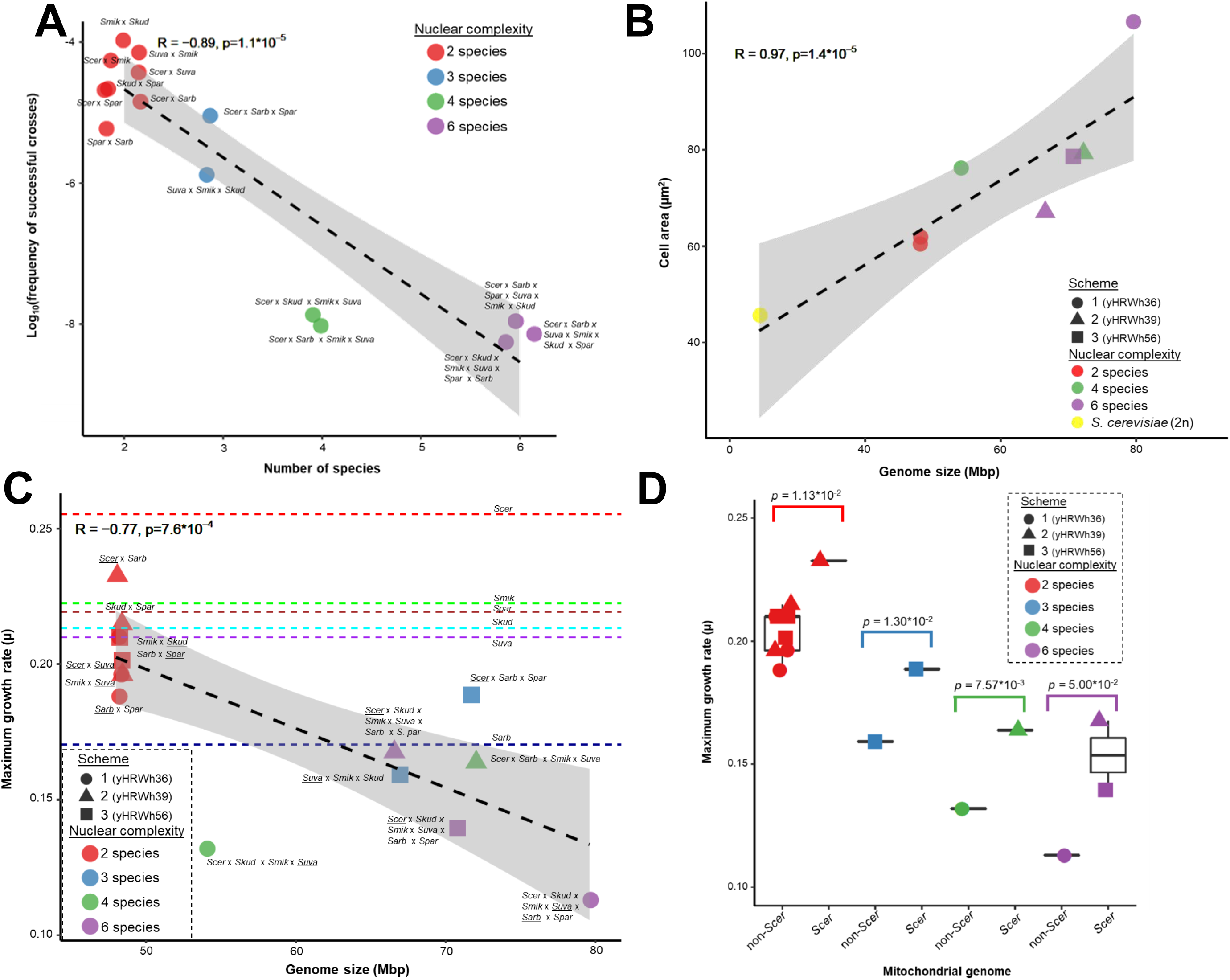
Characteristics of six-species hybrids. A) The number of species contributing genomes to synthetic hybrids is inversely correlated with the frequency of successful matings. B) Genome size is correlated with average cell area (n = 36-78 counted cells). C) Genome size (Supplementary Table 2) versus the average maximum growth rate (µ (n=6), defined as (ln(OD2)-ln(OD1))/(T2-T1)) in rich medium at 20 °C (Supplementary Table 4). Dashed lines are the µ for the parent species indicated close to the line. For *S. uvarum*, the average of two strains with different HyPr plasmids is shown. D) The maximum specific growth rate (µ, defined as (ln(OD2)-ln(OD1))/(T2-T1))) in rich medium at 20°C is higher in interspecies hybrids inheriting *S. cerevisiae* mtDNA. Colors correspond to the number of species contributing genomes to each strain. Synthetic hybrids generated from independent schemes are represented by different shapes in panels B), C), and D). The Spearman rank sum test R and *p*-value are displayed. A linear regression and its 95% confidence interval are represented with a black dashed line and gray shadow, respectively. The mtDNA donor is underlined in the names in panel C). Species composition abbreviations are: *Scer*, *S. cerevisiae*; *Spar*, *S. paradoxus*, *Smik*, *S. mikatae*, *Sarb*, *S. arboricola*; *Skud*, *S. kudriavzevii*; and *Suva*, *S. uvarum*.

### Genome size and stability limits

The largest synthetic hybrid expanded its genome size 3.3 times (from 24 Mb to ∼80 Mbp) (Supplementary Fig. 5A, Supplementary Table 2), and its cell area was 2.3 times larger than a diploid cell (Figure 3B). Some species contributed many fewer chromosomes than others to the synthetic hybrids of six species (defined as the ancestor hybrids) (Figure 2B). During construction, chromosome losses were widespread and outnumbered gains (two-sided t-test t = −3.4408, d.f. = 6, *p*-value < 1.37*10^-2^, Supplementary Table 2). These aneuploidies rose dramatically as the number of species donating genomes increased (linear regression r^2^ = 0.79, *p*-value = 3.01*10^-6^) (Figure 2A). Complete chromosomal aneuploidies were much more common than aneuploidies caused by unbalanced translocations or deletions (97.31% versus 2.69% of total detected chromosomal aberrations). Aneuploidies involving chromosome III, where 88.9% of translocations or deletions in this chromosome were unbalanced (Figure 2B, Supplementary Fig. 3), were especially common because it contains the *MAT* locus being cut by the Ho endonuclease during iHyPr. These results suggest that, in addition to the expected gene conversion events, iHyPr generated mating-competent cells via partial chromosome losses.

Even though the base ploidy of the final six-hybrids was allododecaploidy (12n), a ploidy level acquired by only a few organisms ^21, 22^, none were euploid (Figure 2B). Due to massive chromosome loss, we inferred the six-species hybrid with the largest genome had an average of ∼7 copies of each chromosome (i.e. 12n – 88) when estimated using bioinformatic tools (visual inspection of `sppIDer` plots) or an average of ∼8 copies of each chromosome (i.e. 12n – 64) when total DNA content was estimated using flow cytometry (Figure 2B, Supplementary Fig. 3, 5C, Supplementary Table 2).

### Mitochondrial inheritance affects genotype and phenotype

During interspecific hybridization, hybrids can inherit one of the two parent mitotypes or a recombinant version depending on the budding location ^23^. In general, one of the parent mitotypes was quickly fixed during the generation of our hybrids, except for three cases: the allotetraploid *Saccharomyces kudriavzevii* x *Saccharomyces mikatae* yHRWh4, the allotetraploid *S. cerevisiae* x *Saccharomyces uvarum* yHRWh10, and the six-species hybrid yHRWh36, which were all heteroplasmic (Supplementary Fig. 4). In rich medium at 20 °C, for strains with similar numbers of hybridized species, hybrids with a *S. cerevisiae* mitochondrial genome (mtDNA) grew 7-15 % faster than the hybrids with the mtDNA of another species (Figure 3C,D). mtDNA inheritance was also significantly correlated with nuclear genome retention (ANOVA multifactor *F*-value = 19.9, d.f. = 1, *p*-value = 7.77*10^-4^), with the mtDNA donor tending to contribute more nuclear chromosomes (Figure 4, Supplementary Fig. 4). These results are consistent with recent observations in hybrids used in the fermented beverage industry ^12^.

**Figure 4.**
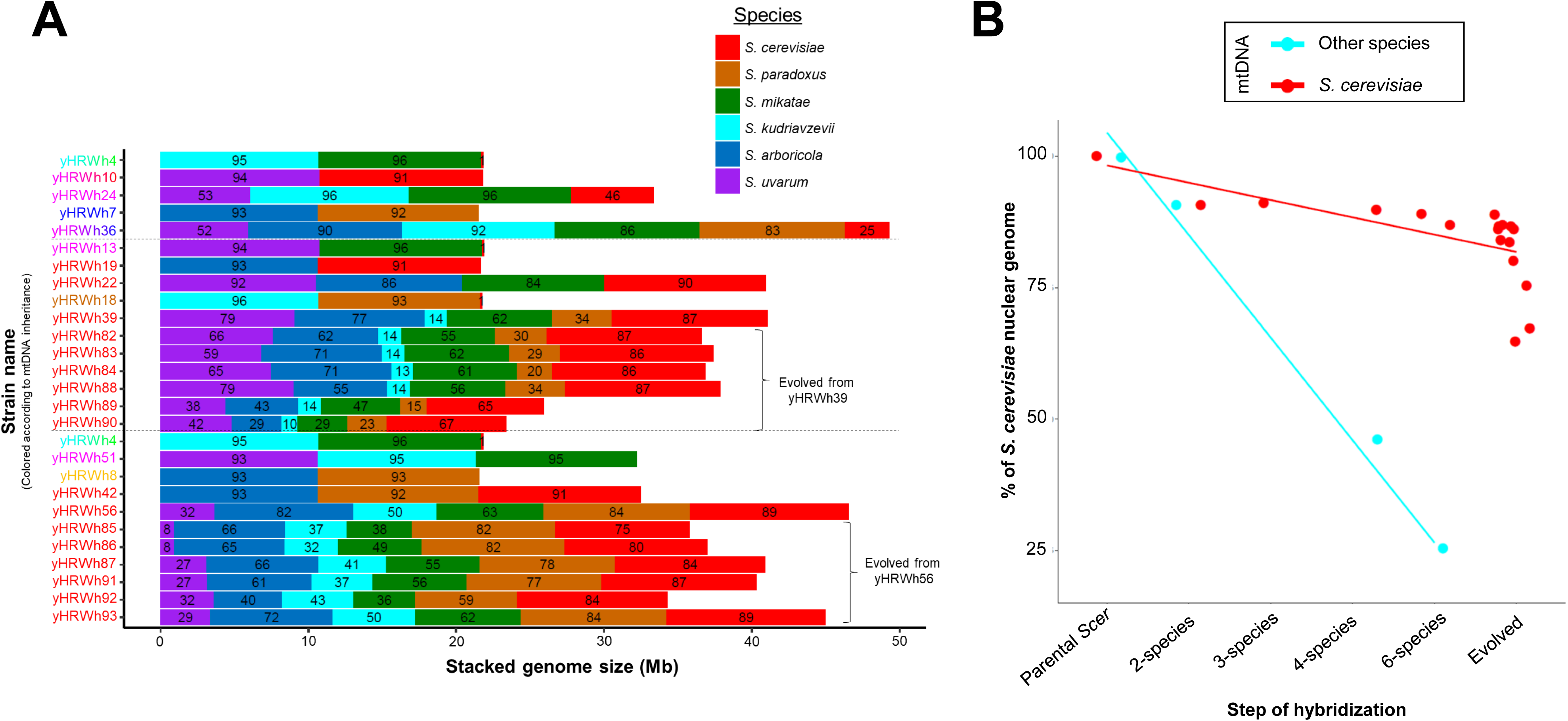
Genome reduction during hybrid construction and adaptive laboratory evolution. A) The genome contribution of each *Saccharomyces* species is stacked, and the percentage of retention is indicated inside the bar plot for each synthetic hybrid. Presence is reported, not copy number. Synthetic hybrids are displayed in the order constructed (Supplementary Fig. 2). yHRWh4 is shown multiple times because of its use in two crossing schemes. We did not expect 100% genome contribution for each *Saccharomyces* species, even for recently created hybrids, because some genomic regions (e.g. repeats) are not unambiguously detectable with Illumina sequencing data. Genome size bars are colored according to each species’ contribution. The strain names are colored based on the mtDNA inheritance inferred from `mitosppIDer` (Supplementary Fig. 4), with two or more mtDNAs or regions shown as a gradient. B) The nuclear compositions of the *S. cerevisiae* parent, synthetic hybrids, and evolved hybrids are plotted according to mtDNA inheritance. Hybrids with *S. cerevisiae* mtDNA or with other mtDNA are colored in red and light blue, respectively.

### Trait combination, adaptive laboratory evolution, and genome stabilization

Higher-order synthetic hybrids allow investigators to rapidly combine traits from many different parents, such as differences in sugar consumption and temperature preferences. To determine if the inherent chromosomal instability of these six-species hybrids could be harnessed as a diversity generator, we tested how these new six-species hybrids altered their kinetic parameters during adaptive laboratory evolution (ALE). ALE was performed for an estimated 80 generations in a medium containing glucose or xylose, a sugar poorly metabolized by most *Saccharomyces* species ^24^. To provide baseline xylose metabolic capability upon which to improve, we chose a *S. cerevisiae* parent strain that had been engineered by inserting xylose utilization genes into Chromosome IV ^25, 26^. Ancestor six-species hybrids grew slowly, and despite differing from each other in chromosomal composition (Figure 2B, 4A), single-colony isolates of all 12 ALE replicates (3 replicates for the two ancestor hybrids retaining the chromosome IV in two ALE conditions) outperformed their ancestors in culturing conditions identical to the ALE (one-sided Wilcoxon rank sum test, *p*-value = 3.51*10^-4^). Many evolved strains even outperformed the *S. cerevisiae* reference strain (Figure 5A). In microtiter plate culturing conditions where more replicates could be achieved, evolved hybrid populations grew as much as 71% faster on xylose than the reference *S. cerevisiae* strain, and populations evolved on xylose outperformed those evolved on glucose (one-sided Wilcoxon rank sum test, *p-*value = 1.29*10^-2^) (Supplementary Fig. 6, Supplementary Table 6). Importantly, all our evolved hybrids grew well at low temperature conditions (4 °C) where the *S. cerevisiae* parent could not grow (Figure 5B), demonstrating that the cold tolerance of the other parents ^27, 28^ had been retained through hybridization and ALE.

**Figure 5.**
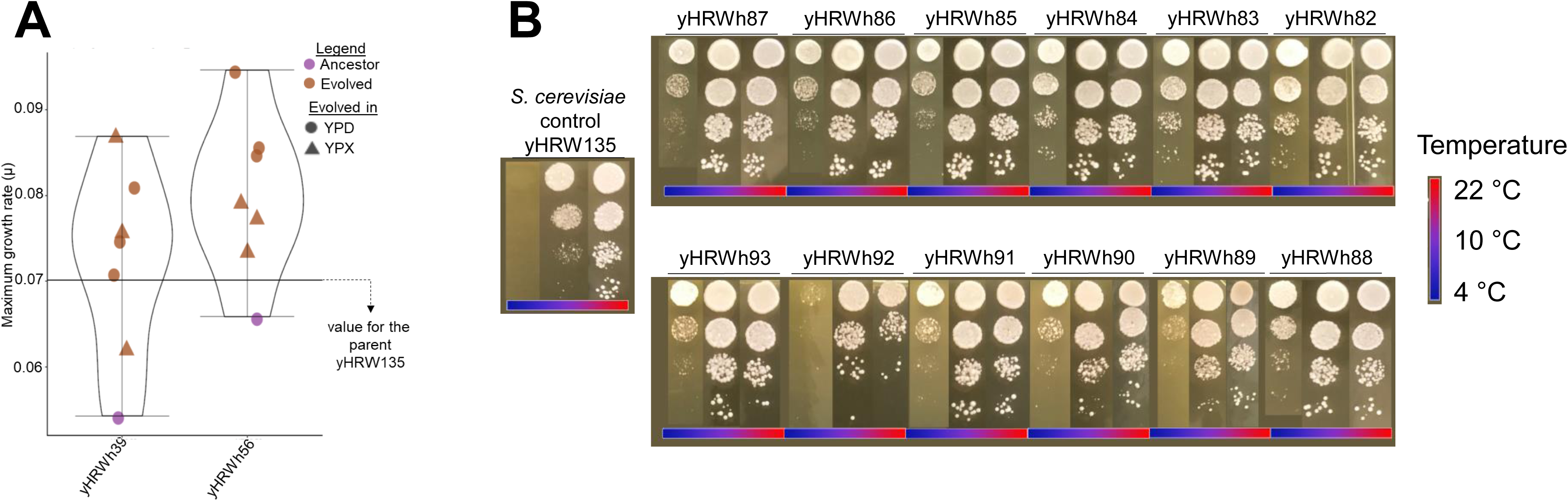
Trait combination and improvement by adaptive laboratory evolution. A) Box plots for the individual evolved colonies isolated from YPX or YPD plates after ALE and their synthetic hybrid ancestors. Kinetic parameters were tested in 3 ml YPX on a rotating culture wheel, identically to how they were evolved for 80 generations. The average values (n=6) of maximum specific growth rates (µ, defined as (ln(OD2)-ln(OD1))/(T2-T1)) for the *S. cerevisiae* reference strain (black line, yHRW135 was derived from yHRW134 by plasmid loss), ancestor six-species hybrids (purple dots), and evolved six-species hybrids (brown dots) are shown (Supplementary Table 5). Different shapes indicate the media in which the synthetic six-species hybrids were evolved. Additional kinetic parameters from microtiter plate experiments performed on evolved populations are shown in Supplementary Fig. 6 and Supplementary Table 6. B) Spot tests for three temperatures (22, 10, and 4 °C) are displayed for the evolved strains and the *S. cerevisiae* reference strain yHRW135. Evolved six-species hybrids retained the ability to grow at 4 °C, a trait not possessed by *S. cerevisiae*, despite the fact that it was not selected during ALE.

Since maximum growth rate on xylose improved considerably regardless of whether hybrids were evolved on xylose or glucose, we hypothesized several factors that could be responsible, such as xylose cassette amplification or genome stabilization. Neither chromosome IV nor the xylose utilization genes themselves were selectively amplified in either condition (Figure 2B, Supplementary Fig. 7, Supplementary Table 7). Evolved hybrids with more reduced genome sizes tended to have slightly higher fitness among ALE replicates, but the correlation was not significant (*p*-value 0.054) (Supplementary Fig. 8). These results suggest that which regions of the genome are lost or amplified as an allopolyploid genome stabilizes matters more for adaptation than total size, which may indicate an important role for the removal of genetic incompatibilities.

Although genome instability increased after each step during the construction of the six-species hybrids (Figure 2A), genome instability decreased after 80 generations of ALE (Figure 6A). Nonetheless, genome sequencing of a random selection of colonies from one of the evolved six-species hybrids demonstrated that genomic diversity was still being generated at a prodigious rate (Figure 6B, C). Thus, genome stabilization was ongoing.

**Figure 6.**
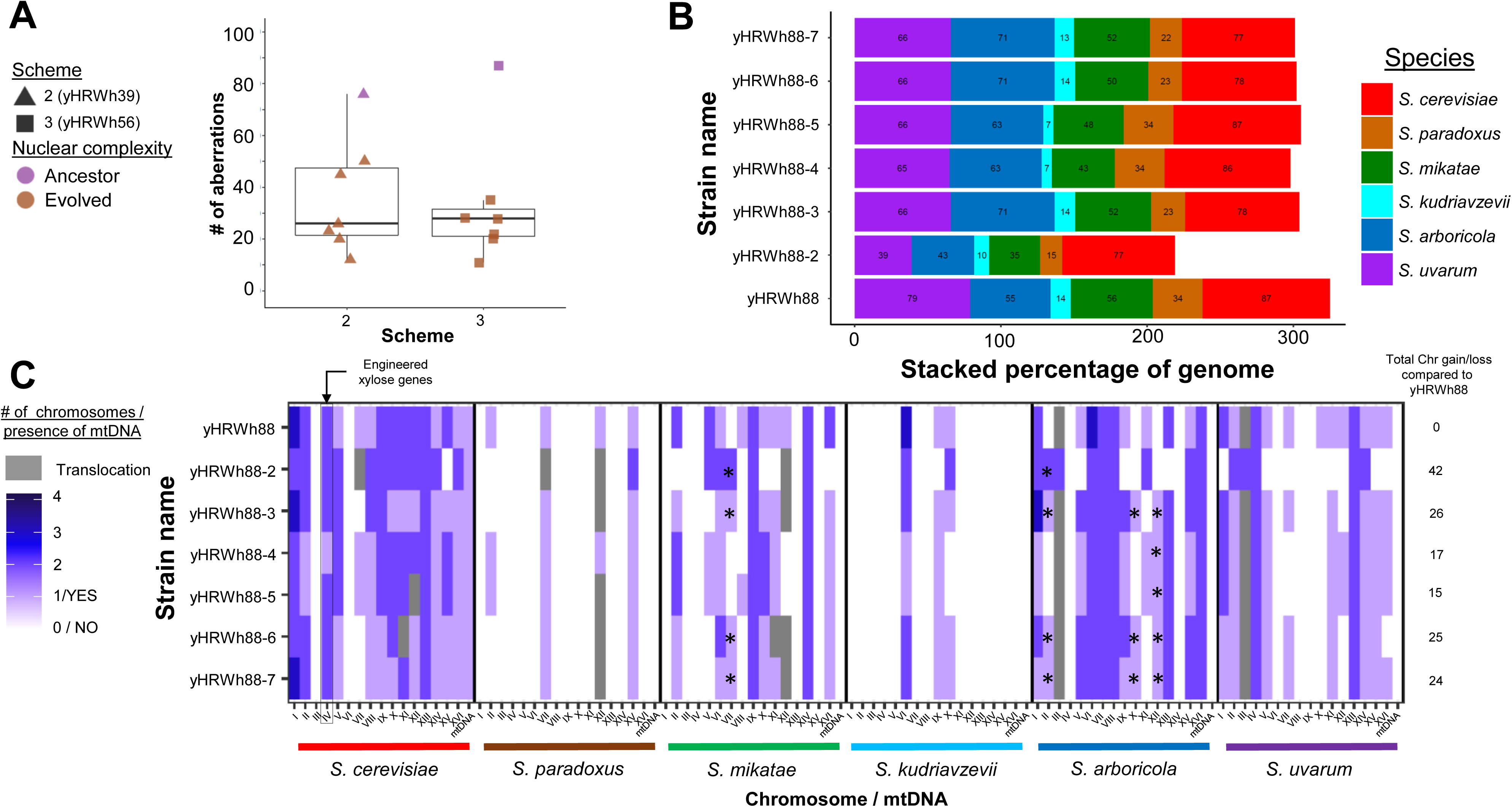
Synthetic hybrids as a tool to study genome instability. A) Boxplots of the number of chromosomal aberrations inferred for ancestor and evolved synthetic hybrids (Figure 2B, Supplementary Fig. 3). Synthetic hybrids generated from each independent scheme are represented with different shapes. Purple and brown color points represent whether six-species hybrids were ancestor or evolved, respectively. B) For each colony isolated from the population sample of the evolved synthetic hybrid yHRWh88, the genome contribution of each *Saccharomyces* species is stacked, and the percentage of retention is indicated inside the bar plot. The percentage of each species’ contribution are colored according to the legend. C) The number of chromosomes were inferred from `sppIDer` plots and corrected based on flow cytometry. The chromosome content was colored according to the species donor. The numbers of chromosomes for each species are colored according to the heatmap legend. Recombinant chromosomes are colored in gray. Asterisks indicate chromosomes that were retained in a particular colony but were not observed in the evolved yHWRh88 population sample, highlighting the instability of these hybrids. *S. cerevisiae* chromosome IV, where the xylose utilization genes were inserted, is indicated by the black square.

## Discussion

Collectively, our results show that iHyPr can generate and select for genome diversity, while combining industrially relevant traits from multiple parents. Specifically, we combined xylose utilization from a biofuel strain of *S. cerevisiae* with cold tolerance, a trait critical for the production of many fermented beverages ^29–31^.

### New *Saccharomyces* ploidy heights were reached using iHyPr

Previous efforts to generate higher ploidy *Saccharomyces* cells were arduous. A documented autohexaploid *S. cerevisiae* strain was produced by using a complex combination of auxotrophic intermediates ^9^, and allotetraploids of *S. cerevisiae* x *S. kudriavzevii* have been generated by using protoplast fusion and rare-mating ^32^. Recently, a CRISPR/Cas9 system was developed to switch mating-types and generate tetraploid yeast cells in a manner similar to HyPr ^33^. We show here that iHyPr can be used to produce higher-order hybrids iteratively without additional transformations. Although six-species hybrids were unstable and quickly lost chromosomes, they expanded yeast allopolyploidy to levels acquired by only a handful of plants and animals^21^.

iHyPr exploits the *Saccharomyces* mating system by heterologously expressing the *HO* gene from differentially marked plasmids ^20^. We expected that Ho would cut one copy of the heterozygous *MAT* locus of the diploid strain and use homology repair to convert the locus from heterozygous to homozygous, presumably over a small gene conversion patch using either a homologous chromosome or a silent mating cassette as the template. Although both this mechanism and larger breakage-induced replication events likely occur some of the time at this locus, 88.9% of the translocations or deletions involving chromosome III were unbalanced, suggesting other repair mechanisms are also leading to mating-type locus hemizygosity or homozygosity. For example, the high number of unbalanced translocations or deletions targeting chromosome III (40 %, or 6/15 hybrids) might support the occurrence of imperfect non-homologous end joining (NHEJ) events, perhaps promoted by overexpression of *HO* ^34^, or this chromosome might be inherently less stable ^35^. Synthetic hybrids between *S. cerevisiae* and *S. kudriavzevii* have demonstrated how easily chromosome III of one of the parents can be lost, rendering the hybrid competent to mate again ^36^. Recent studies of interspecies hybrids from the genus *Zygosaccharomyces* have also shown that inactivation of one of the *MAT* locus copies can also restore sexual competency ^37^. Regardless of the precise mechanisms at work, the iHyPr method clearly facilitated the iterative recovery of the sexual competency of higher-order hybrids by controlling and exploiting these naturally occurring mechanisms to generate interspecies hybrids with levels of allopolyploidy never seen in budding yeasts.

### iHyPr as a tool to study allopolyploids

The levels of allopolyploidy reached in our study will facilitate the understanding of the cellular fitness consequences in eukaryotes. The increase in ploidy was associated with a short-term fitness defect. However, ALE rapidly improved fitness, while allowing multiple parent traits, such as cold tolerance and xylose utilization, to be retained. Since both ALE selection regimes improved performance on xylose, improvements were likely driven partly by genome stabilization through the removal of interspecies genetic incompatibilities and partly due to condition-specific effects. The capacity of this approach to generate extensive karyotypic and phenotypic diversity will be of great interest for many industrial applications.

Mitochondrial inheritance also greatly influenced the genotypes and phenotypes of our synthetic hybrids. Even though a homoplasmic mtDNA state was quickly reached in most cases, a heteroplasmic state was detected in three exceptions that were all part of the same crossing scheme, and we offer a set of related possible explanations. The presence of selfish elements, such as homing endonucleases, could explain why multiple mitotypes were retained in yHRWh4. In this case, a portion of *S. mikatae COX1*, a gene with a high number of introns invaded by homing endonuclease genes ^38^, seems to have been introduced into the *S. kudriavzevii* mtDNA (Supplementary Fig. 4A,C). An even more intriguing result occurred while generating the yHRWh10 hybrid, which remained in a heteroplasmic state and retained most of the mtDNAs of both parents (*S. cerevisiae* and *S. uvarum*) (Supplementary Fig. 4A). We recently demonstrated that, during the formation of *S. cerevisiae* x *S. uvarum* hybrids, the frequency of strains without a functional mtDNA was higher when the hybrid inherited a *S. uvarum* mtDNA, but introgression of the F*-Sce*III homing endonuclease gene restored normal mitochondrial retention ^39^. Therefore, the absence of F*-Sce*III in yHRWh10 may have influenced the loss of mtDNA in its descendants, such as the six-species hybrid yHRWh36, which retained only small regions of *Saccharomyces arboricola* and *S. uvarum* mtDNAs (Supplementary Fig. 4C). In another recent study, mtDNA inheritance was dominated by one parent due to nuclear-mitochondrial interactions, rather than occurring stochastically ^40^.

The loss of mtDNAs in particular hybrid combinations, as well as the unusually high or low coverage in others (Supplementary Fig. 4), might further suggest that interactions between nuclear-encoded mitochondrial proteins with the mtDNA were unbalanced. In such cases, one model proposes that an oligomeric circular mtDNA form precedes mtDNA loss ^41^. Although technical artifacts from Illumina sequencing cannot be excluded, the read coverages for some regions of the mtDNAs were surprisingly varied in some hybrids, such as yHRWh8, yHRWh13, and most of the hybrids in the yHRWh36 crossing scheme (Supplementary Fig. 4). Formation and subsequent mis-regulation of mtDNA concatemers by Din7p and Mhr1p ^41^ provide a possible model for how specific mitochondrial regions increase or decrease in copy numbers in hybrids, and this phenomenon merits further study.

### Conclusions

In summary, we generated and extensively characterized two-, three-, four-, and six-species synthetic hybrids, using iHyPr. We also improved the fitness of evolved strains, which mimic the genetic processes seen in tumor cells escaping antitumorigenic treatments, where polyploidy drives genome instability and evolution ^8^. Our higher-order allopolyploids acquired genome aberrations involving multiple species as they rapidly adapted to new environmental conditions. This new technology pushes the budding yeast cell toward its limits in pursuit of basic research questions in chromosome biology and evolutionary genetics, as well as potential industrial applications.

## Methods

### Yeast strains and maintenance

The reference strain chosen for improvement was GLBRCY101, a haploid derivative of the *Saccharomyces cerevisiae* GLBRCY73 strain, which had been engineered with xylose utilization genes from *Scheffersomyces (Pichia) stipitis* and aerobically evolved for the consumption of xylose ^24–26^. Representative strains were selected from five additional *Saccharomyces* species based on published nuclear and mtDNAs (Supplementary Table 1). These six parent strains were used to generate the six-species hybrids. Yeast strains were stored in cryotubes with YPD (1 % yeast extract, 2 % peptone, and 2 % glucose) and 15 % glycerol at −80 °C. Routine cultures were maintained in YPD plus 2 % agar plates at 24 °C.

### Two new *Hy*brid *Pr*oduction (HyPr) plasmids

We previously published two HyPr plasmids with *natMX* (pHCT2) and *hphMX* (pHMK34) resistance cassettes ^20^. Following our previously described methodology, we amplified the *ble* (ZEOcyn resistance) and *nptII* (G418 resistance) coding regions for marker swaps to generate pHRW32 and pHRW40 plasmids, respectively (Supplementary Table 8). The new HyPr plasmids enabled complex, iterative crossing schemes without adding extra steps to remove one of the two HyPr plasmids between the hybridization steps (Supplementary Fig. 1).

### *Saccharomyces* transformation with HyPr plasmids

Before transforming GLBRCY101 with a HyPr plasmid, we removed its nuclear *kanMX* cassette by swapping the *kanMX* marker to *tkMX* ^42^. Next, we transformed this strain using a short DNA fragment designed to allow the *tkMX* gene to be removed via homologous recombination and selecting for successful marker loss on synthetic complete (SC) + FUdR medium (0.17 % yeast nitrogen base, 0.5 % ammonium sulfate, 0.2 % complete drop out mix, 2 % glucose, and 50 µg/ml 5-fluorodeoxyuridine). *S. cerevisiae* yHWA85 and representative strains of *Saccharomyces paradoxus*, *S. mikatae*, *S. kudriavzevii*, *S. arboricola*, and *S. uvarum* were transformed with one of the four HyPr plasmid versions (Supplementary Table 1). The diploid parent strains contain levels of heterozygosity lower than 0.037 % (Supplementary Table 1).

Transformation of yeast strains was done using the lithium acetate/PEG-4000/carrier DNA method ^43^ with previously described modifications for particular species ^20^. *S. cerevisiae* yHWA85 was first diploidized using the HyPr plasmid pHRW40, creating yHRW134 for subsequent crosses. The generation of this diploid strain occurred in one step, which was confirmed by polymerase chain reaction (PCR) amplification of the *MAT* loci (see below). The experimental reference strain yHRW135 was derived from yHRW134 by screening for spontaneous plasmid loss.

### iHyPr (*i*terative HyPr) method for sequentially generating higher-order hybrids

Following the HyPr method to facilitate mating-type switch ^20^, we pre-cultured strains with differentially marked HyPr plasmids in the presence of doxycycline to express the endonuclease encoded by *HO*, which is under a Tet-ON promoter (Supplementary Fig. 1); each plasmid also contains the full machinery for inducible expression of the promoter. To generate the six-species hybrids yHRWh36 and yHRWh39, we first hybridized three separate pairs of species, generating two-species hybrids (Supplementary Fig. 2A,B). In each case, once the three two-species hybrids were generated, two of those two-species hybrids were themselves hybridized to create a four-species hybrid, which finally was hybridized with the last two-species hybrid to generate the six-species hybrid. To generate the six-species hybrid yHRWh56, two two-species hybrids were separately crossed with diploid *Saccharomyces* strains from other species to create two separate three-species hybrids, which were then mated to generate the six-species hybrid (Supplementary Fig. 2C). Before each cross, parent strains were transformed with differentially marked HyPr plasmids (Supplementary Table 1, 8, Supplementary Fig. 1, 2) and treated with doxycycline in YPD at room temperature, except for *S. cerevisiae* which was incubated at 30 °C. As previously described ^20^, the doxycycline triggers the expression of the Ho endonuclease, which cuts one or more *MAT***a**/*MATα* idiomorphs and generates mating-compatible strains that behave as either *MAT***a** or *MATα*. A sample of each culture was combined in a 1-ml Eppendorf tube and patched on a YPD plate. After 2-3 days, a sample was taken with a toothpick and streaked on a YPD plate supplemented with the corresponding drugs to select for successful matings. In contrast to the original HyPr method, we pre-cultured the new hybrid in YPD with one of the two drugs used during the selective medium step, and that hybrid was then crossed with another strain containing one or two of the other HyPr plasmids not used previously (Supplementary Fig. 1, 2). During these subsequent steps, we expected (and phenotypically verified) the loss of the HyPr plasmid containing the drug-resistance cassette not under selection. This approach and the additional HyPr plasmids made for this study facilitated the iterative crosses required to make six-species hybrids by avoiding the steps of plasmid removal and minimizing the number of generations between crosses (Supplementary Fig. 1).

The frequency of successful two-, three-, four-, and six-species hybrid generation were quantified in duplicates (n=2) (Supplementary Table 3). The patch of co-culture was diluted in sterile H_2_O, and a sample was spread onto both YPD plates and YPD supplemented with the appropriate drugs. The frequency of successful matings was calculated as the ratio between the number of colonies observed in YPD supplemented with the corresponding drugs and the number of colonies observed in YPD.

### Mating-type and PCR-RFLP confirmation of strains

Diploidization of the *S. cerevisiae* strain was confirmed by PCR at the mating-type locus. Hybrid statuses were confirmed by restriction fragment length polymorphism (RFLP) analysis. We used the Standard Taq Polymerase (New England Biolabs, Ipswich, MA) and the primers listed in Supplementary Table 9. Genomic DNA was extracted using the phenol:chloroform method on a strain grown from pre-culture to saturation in YPD. Aliquots of 700 µl of saturated culture were located in 1.5 ml microcentrifuge tubes that contained acid-washed beads. Each tube was centrifuged at maximum speed (15000 rpm) for 5 minutes, and the supernatant was removed. 200 µl of buffer EB (10 mM Tris-Cl, pH 8.0), 200 µl of DNA lysis buffer (10 mM Tris pH 8.0, 1 mM EDTA, 100 mM NaCl, 1 % SDS, 2 % Triton X-100), and 200 µl of phenol:chloroform were added to each tube. Vigorous vortexing was performed for 3-4 minutes, followed by 5 minutes of centrifugation at maximum speed. The top aqueous layer was transferred to 1 ml 100 % EtOH. After an inversion mixture, DNA was precipitated at −80 °C for at least 10-15 minutes. A second centrifugation at maximum speed was performed, and the supernatant was discarded. We washed the pellet with 700 µl of 70 % EtOH, and we centrifuged again to remove any residue or trace of the supernatant. The pellet was dried and resuspended in 100 µl of EB at 50-60 °C for 30 minutes. To remove RNA, we incubated the solution with 0.5 µl of 10 mg/ml RNase A for 30 minutes at 37 °C. DNA was quantified with a Qubit 2.0 Fluorometer (ThermoFisher Scientific).

For PCR-RFLP, resulting PCR products were digested with a restriction enzyme or a combination of multiple restriction enzyme assays able to discriminate among *Saccharomyces* species (New England Biolabs, Ipswich, MA). An extended PCR-RFLP pattern, developed in previous publications ^20, 44^ and this study, are detailed in Supplementary Table 10. Undigested PCR products were visualized on a 1.5 % agarose gel, while digested PCR products were visualized on a 3 % agarose gel.

### Ploidy estimation by flow cytometry

Both asynchronous and hydroxyurea-arrested (G1/S arrested) mid-log cultures were prepared for each strain. Hydroxyurea-arrested strains were prepared to assist in the identification of G1 peaks in samples with broad and undefined cell cycle peaks. Briefly, cultures were grown to saturation and then diluted back 1:200. Back-diluted cultures were grown to a 0.4-0.6 optical density at 600 nm (OD_600_). For each strain, 1 ml mid-log culture was transferred into 200 μl 1 M hydroxyurea and incubated on a room temperature culture wheel for approximately half of the time that the respective strain took to grow from back-dilution to an OD_600_ of 0.4-0.6. This ranged between 3 and 12 hours. At the same time, 1 ml of asynchronous mid-log culture was harvested for fixation. All samples were fixed in 70 % ethanol overnight, treated with RNase and Proteinase K, and finally stained with SYTOX Green dye (Molecular Probes) ^45^. Stained cell suspensions were sonicated before flow cytometry. Fluorescence was measured with a BL1 laser (488 nm) on an Attune NxT (Invitrogen) flow cytometer at the lowest available flow rate. To accommodate for extremes in ploidy and cell size, voltage was adjusted to 250 for FITC dump channel (BL1) and FSC (Forward SCatter). All samples were run at the same voltage.

Flow cytometry data files were processed in `FlowJo v10.4.2` ^46^. Samples were first gated on SSC (Side SCatter) and FSC to remove debris. Doublets were then removed by gating on BL1-A and FSC-A. A histogram of BL1-A values were then generated for remaining cells. Hydroxyurea peaks were identified and gated manually. Asynchronous G1 and G2 peaks were identified by applying a Watson (Pragmatic) Cell Cycle model and identifying G1 and G2 means. When cell cycle models did not fit the asynchronous sample data automatically, hydroxyurea samples were used to identify G1 peaks and these were manually gated to constrain G1 in asynchrounous samples. Ploidy estimation were performed by comparing with fluorescence values of a haploid laboratory reference *S. cerevisiae* strain, S288C (Supplementary Table 1).

### Cell size estimation

A subset of strains were used for microscopy analysis of cell size. Each strain was spotted from frozen stock onto YPD agar plates and grown at room temperature for 4 days. Water-cell suspensions were prepared for each strain, which were bright-field-imaged on an EVOS FL Auto 2.0 (Invitrogen) imaging system at 400x. Cell area was analyzed in `FIJI v2.0.0-rc-34/1.5a` ^47^ using the Analyze Particles tool.

### Fitness quantification of the newly generated hybrids

To measure the impact of genome size increases on fitness, we performed a growth test in a rich medium. All parent species and the two-, three-, four-, and six-species hybrids were pre-cultured in 3 ml of YPD at room temperature. After pre-culture, 10 µl of saturated culture was inoculated into a 96-well plate (Nunc, Roskilde, Denmark) containing 240 µl of YPD. Spaces between the wells in the plates were filled with sterile H_2_O to maintain the humidity of the plates and limit culture evaporation.

To monitor the growth of strains and populations in the different media conditions, the inoculated 96-well plate was placed in a BMG FLUOstar Omega (Ortenberg, Germany) at 20 °C. Absorbance at 595 nm was monitored every 15 min for 4 days. Background absorbance was subtracted from the average of nine negative controls containing the uninoculated medium being tested. Kinetic parameters for each condition were calculated in `GCAT v6.3` ^48^. Median and standard deviations from six independent biological replicates, except yHRWh36 and yHRWh56 from which we obtained three replicates, were calculated in `R` ^49^ (Supplementary Table 4).

### Genome sequencing and chromosome composition analyses

Genomic DNA (gDNA) samples from the diploidized *S. cerevisiae* strain and the two-, three-, four-, and six-species hybrids were submitted to the DOE Joint Genome Institute for paired-end Illumina sequencing. Evolved six-species hybrids and six individual colonies from yHRWh88 (see ALE section) were also submitted for sequencing. Libraries were constructed according to the manufacturer’s instructions. Sequencing of the flow cell was performed on an Illumina MiSeq using MiSeq Reagent kits, following a 2×150 nucleotide, indexed run recipe. Curated raw reads were submitted to the SRA database as Bioproject PRJNA476226 (Supplementary Table 11).

Genomic characterization was performed with `sppIDer v1` ^50^. Our combined nuclear reference genome was built with the genome assemblies of *S. cerevisiae* GLBRCY22-3 ^51^, which is a close relative of the biofuel reference strain used here; *S. paradoxus* CBS432; *S. arboricola* CBS10644 ^52^; *S. mikatae* IFO1815; *S. kudriavzevii* ZP591; *S. uvarum* CBS7001 ^18^; and *Saccharomyces eubayanus* FM1318 ^53^. Our combined mitochondrial reference genome was built with the mitochondrial assemblies of the aforementioned strains ^52–54^, except for CBS7001, whose mtDNA is still not completely assembled ^39^. Instead, we used the mtDNA of a close relative, *S. uvarum* CBS395 ^54^. Raw Illumina paired-end reads and the combined reference genomes were the input data of `sppIDer`, which is a wrapper that runs published tools to map the short reads to the combined reference genomes and creates several colorful and visually intuitive outputs ^50^. Here, we show depth of coverage plots from those species contributing genomes.

For each strain, the number of chromosomes and the ploidy were estimated from the `sppIDer` plots. This approximation gave a significant positive correlation with the ploidy estimated by flow cytometry (Spearman rank test r = 0.91, *p*-value = 3.2*10^-6^) (Supplementary Fig. 5C). The number of chromosomal aberrations was based on the number of gains, losses, or unbalanced translocations detected in the `sppIDer` plots (Supplementary Table 2). One chromosomal gain, loss, or unbalanced translocation was counted as one aberration. Aberrations observed in one hybrid and maintained in the offspring of subsequent crosses were not counted again; only new aberrations for each cross were reported in the aberration plot (Figure 2A, 6A). Chromosomal aberrations involving parts of chromosomes were conservatively counted only in cases where there were clear fusions of entire chromosome arms.

### Genome size and ploidy quantification from short-read sequences

Two different approaches were performed to quantify the genome size of the sequenced strains. In the first approach, genome assemblies were performed using the collection of assemblers included in `iWGS v1.1` ^55^. The assembly with the best assembly stats reported by `iWGS` was selected, and the genome size was reported (Supplementary Table 2). In the second approach, `sppIDer` coverage outputs (StrainName_winAvgDepth-d.txt) were parsed to quantify the percentage of each *Saccharomyces* nuclear genome retained in the hybrid, which was calculated as follows:

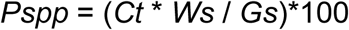

where *Pspp*, is the percentage for one of the parent species; *Ct*, is the number of windows with a coverage mean value above 2; *Ws*, is the window size; and *Gs* is the reference genome size for that parent species. These two calculations yielded a good approximation of the increased genome size, but both generated estimates that assumed the highly homozygous genome donated by each parent was haploid; `iWGS` and `sppIDer` plots were significantly correlated (Spearman rank test r = 0.95, *p*-value = 2.2*10^-16^, Supplementary Fig. 5D).

To get a better approximation of the genome size of each allopolyploid, we first determined the total number of copies of each chromosome contributed by each species, as quantified by `sppIDer`. Genome size was then calculated by multiplying the number of copies of each chromosome by its length and adding all these values together. Genome size and flow cytometry fluorescence were correlated (Spearman rank test r = 0.93, *p-*value = 1.1*10^-7^, Supplementary Fig. 5B).

### Quantification of the number of copies of the xylose utilization cassette

Illumina reads were extracted using the xylose utilization cassette sequence (8.7 Kbp) as bait for `HybPiper v1.2` ^56^. The generated bam files were viewed and sorted with `samtools v1.4` ^57^, and the coverage for each nucleotide was quantified with `genomeCoverageBed`, which is included in `bedtools v2.2.27` ^58^. The mean coverage of the coding sequence of the three engineered xylose utilization genes (*XYL1*, *XYL2*, and *XYL3*) (3.9 Kbp), was calculated from the `genomeCoverageBed` output. For the chromosome IV, mean coverage values for windows of 3.9 Kbp were calculated from the `genomeCoverageBed` output generated by `sppIDer`. The cassette value and chromosome distributions for each strain were compared by a one-side Wilcoxon rank sum test for a significant deviation from the expected ratio 1:1 (1 copy of the cassette to one copy of chromosome IV) (Supplementary Table 7).

### Adaptive laboratory evolution (ALE) and colony selection

Two of the three six-species hybrids (during construction, the third lost *S. cerevisiae* chromosome IV, where *Sch. stipitis* xylose utilization genes had been inserted) were evolved in triplicate at room temperature in tubes with two independent media conditions: 3.0 ml YPD or 3.0 ml YPX (1 % yeast extract, 2 % peptone, and 2 % xylose). Three to five days of fermentation were performed to allow cells to consume the sugars, and an aliquot of each replicate was transferred at of 0.1 OD_600_ to a fresh medium until it reached approximately 80 generations. A colony from each independent ALE experiment, regardless of whether they were evolved in glucose or xylose, was selected on YPX plates (1 % yeast extract, 2 % peptone, 2 % xylose, and 2 % agar) and cryopreserved.

### Microtiter plate growth curves

We compared the growth kinetics of the *S. cerevisiae* reference strain yHRW135, the ancestors of the two six-species hybrids retaining the chromosome IV (yHRWh39, yHRWh56), and populations of the evolved hybrids. Growth was tested in YPD and YPX at room temperature. Strains or populations were pre-cultured in 3.0 ml YPD or YPX, depending of the medium tested. After pre-culture, 10 µl of saturated culture was inoculated into a 96-well plate (Nunc, Roskilde, Denmark) containing 240 µl of identical medium as the pre-culture. Spaces between the wells in the plates were filled with sterile H_2_O to maintain the humidity of the plates. The reference strain was cross-inoculated in all conditions; for example, yHRW135 pre-cultured in YPX was tested in both YPD and in YPX.

To monitor the growth of strains and populations in the different media, we inoculated 96-well plates and placed them in a BMG FLUOstar Omega at 20 °C. Absorbance at 595 nm was monitored every 15 min for 5 days. Background absorbance was subtracted from the average of three negative controls containing the uninoculated medium being tested. Kinetic parameters for each condition were calculated in `GCAT v6.3` ^48^. Median and standard deviations from three independent biological replicates were calculated in `R` ^49^ (Supplementary Table 6). For each medium, parameters were normalized against the data generated by the reference strain yHRW135 when it was pre-cultured and grown in the medium tested.

### Cold tolerance spot test

Temperature growth profiles are well known to vary among *Saccharomyces* species ^27, 28^. In particular, *S. uvarum* and *S. kudriavzevii* are able to grow at low temperatures where *S. cerevisiae* cannot grow. To test if some phenotypic traits might be retained independently of the media regime, we performed spot tests in rich medium at different temperatures (22 °C, 10 °C, and 4 °C). The *S. cerevisiae* reference strain (yHRW135) and the evolved six-species hybrids were compared. All strains were pre-cultured in liquid YPD medium at room temperature to saturation. Cultures were subjected to a series of 10-fold dilutions in YPD. 5 µL of each dilution was spotted onto three YPD-agar plates, identically. Plates were incubated in sealed plastic bags to keep them from drying out at the temperatures mentioned above. Each plate was photographed when most strains exhibited significant growth (4 days for 22°C, 11 days for 10°C, and 38 days for 4°C).

### Culture wheel growth curves

Strains isolated from single colonies from evolved hybrids, ancestor hybrids, and the reference strain (yHRW135) were pre-cultured in YPX and inoculated at an initial OD_600_ of 0.1 into 3 ml glass tubes containing YPX. Growth was monitored by measuring OD_600_. Kinetic parameters were calculated as above. Median and standard deviations from six independent biological replicates were calculated as above. These experimental conditions most closely matched the conditions in which the strains were evolved, and they are reported in Figure 5A and Supplementary Table 5.

### Statistical analyses

Data analyses and plots were performed in `R` ^49^. Linear models of regressions were added to the plots in Figure 3, 4A, Supplementary Fig. 5, 8 using the `geom_smooth` option in the R package `ggplot2`. A LOESS regression line was added to the plot in Figure 2A using the `geom_smooth` option in the `R` package `ggplot2.` For aberration data (Figure 2A), r^2^ and significance of regression were calculated with `summary(lm(y ∼ x)),` where x was the number of species, and y was the number of observed aberrations. Correlations for ploidy and assembly comparisons were calculated in `R` using the `ggpubr` package to apply a Spearman rank sum test (Figure 3, Supplementary Fig. 5, 8), and plots were generated using `ggplot2`.

The impact of mitochondrial inheritance (Figure 4B) in the retention of the nuclear genome of those hybrids involving *S. cerevisiae* was tested using a multifactor ANOVA in `R`, using `summary(aov(P ∼ M * C))`, where *P* is the percentage retained of the *S. cerevisiae* nuclear genome; *M* is the mtDNA, which was encoded as a binary character (either as the *S. cerevisiae* mtDNA or that of another species); and *C* is the type of strain (i.e. classified as the *S. cerevisiae* parent; two-, three-, four-, or ancestor six-species hybrid; and evolved six-species hybrid).

t-tests for significant differences between frequency of chromosome gains and losses and Wilcoxon rank sum tests for significant differences in the kinetic parameters shown in Figure 3D, 5A and Supplementary Fig. 6, respectively, were performed in `R`.

Flow cytometry data were analyzed and plotted in `R`. Correlations were tested in `R` using a Spearman rank sum test and plotted using `ggplot2`.

## Acknowledgments

We thank Trey K. Sato for providing GLBRCY101, Srivatsan Raman for flow cytometry access, Amanda B. Hulfachor for assistance with Figure 5B, the Joint Genome Institute (JGI) for providing Illumina Sequencing services, and Miguel Morard for feedback on preliminary figures. This material is based upon work supported in part by the Great Lakes Bioenergy Research Center, U.S. Department of Energy, Office of Science, Office of Biological and Environmental Research under Award Numbers DE-SC0018409 and DE-FC02-07ER64494; the National Science Foundation under Grant Number DEB-1253634; the USDA National Institute of Food and Agriculture Hatch Project Number 1020204, and the Robert Draper Technology Innovation Fund from the Wisconsin Alumni Research Foundation (WARF). DP is a Marie Sklodowska-Curie fellow of the European Union’s Horizon 2020 research and innovation programme, grant agreement No. 747775. KJF is a Morgridge Metabolism Interdisciplinary Fellow of the Morgridge Institute for Research. CTH is a Pew Scholar in the Biomedical Sciences, a Vilas Early Career Investigator, and a H. I. Romnes Faculty Fellow, supported by the Pew Charitable Trusts, Vilas Trust Estate, and Office of the Vice Chancellor for Research and Graduate Education with funding from WARF, respectively. The work conducted by the U.S. Department of Energy Joint Genome Institute, a DOE Office of Science User Facility, is supported under Contract No. DE-AC02-05CH11231.

## Authors’ contributions

Conceived the experiments: DP, WGA, KF, RLW, and CTH. DP and CTH mentored RVM, and WGA mentored MGB and EJU. Engineered strains: WGA, RVM, and RLW. Data generation: WGA, KF, MGB, EJU, and RLW. Data analysis: DP and KF. Supervised the study: DP and CTH. Wrote the paper with editorial input from all co-authors: DP and CTH.

## Additional information

Supplementary Information:

Supplementary Information

Supplementary Figures 1-9.

Supplementary Tables 1-11.

## Data availability

Raw genome sequencing data has been deposited in NCBI’s SRA database, Bioproject PRJNA476226. HyPr plasmids are being deposited in Addgene as deposit 77444.

## Materials & Correspondence

Requests for materials should be addressed to. Correspondence should be addressed to.

## Supplementary Figure Legends

**Supplementary Figure 1 | The iHyPr method enabled the formation of higher-order synthetic hybrids using iterative crosses.** A simplified scheme comparing the protocol to generate an allohexaploid (6n) synthetic hybrid using iHyPr is displayed, in contrast with HyPr, which is not iterative. NAT, Nourseothricin; HYG, hygromycin; ZEO, zeocin. *MAT* idiomorphs examples are shortened to **a** and *α*.

**Supplementary Figure 2 | Schematics for the generation of three six-species *Saccharomyces* hybrids.** The hybridization steps necessary to generate the six-species hybrids yHRWh36, yHRWh39, and yHRWh56 are represented in panels A), B) and C), respectively. Yeast cells are represented in gray, and chromosomes are colored according to the *Saccharomyces* species. The strain names of our lab’s copy of some strains (Supplementary Table 1) are displayed in parentheses below the original culture collection strains. Drug resistance is indicated above yeast cells according to the abbreviations in Supplementary Figure 1. Systematic crosses are highlighted with arrows to form a pedigree. The black lightning bolt symbol represents the doxycycline shock to promote mating type switching or loss to facilitate hybridization.

**Supplementary Figure 3 | Nuclear genome composition of the diploidized *S. cerevisiae* reference strain and the synthetic and evolved hybrids.** Panels A-Z,AB are the `sppIDer` outputs for the diploidized reference strain of *S. cerevisiae* (GLBRCY101) and the synthetic and evolved hybrids. Sequencing coverage values are colored according to each *Saccharomyces* species’ contribution in that portion of the genome. Panels were ordered to represent synthetic hybrid data based on the order they were used to generate the next hybrid (Supplementary Fig. 2). `sppIDer` produces multiple plots ^50^, but here we show the log_2_ of the average coverage of ∼8 Kbp-windows normalized to the genome-wide average coverage. To improve the resolution of the three-, four-, and six-species hybrid plots, window coverage values were normalized to the genome-wide average coverage, and values were limited to the 99% percentile and below.

**Supplementary Figure 4 | Mitochondrial genome inheritance of the diploidized *S. cerevisiae* reference strain and the synthetic and evolved hybrids.** Panels A-C are the `sppIDer` outputs for the diploidized reference strain of *S. cerevisiae* (GLBRCY101) and the synthetic and evolved hybrids. Sequencing coverage values are colored according to each *Saccharomyces* species’ contribution in that portion of the mtDNA. Each panel contains the mtDNA inheritance for the synthetic hybrid used to generate that particular six-species hybrids (Supplementary Fig. 2). `sppIDer` produces multiple plots ^50^, but here we show the log_2_ of the average coverage of 44-bp windows normalized to the mtDNA-wide average coverage. When a synthetic hybrid is formed between parent strains that both contain mtDNA, a heteroplasmic state can be maintained for several generations, but eventually, a parent or recombinant mtDNA is generally fixed ^23^. In some hybrids, this heteroplasmic state persisted, and the names of hybrids are colored in a gradient according to the detected mtDNAs; these colors are also displayed in Figure 4A. Due to the unusually high coverage of *ATP9* or *ATP9*-*VAR1-15S rRNA* of *S. uvarum* in panel A), additional inset plots with limited y-axes are shown.

**Supplementary Figure 5 | Ploidy and genome size estimations were well-correlated among different methods.** A) Genome size (Supplementary Table 2) was correlated with the ploidy estimates from flow cytometry. B) Genome size was correlated with the mean fluorescence (n=10000, counts per strain) of SYTOX Green. C) Ploidy estimated from `iWGS` (Supplementary Table 2) was correlated with the ploidy estimated using flow cytometry. D) The estimates of the amount of unique DNA present were correlated between `iWGS` and `sppIDer`. `sppIDer` values were corrected for copy number to generate the genome size estimates in the other panels. Points are colored according to the number of species (nuclear complexity) genomes contributing to the strain. Synthetic hybrids generated from each independent scheme are represented with different shapes. The Spearman rank sum test R and *p*-value are displayed. Linear regression lines and their 95% confidence intervals of the fit are represented with a black line and gray shadow, respectively.

**Supplementary Figure 6 | Growth kinetics for ancestor and evolved six-species populations in a microtiter plate.** Growth measured as area under the curve (AUC) for the reference *S. cerevisiae* strain and evolved six-species hybrids, following normalization to the *S. cerevisiae* reference strain yHRW135 (full microtiter plate kinetic parameters are reported in Supplementary Table 4). Different shapes indicate the media in which the synthetic six-species hybrids were evolved. Colors differentiate the reference strain (orange), and the ancestor of evolved hybrids (red for hybrids evolved from yHRWh39 and purple for hybrids evolved from yHRWh56), while each data point represents an evolved replicate population.

**Supplementary Figure 7 | The fitness improvement during adaptive laboratory evolution was not due to an increase in the number of copies of xylose utilization genes.** A) Schematic representation of the metabolic pathway for xylose utilization. The engineered xylose utilization genes are highlighted in blue. B) Boxplots of coverage levels for 3.9 Kbp windows of chromosome IV are displayed for each strain. Median values for the strains are represented by a horizontal line inside the box, and the upper and lower whiskers represent the highest and lowest values of the 1.5 * IQR (inter-quartile range), respectively. Color dots show the coverage values for the coding sequences of the engineered xylose utilization genes. Points are colored according to the number of species (nuclear complexity) contributing to the strain. The coverage values of the xylose utilization genes were not significantly higher than the values for the chromosome IV (Supplementary Table 7).

**Supplementary Figure 8 | The fitness improvement during adaptive laboratory evolution was not simply due to genome reduction**. **Genome size (Supplementary** Table 2) was not significantly correlated with the maximum specific growth rate (µ, defined as (ln(OD2)-ln(OD1))/(T2-T1))) (Supplementary Table 5). Ancestor (purple dots) and evolved six-species strains (brown dots) are shown. Different shapes indicate the media in which the synthetic six-species hybrids were evolved. Color points differentiate the ancestor from the evolved hybrids.

