## Supplementary Figures 1-8 for "Synthetic hybrids of six yeast species"

### Supplementary figure 1

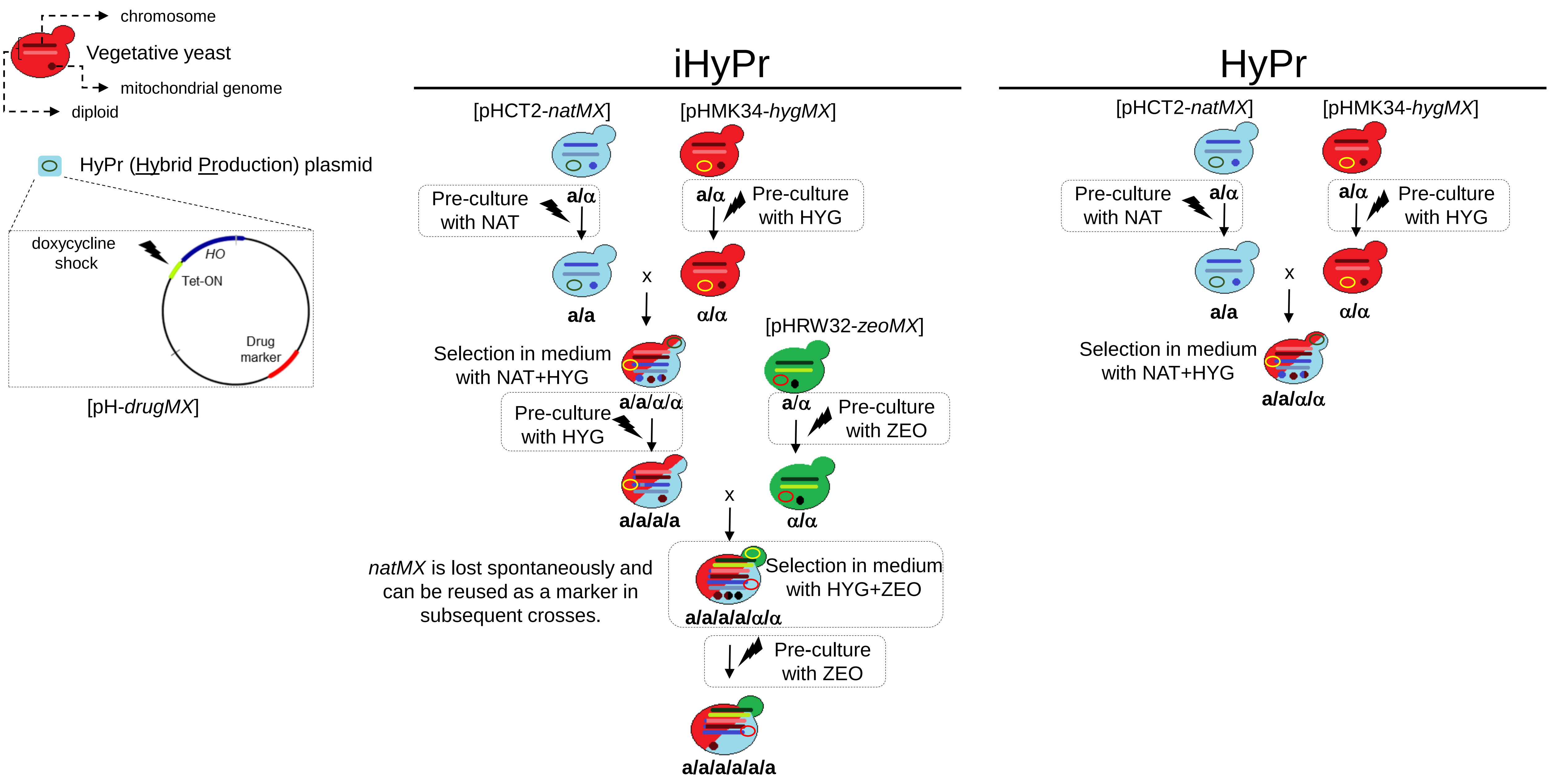

**Supplementary figure 2**

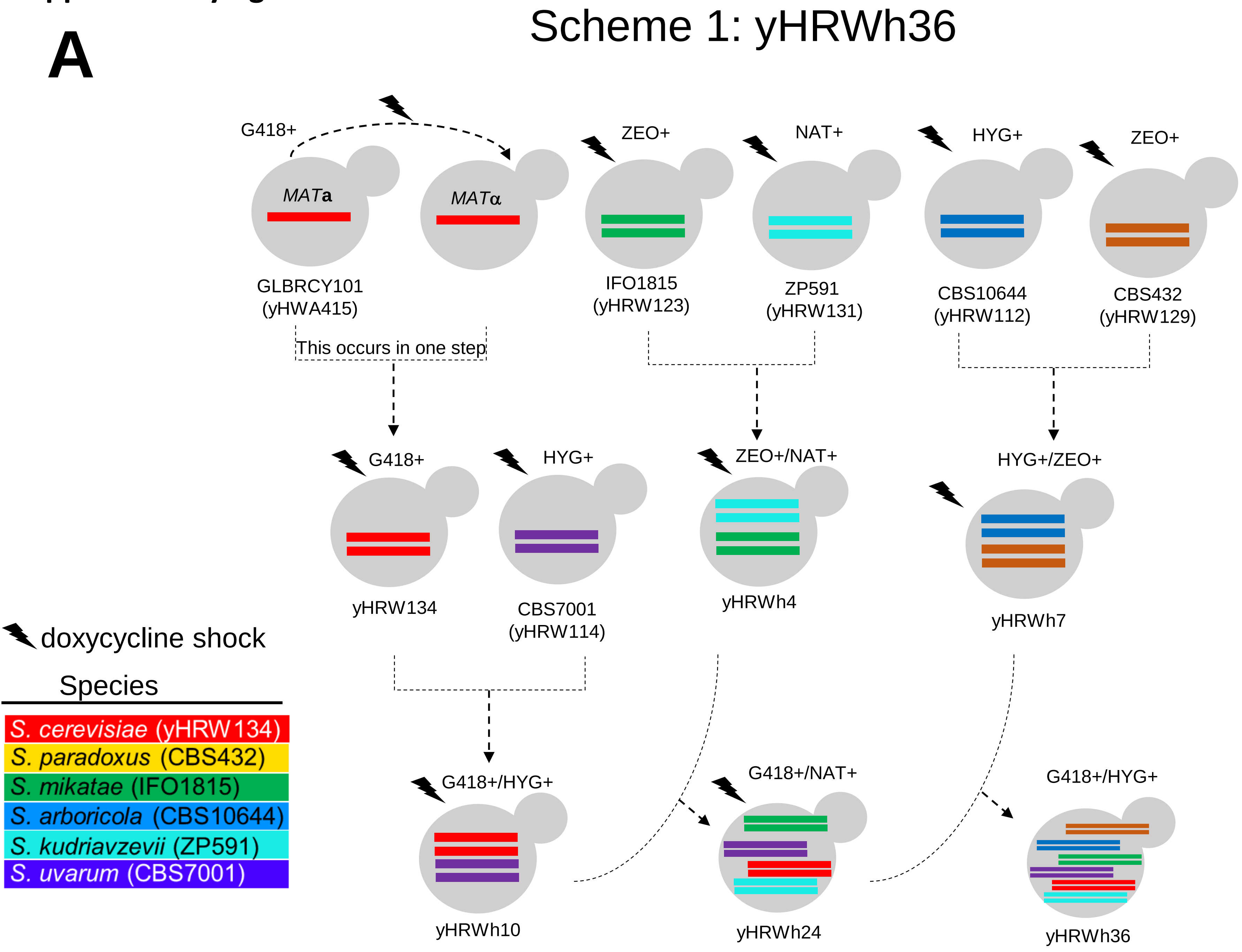

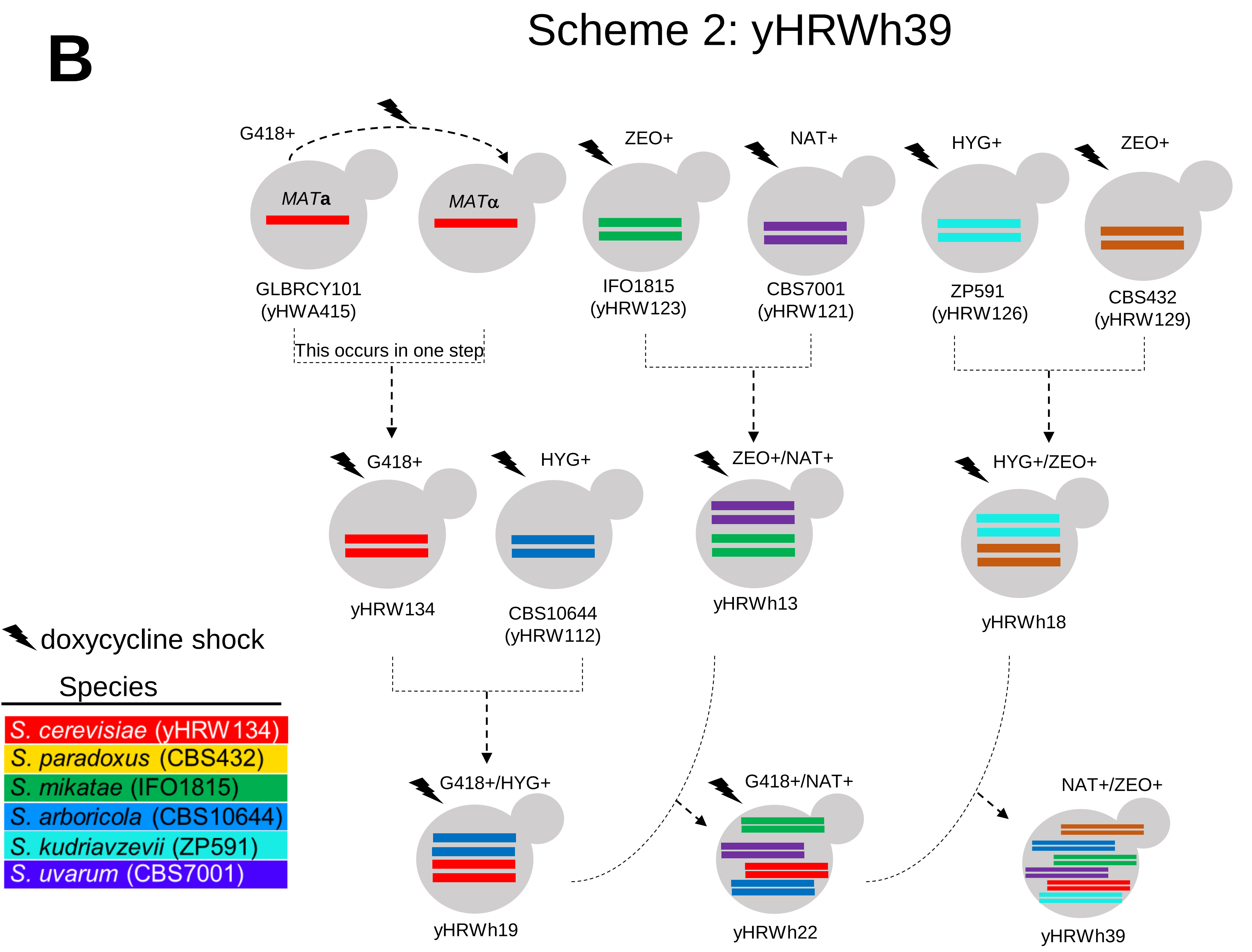
B
Scheme 2: yHRWh39
G418+
MATa
MATα
GLBRCY101
(yHWA415)
ZEO+
IFO1815
(yHRW123)
NAT+
CBS7001
(yHRW121)
HYG+
ZP591
(yHRW126)
ZEO+
CBS432
(yHRW129)
This occurs in one step
G418+
yHRW134
HYG+
CBS10644
(yHRW112)
ZEO+/NAT+
yHRWh13
HYG+/ZEO+
yHRWh18
doxycycline shock
Species
S. cerevisiae (yHRW134)
S. paradoxus (CBS432)
S. mikatae (IFO1815)
S. arboricola (CBS10644)
S. kudriavzevii (ZP591)
S. uvarum (CBS7001)
G418+/HYG+
yHRWh19
G418+/NAT+
yHRWh22
NAT+/ZEO+
yHRWh39

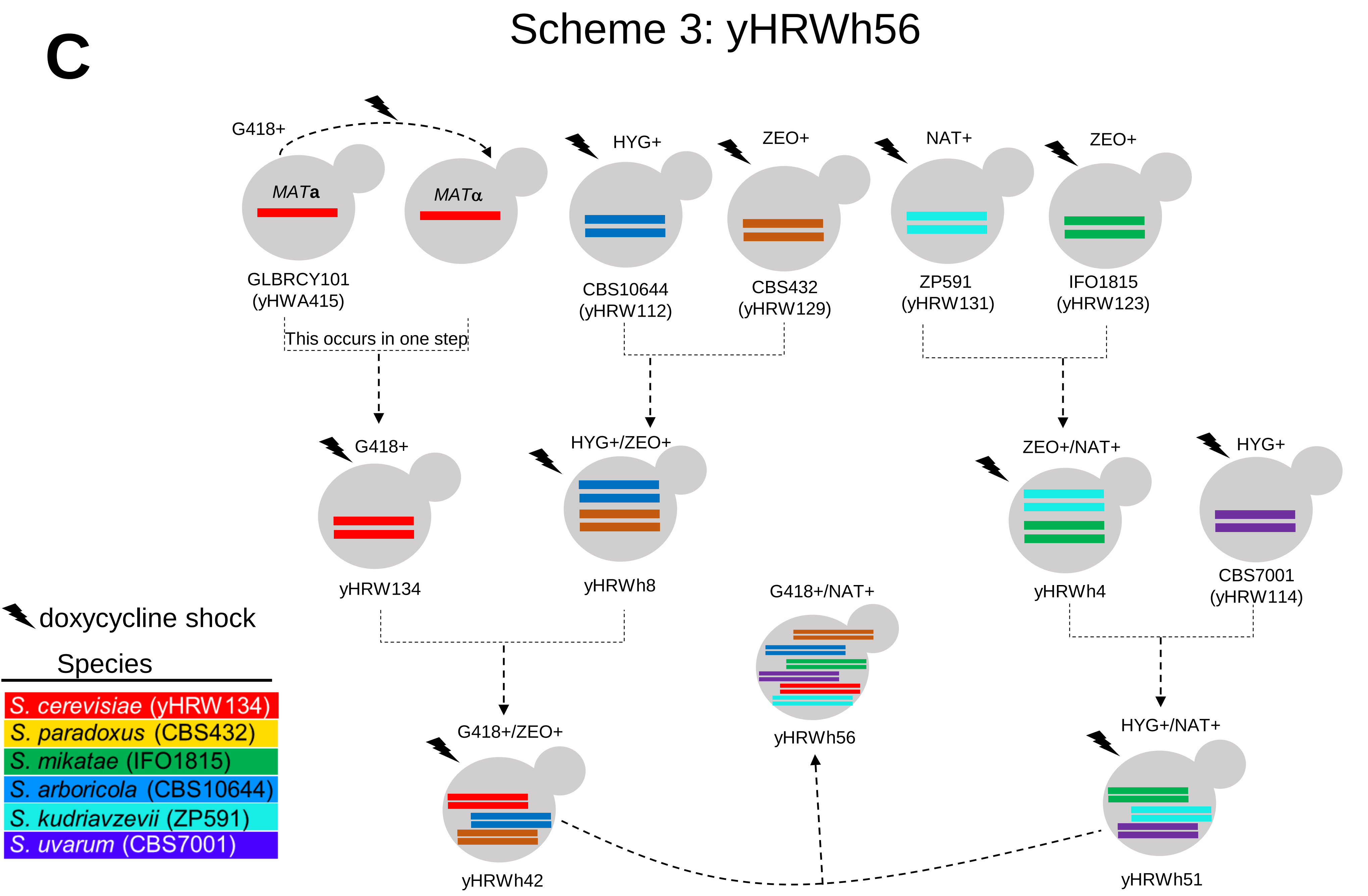
C
Scheme 3: yHRWh56
G418+
MATa
MATα
GLBRCY101
(yHWA415)
HYG+
CBS10644
(yHRW112)
ZEO+
CBS432
(yHRW129)
NAT+
ZP591
(yHRW131)
ZEO+
IFO1815
(yHRW123)
This occurs in one step
G418+
yHRW134
HYG+/ZEO+
yHRWh8
ZEO+/NAT+
yHRWh4
HYG+
CBS7001
(yHRW114)
G418+/NAT+
yHRWh56
G418+/ZEO+
yHRWh42
HYG+/NAT+
yHRWh51
doxycycline shock
Species
S. cerevisiae (yHRW134)
S. paradoxus (CBS432)
S. mikatae (IFO1815)
S. arboricola (CBS10644)
S. kudriavzevii (ZP591)
S. uvarum (CBS7001)

**Supplementary figure 3**

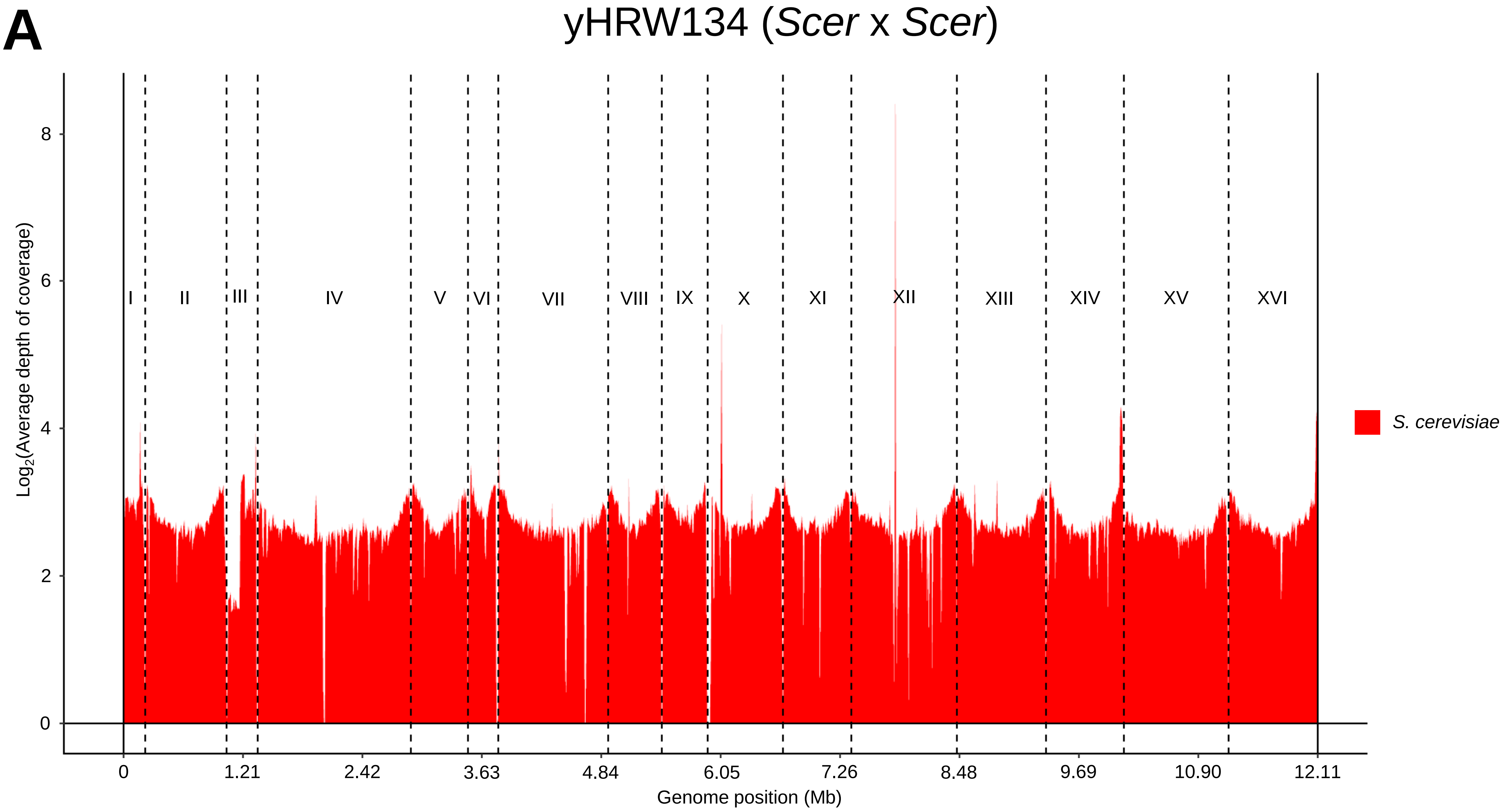

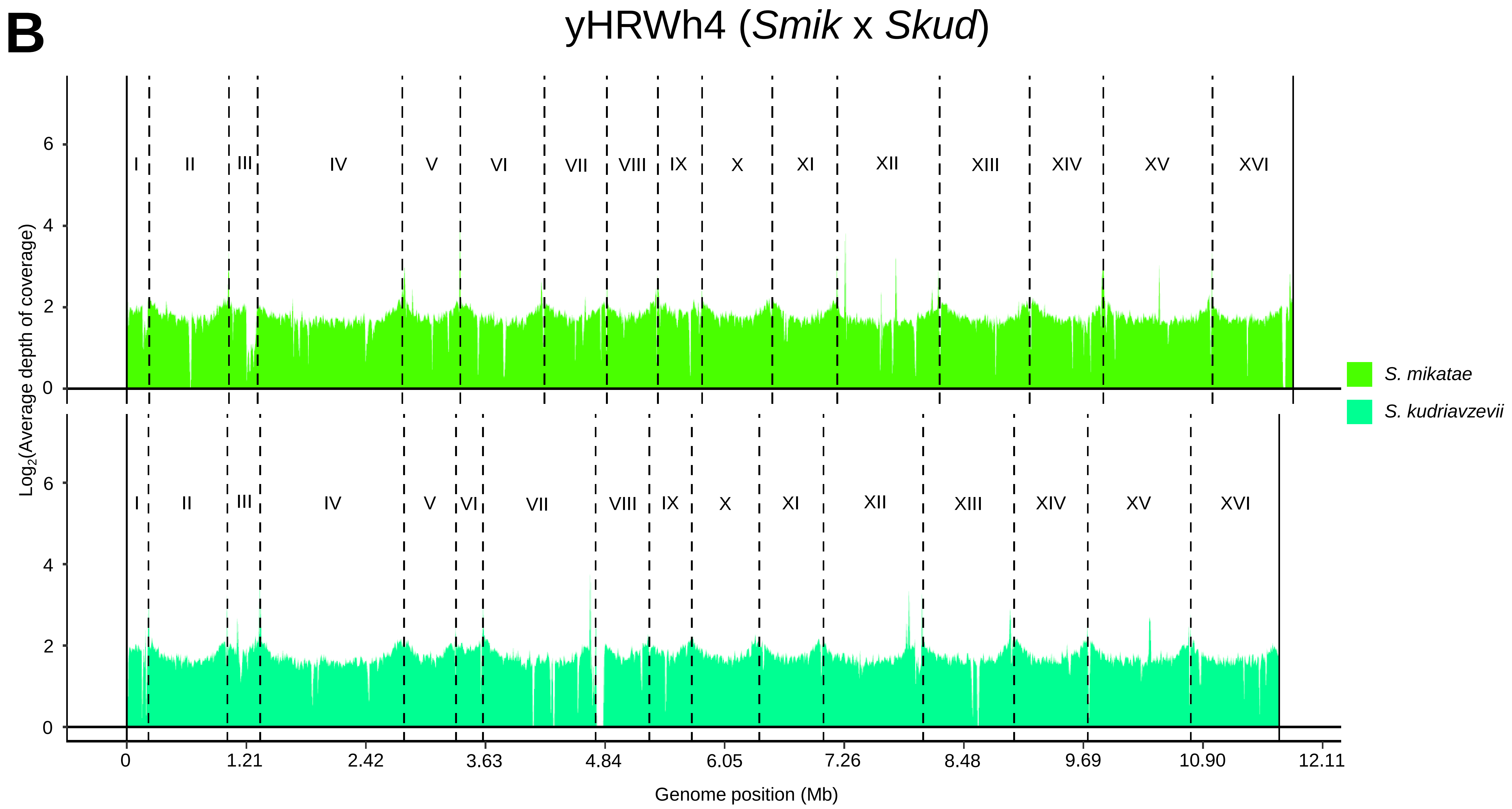

B
yHRWh4 (*Smik* x *Skud*)
I
II
III
IV
V
VI
VII
VIII
IX
X
XI
XII
XIII
XIV
XV
XVI
0
2
4
6
Log2(Average depth of coverage)
Genome position (Mb)
0
1.21
2.42
3.63
4.84
6.05
7.26
8.48
9.69
10.90
12.11
*S. mikatae*
*S. kudriavzevii*

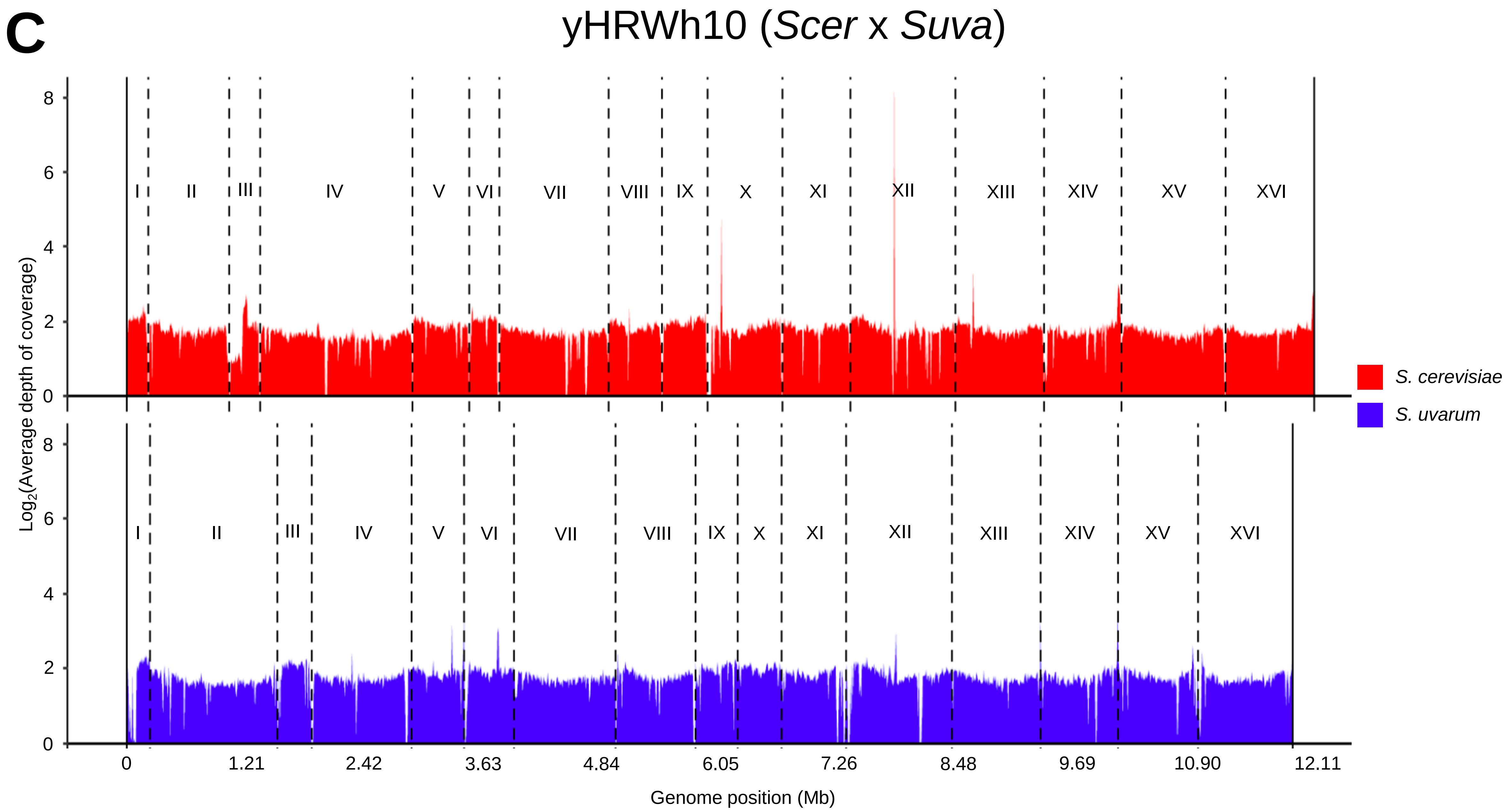
C
yHRWh10 (*Scer* x *Suva*)
I
II
III
IV
V
VI
VII
VIII
IX
X
XI
XII
XIII
XIV
XV
XVI
Log2(Average depth of coverage)
0
2
4
6
8
0
1.21
2.42
3.63
4.84
6.05
7.26
8.48
9.69
10.90
12.11
Genome position (Mb)
S. cerevisiae
S. uvarum

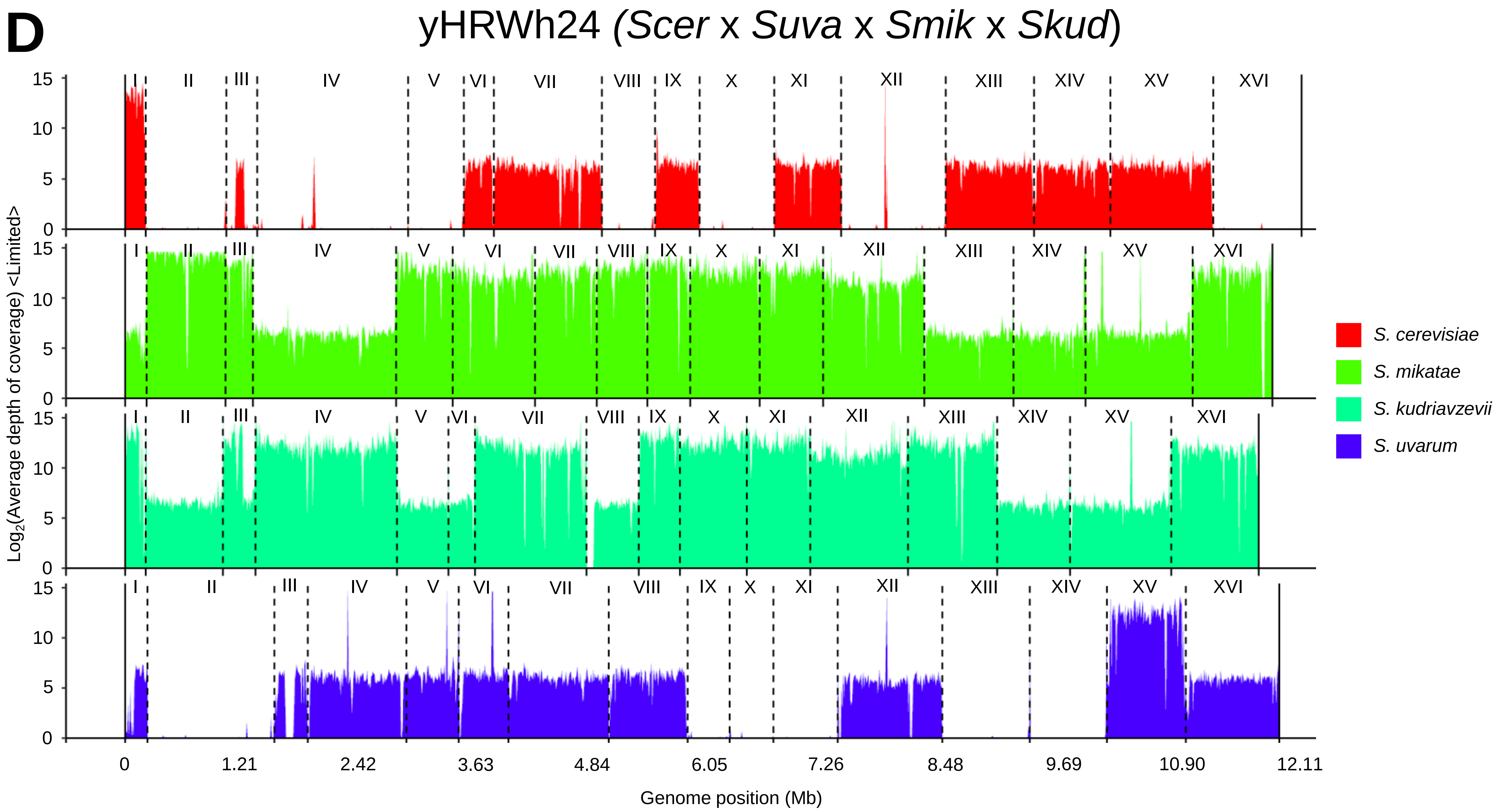
D
yHRWh24 (*Scer* x *Suva* x *Smik* x *Skud*)
Log2(Average depth of coverage) <Limited>
I
II
III
IV
V
VI
VII
VIII
IX
X
XI
XII
XIII
XIV
XV
XVI
0
5
10
15
0
1.21
2.42
3.63
4.84
6.05
7.26
8.48
9.69
10.90
12.11
Genome position (Mb)
S. cerevisiae
S. mikatae
S. kudriavzevii
S. uvarum

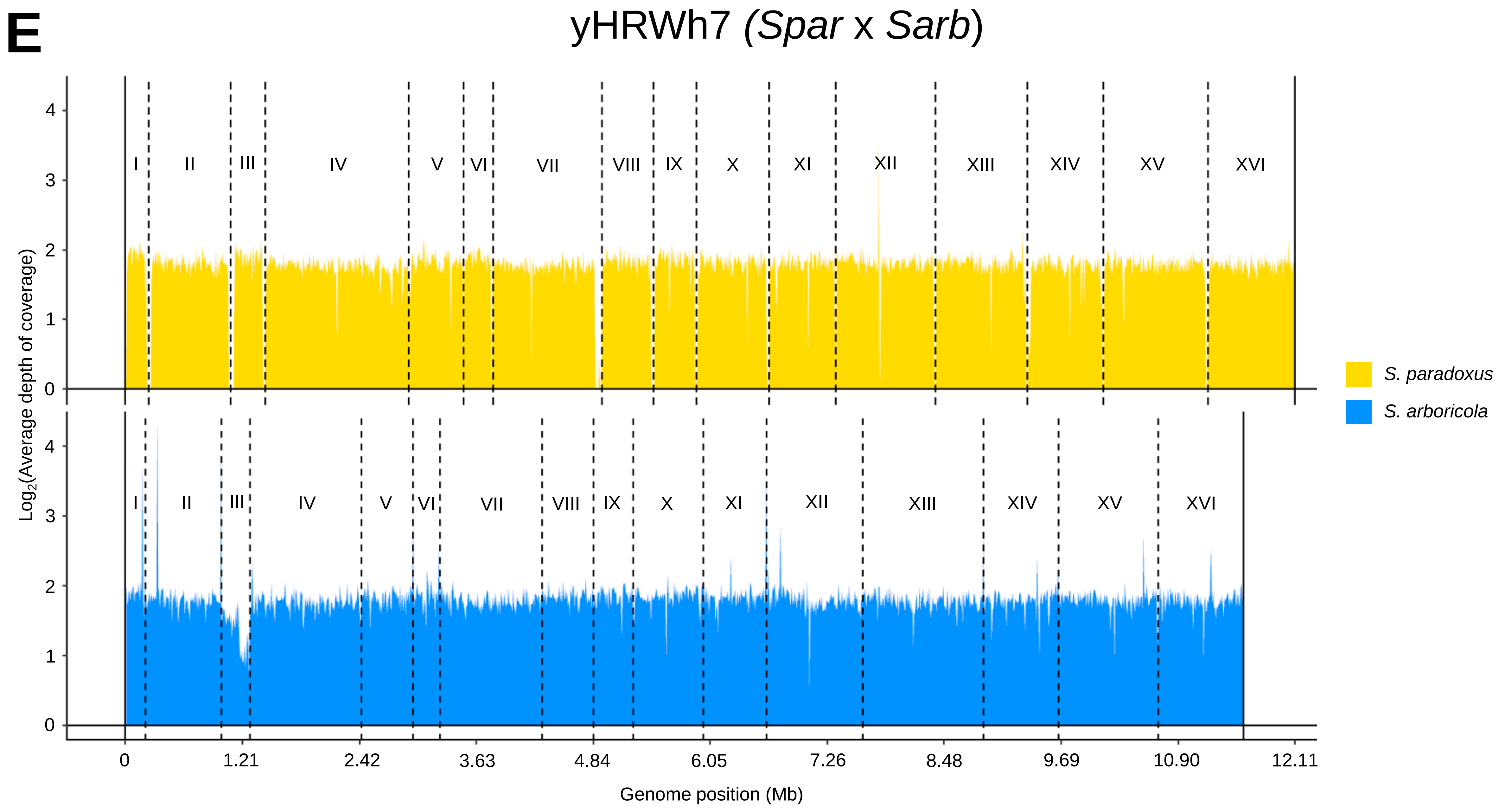
E
yHRWh7 (*Spar* x *Sarb*)
Log2(Average depth of coverage)
0
1
2
3
4
I
II
III
IV
V
VI
VII
VIII
IX
X
XI
XII
XIII
XIV
XV
XVI
0
1.21
2.42
3.63
4.84
6.05
7.26
8.48
9.69
10.90
12.11
Genome position (Mb)
*S. paradoxus*
*S. arboricola*

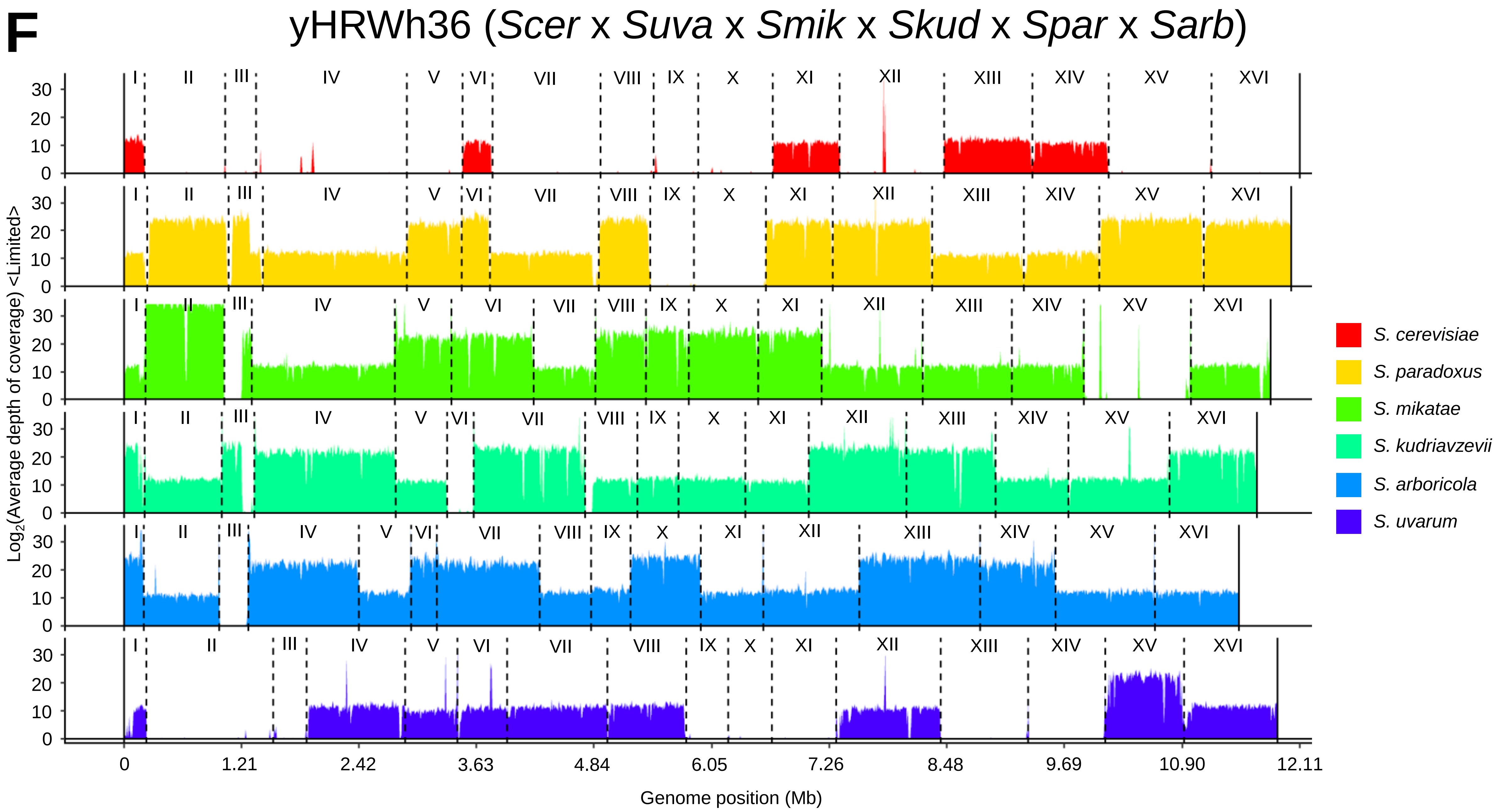
F
yHRWh36 (*Scer* x *Suva* x *Smik* x *Skud* x *Spar* x *Sarb*)
I II III IV V VI VII VIII IX X XI XII XIII XIV XV XVI
0 10 20 30
Log2(Average depth of coverage) <Limited>
*S. cerevisiae*
*S. paradoxus*
*S. mikatae*
*S. kudriavzevii*
*S. arboricola*
*S. uvarum*
0 1.21 2.42 3.63 4.84 6.05 7.26 8.48 9.69 10.90 12.11
Genome position (Mb)

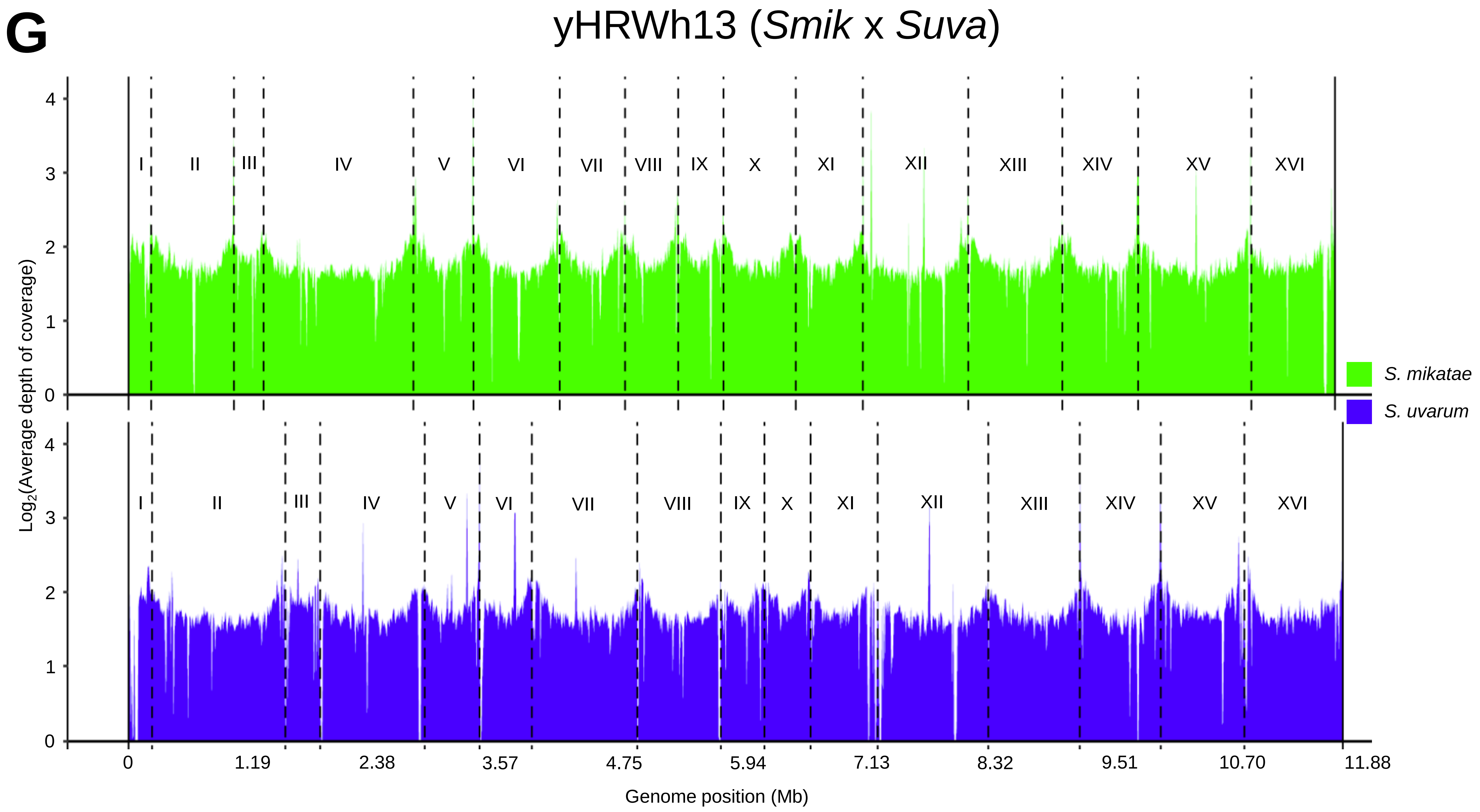

G
yHRWh13 (*Smik* x *Suva*)
I
II
III
IV
V
VI
VII
VIII
IX
X
XI
XII
XIII
XIV
XV
XVI
Log2(Average depth of coverage)
0
1
2
3
4
0
1.19
2.38
3.57
4.75
5.94
7.13
8.32
9.51
10.70
11.88
Genome position (Mb)
S. mikatae
S. uvarum

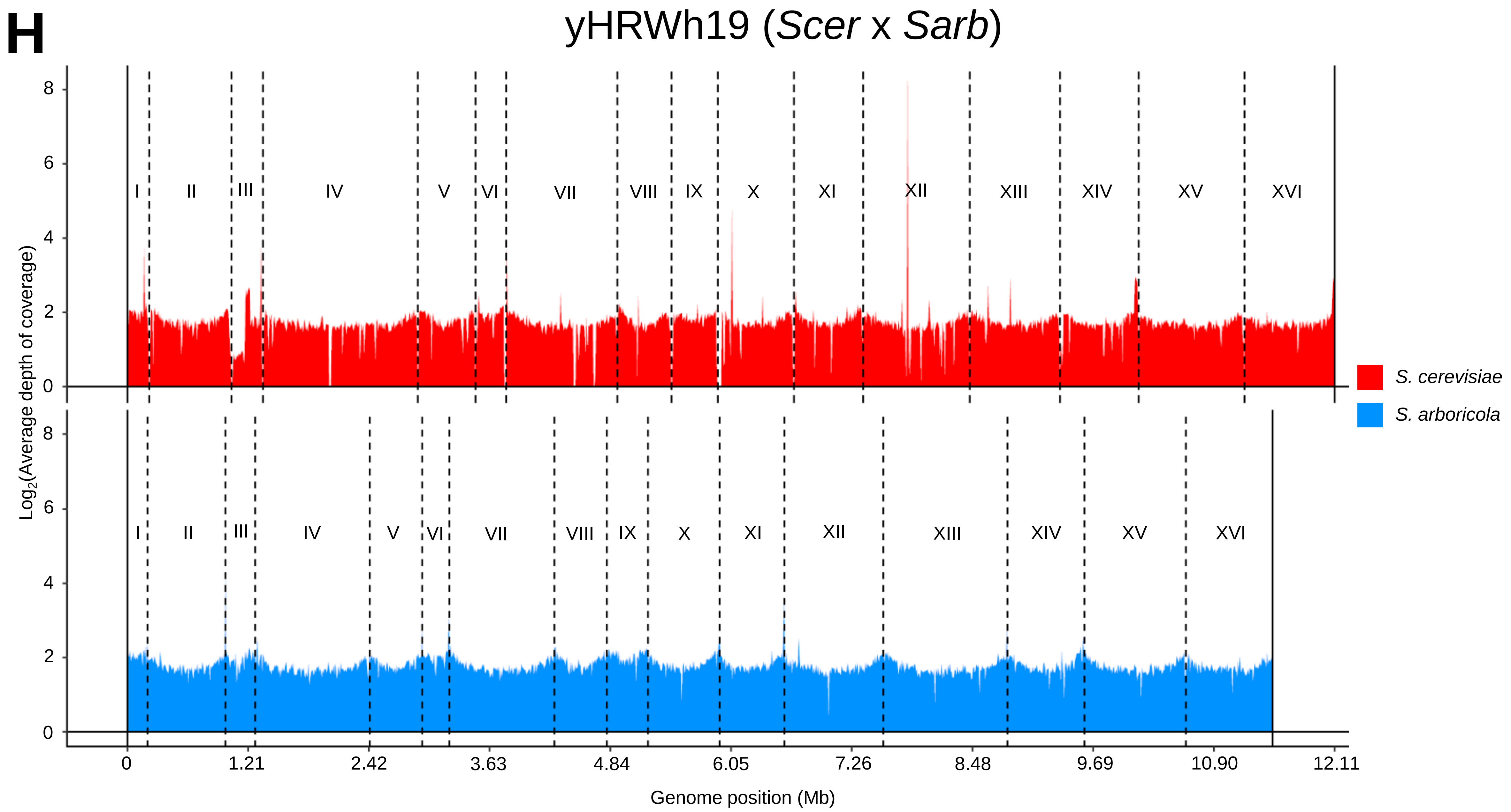
H
yHRWh19 (Scer x Sarb)
Log2(Average depth of coverage)
8
6
4
2
0
I
II
III
IV
V
VI
VII
VIII
IX
X
XI
XII
XIII
XIV
XV
XVI
S. cerevisiae
S. arboricola
0
1.21
2.42
3.63
4.84
6.05
7.26
8.48
9.69
10.90
12.11
Genome position (Mb)

**I**

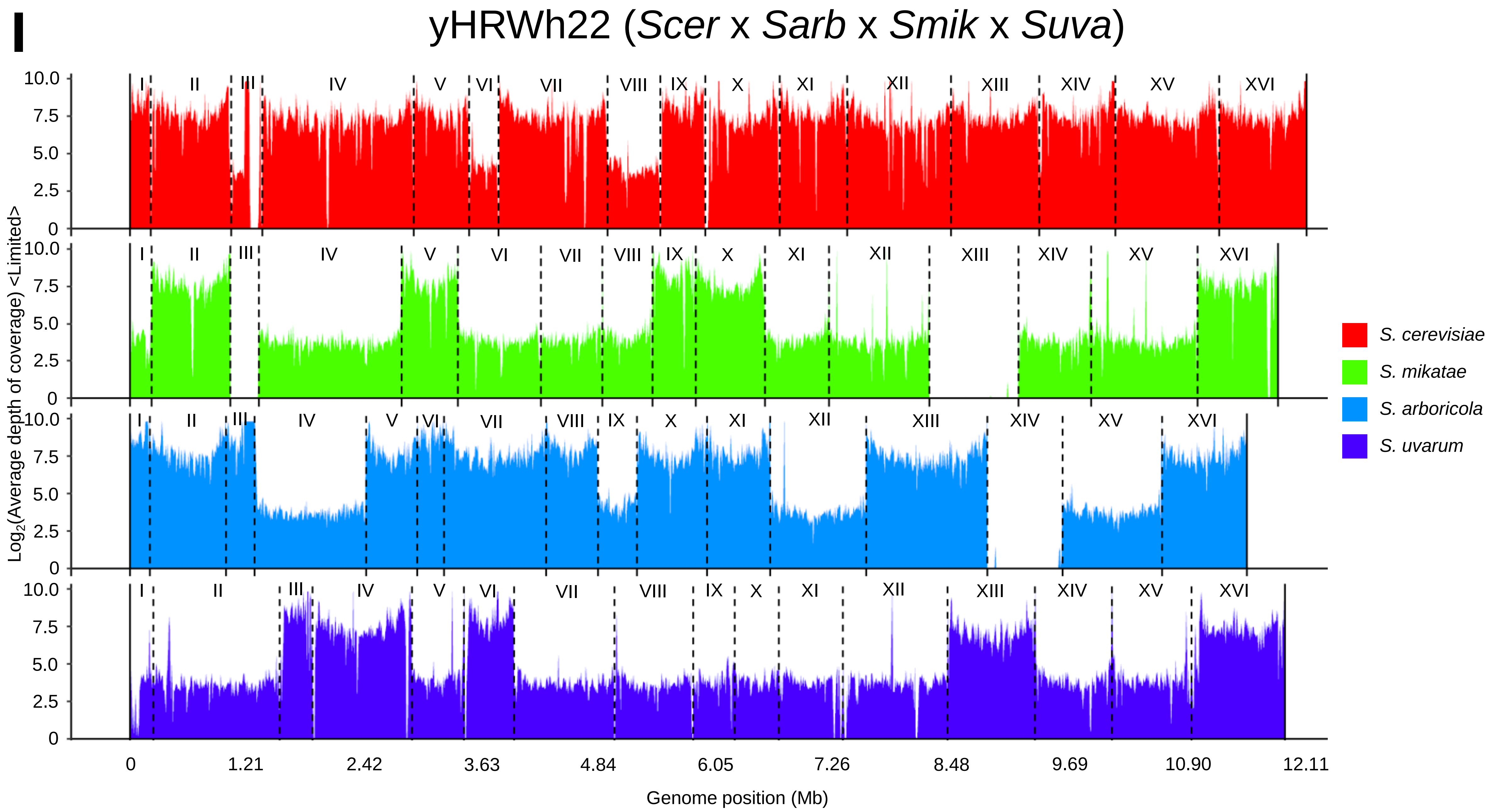
yHRWh22 (*Scer* x *Sarb* x *Smik* x *Suva*)
I II III IV V VI VII VIII IX X XI XII XIII XIV XV XVI
10.0 7.5 5.0 2.5 0
Log2(Average depth of coverage) <Limited>
S. cerevisiae
S. mikatae
S. arboricola
S. uvarum
0 1.21 2.42 3.63 4.84 6.05 7.26 8.48 9.69 10.90 12.11
Genome position (Mb)

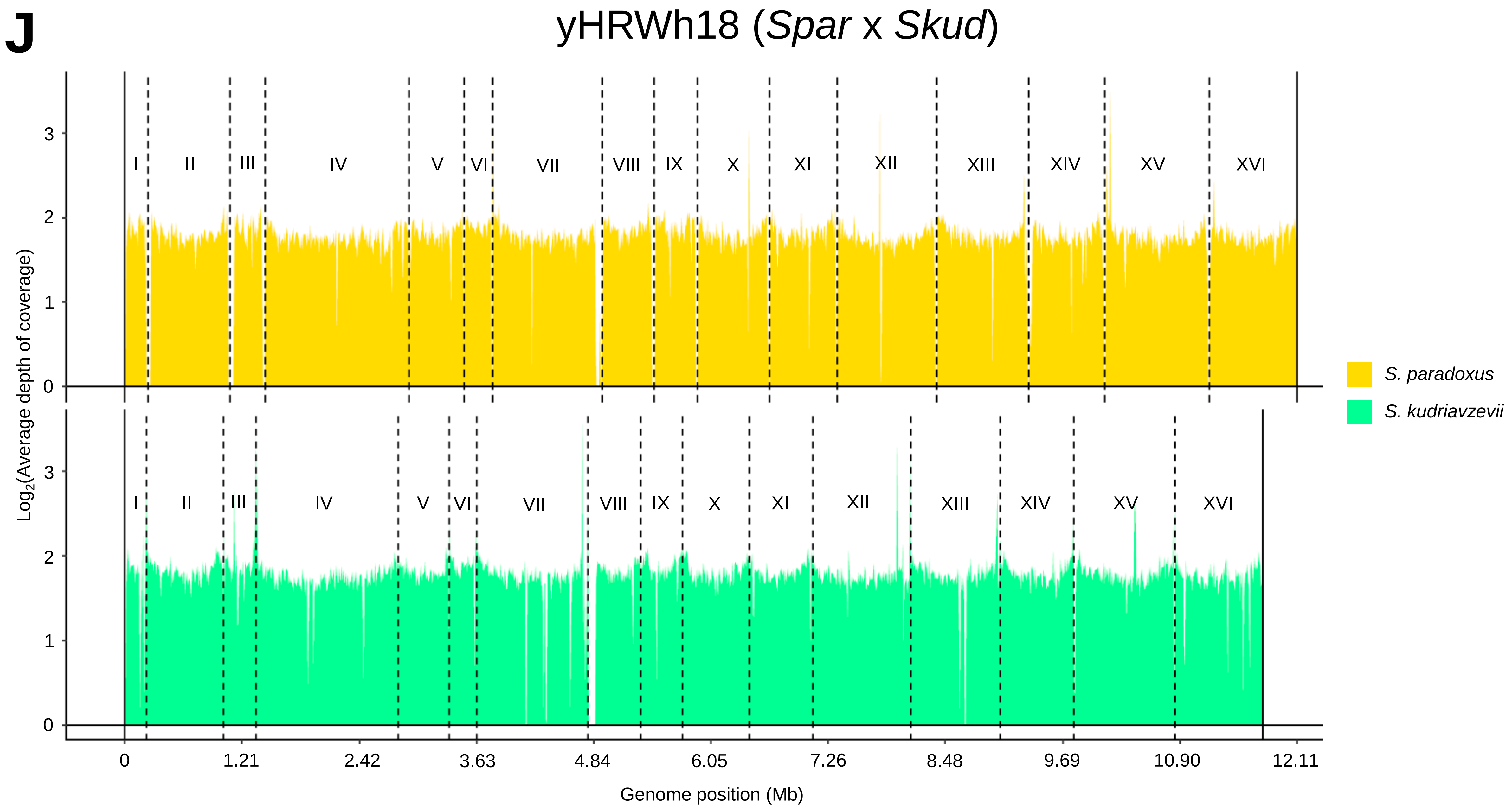
J
yHRWh18 (*Spar* x *Skud*)
Log2(Average depth of coverage)
0
1
2
3
I
II
III
IV
V
VI
VII
VIII
IX
X
XI
XII
XIII
XIV
XV
XVI
0
1.21
2.42
3.63
4.84
6.05
7.26
8.48
9.69
10.90
12.11
Genome position (Mb)
*S. paradoxus*
*S. kudriavzevii*

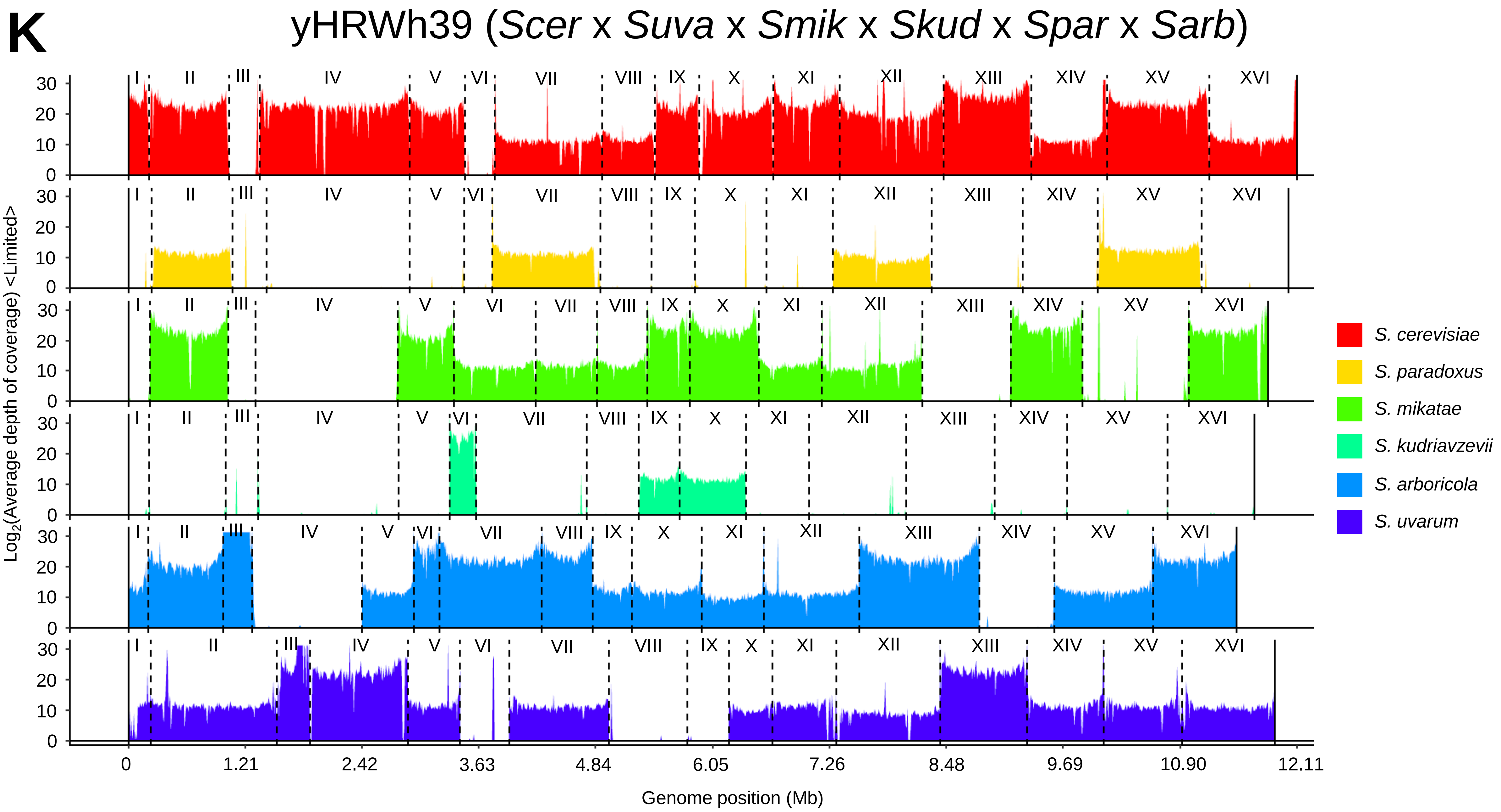
K
yHRWh39 (Scer x Suva x Smik x Skud x Spar x Sarb)
Log2(Average depth of coverage) <Limited>
Genome position (Mb)
0
1.21
2.42
3.63
4.84
6.05
7.26
8.48
9.69
10.90
12.11
S. cerevisiae
S. paradoxus
S. mikatae
S. kudriavzevii
S. arboricola
S. uvarum

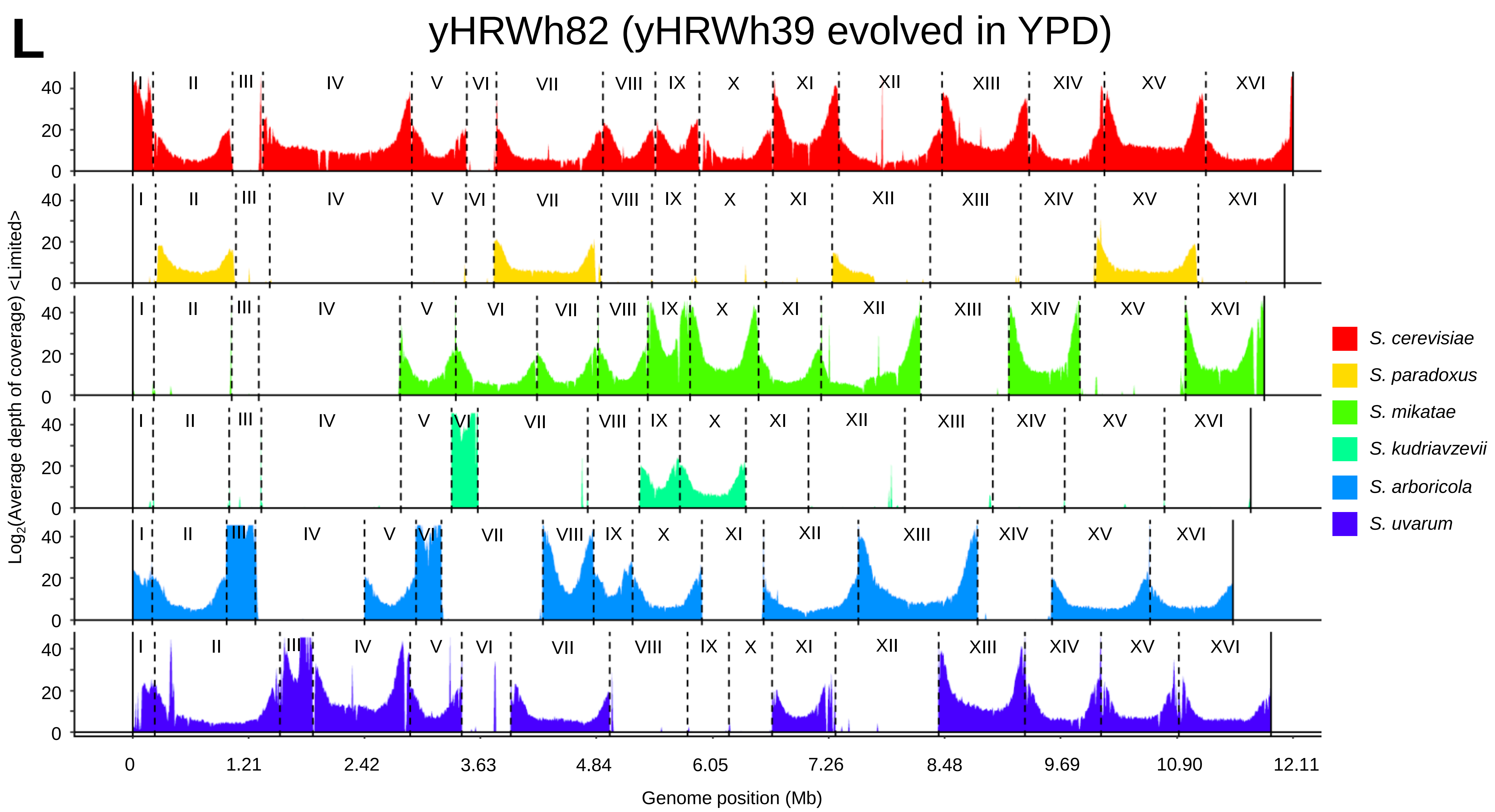

L
yHRWh82 (yHRWh39 evolved in YPD)
I
II
III
IV
V
VI
VII
VIII
IX
X
XI
XII
XIII
XIV
XV
XVI
0
20
40
Log2(Average depth of coverage) <Limited>
0
1.21
2.42
3.63
4.84
6.05
7.26
8.48
9.69
10.90
12.11
Genome position (Mb)
S. cerevisiae
S. paradoxus
S. mikatae
S. kudriavzevii
S. arboricola
S. uvarum

# M

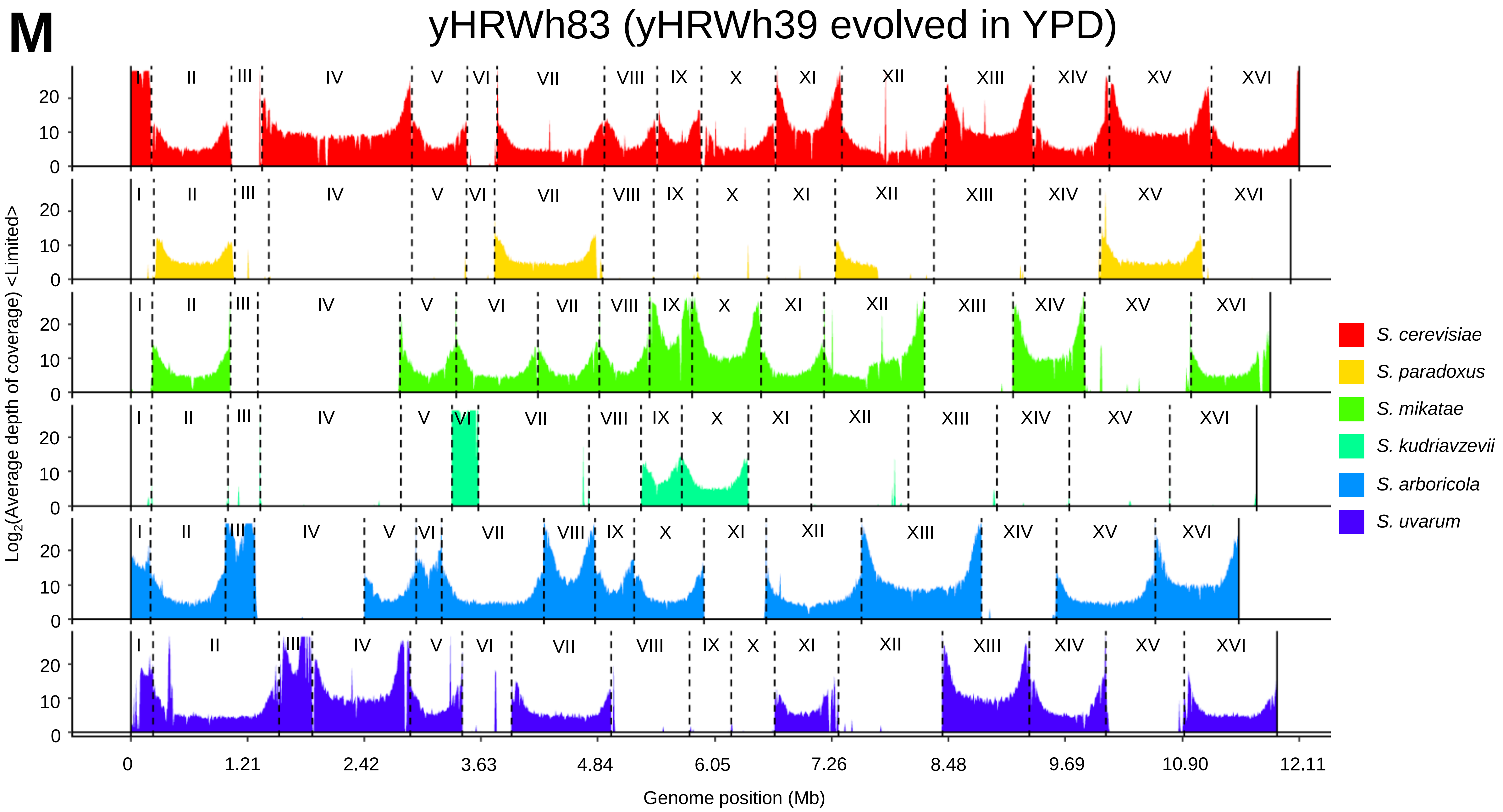
yHRWh83 (yHRWh39 evolved in YPD)
Log2(Average depth of coverage) <Limited>
Genome position (Mb)
0
1.21
2.42
3.63
4.84
6.05
7.26
8.48
9.69
10.90
12.11
I
II
III
IV
V
VI
VII
VIII
IX
X
XI
XII
XIII
XIV
XV
XVI
S. cerevisiae
S. paradoxus
S. mikatae
S. kudriavzevii
S. arboricola
S. uvarum

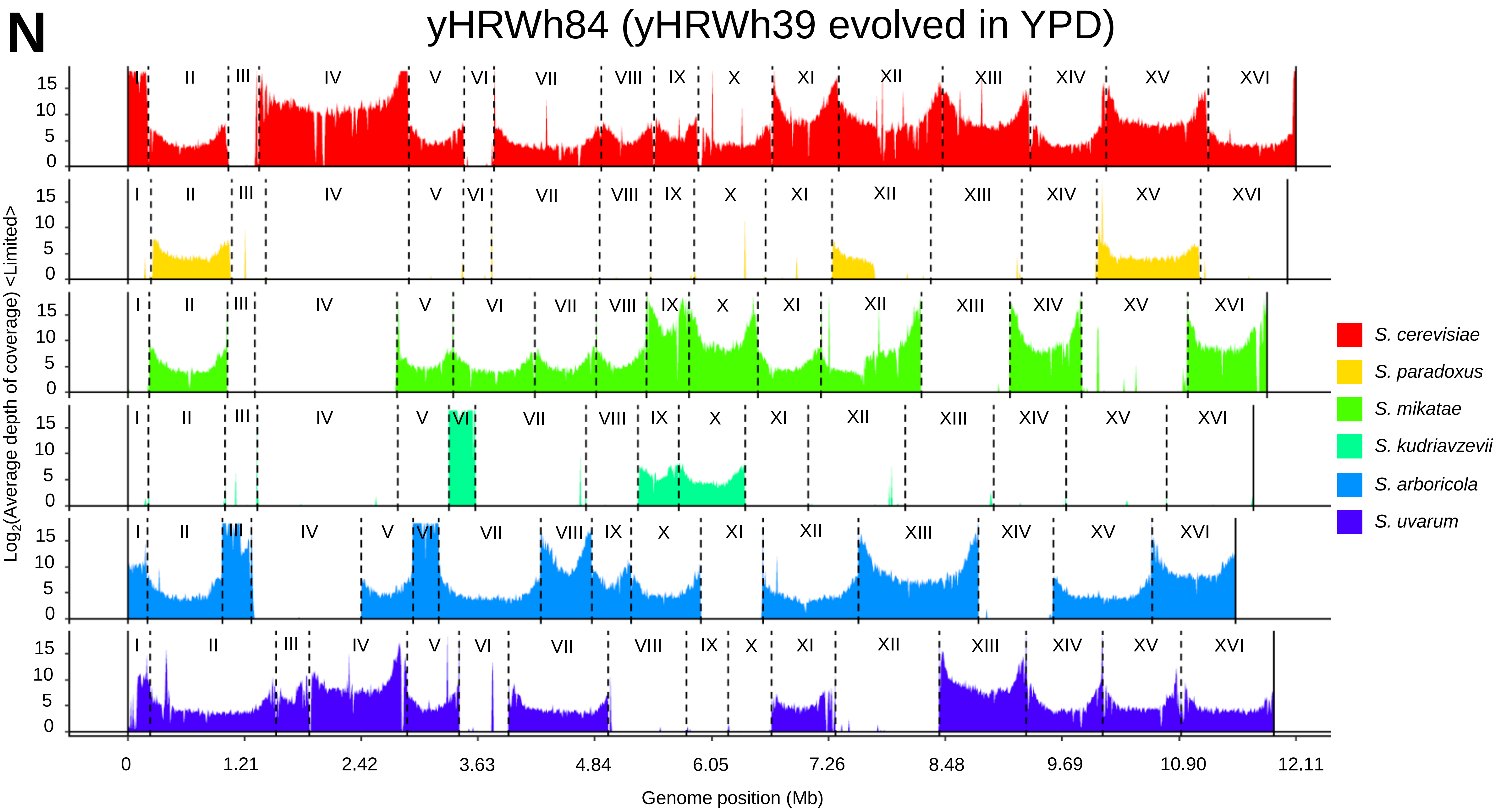
N
yHRWh84 (yHRWh39 evolved in YPD)
Log2(Average depth of coverage) <Limited>
15
10
5
0
I
II
III
IV
V
VI
VII
VIII
IX
X
XI
XII
XIII
XIV
XV
XVI
S. cerevisiae
S. paradoxus
S. mikatae
S. kudriavzevii
S. arboricola
S. uvarum
0
1.21
2.42
3.63
4.84
6.05
7.26
8.48
9.69
10.90
12.11
Genome position (Mb)

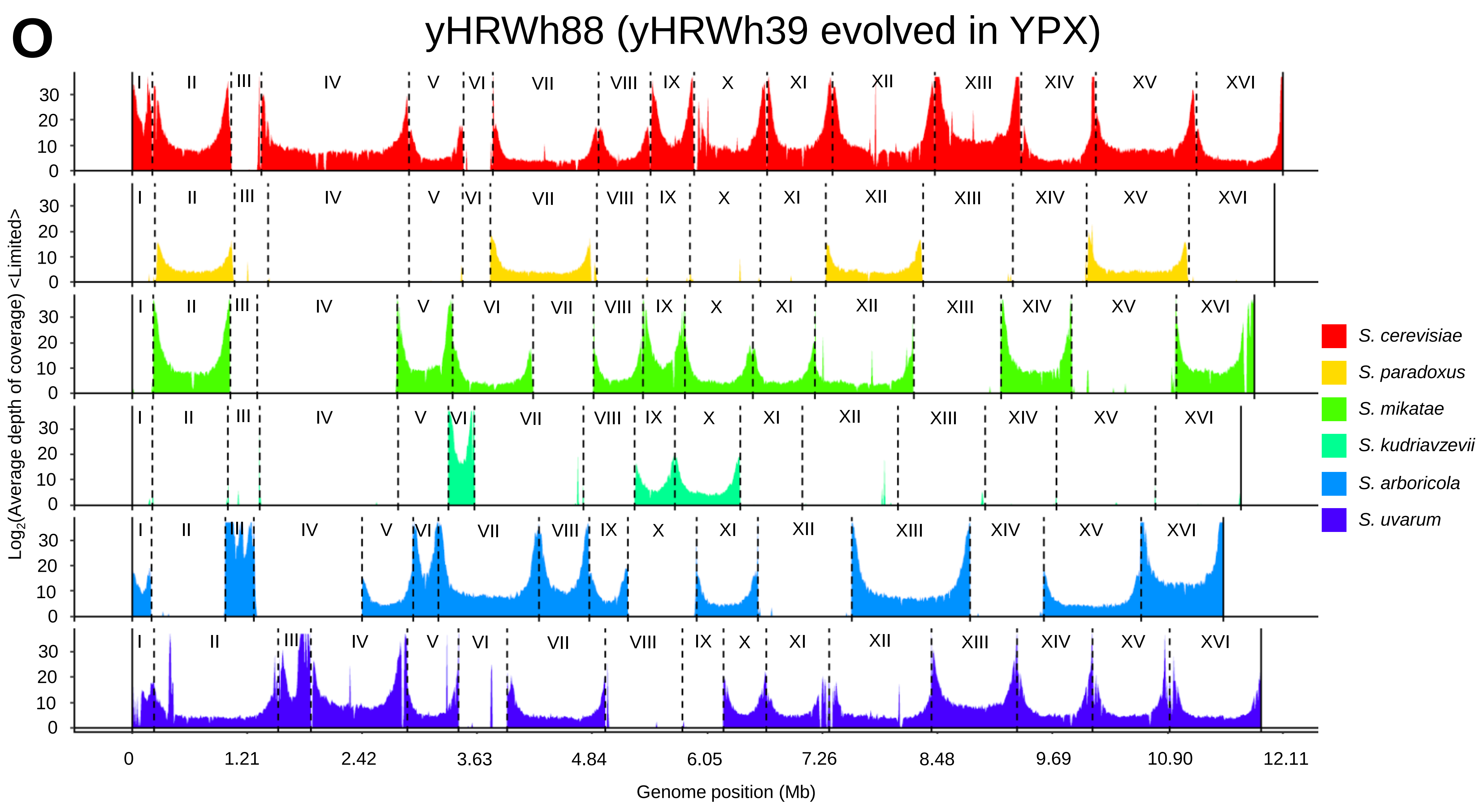
O
yHRWh88 (yHRWh39 evolved in YPX)
Log2(Average depth of coverage) <Limited>
Genome position (Mb)
0
1.21
2.42
3.63
4.84
6.05
7.26
8.48
9.69
10.90
12.11
I
II
III
IV
V
VI
VII
VIII
IX
X
XI
XII
XIII
XIV
XV
XVI
S. cerevisiae
S. paradoxus
S. mikatae
S. kudriavzevii
S. arboricola
S. uvarum

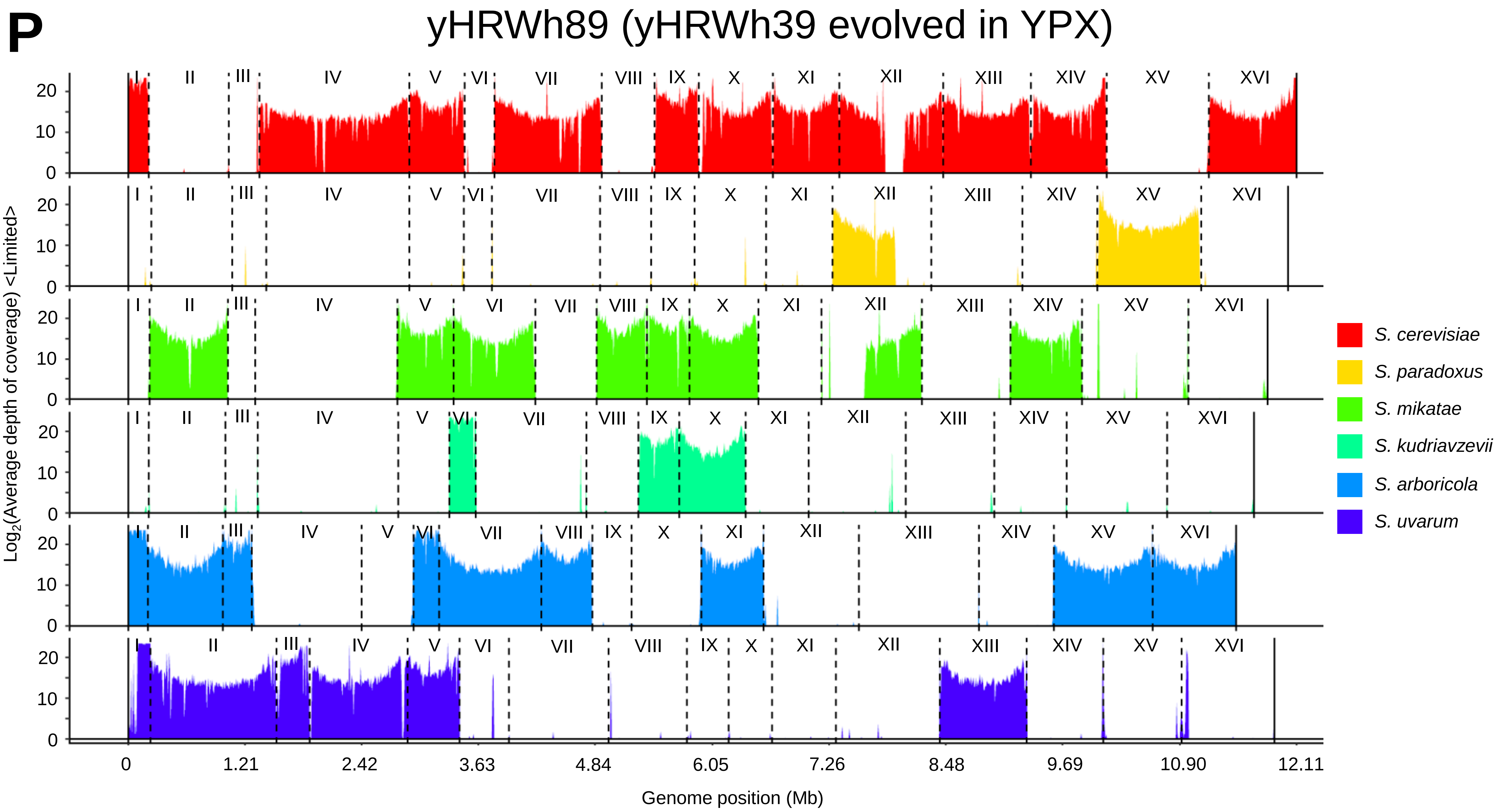

P
yHRWh89 (yHRWh39 evolved in YPX)
I II III IV V VI VII VIII IX X XI XII XIII XIV XV XVI
Log2(Average depth of coverage) <Limited>
0 10 20
0 1.21 2.42 3.63 4.84 6.05 7.26 8.48 9.69 10.90 12.11
Genome position (Mb)
S. cerevisiae
S. paradoxus
S. mikatae
S. kudriavzevii
S. arboricola
S. uvarum

**Q**

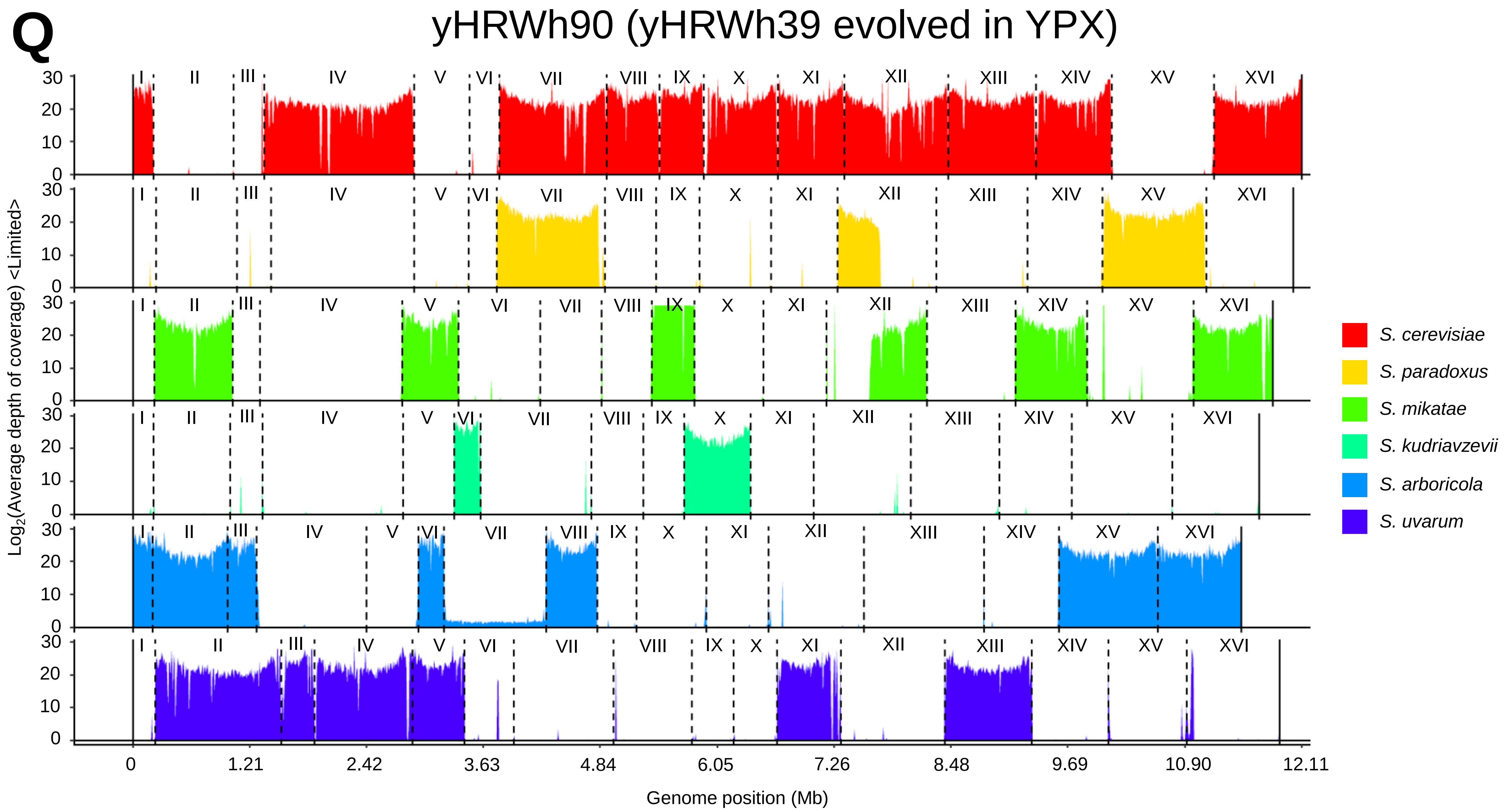
yHRWh90 (yHRWh39 evolved in YPX)
Log2(Average depth of coverage) <Limited>
Genome position (Mb)
0
1.21
2.42
3.63
4.84
6.05
7.26
8.48
9.69
10.90
12.11
I
II
III
IV
V
VI
VII
VIII
IX
X
XI
XII
XIII
XIV
XV
XVI
S. cerevisiae
S. paradoxus
S. mikatae
S. kudriavzevii
S. arboricola
S. uvarum

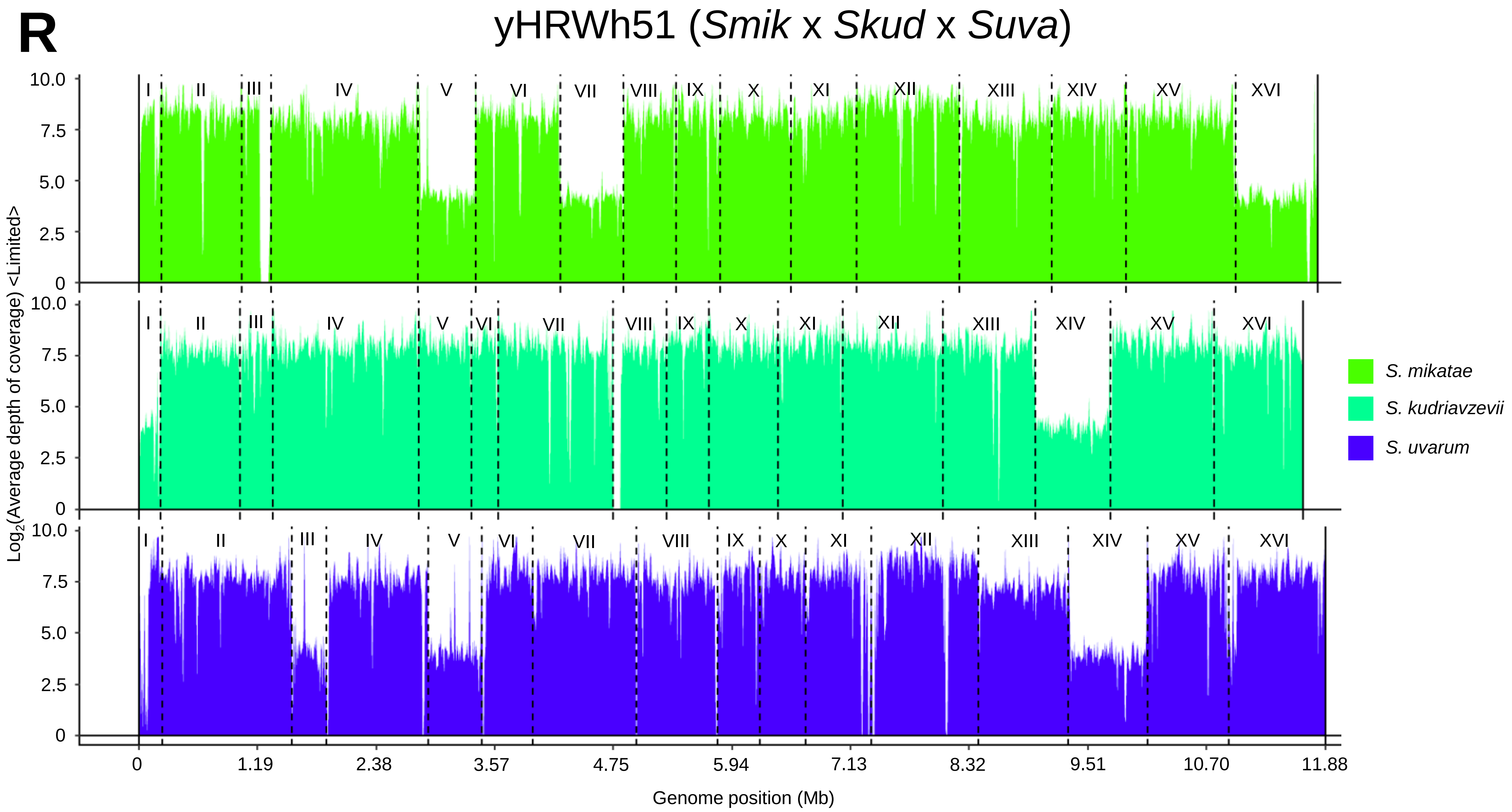
R
yHRWh51 (Smik x Skud x Suva)
Log2(Average depth of coverage) <Limited>
10.0
7.5
5.0
2.5
0
I II III IV V VI VII VIII IX X XI XII XIII XIV XV XVI
S. mikatae
S. kudriavzevii
S. uvarum
0 1.19 2.38 3.57 4.75 5.94 7.13 8.32 9.51 10.70 11.88
Genome position (Mb)

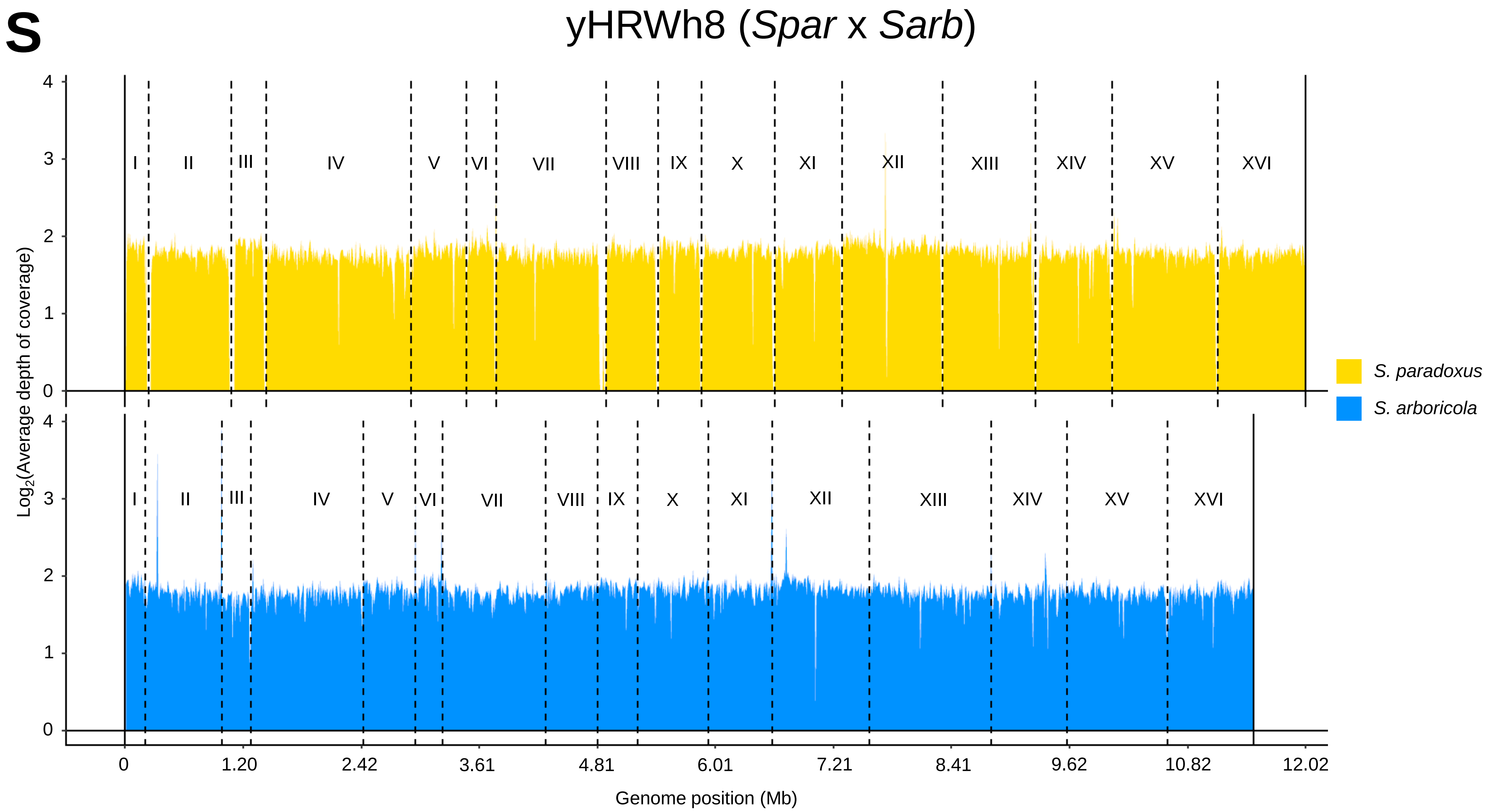
S
yHRWh8 (*Spar* x *Sarb*)
I
II
III
IV
V
VI
VII
VIII
IX
X
XI
XII
XIII
XIV
XV
XVI
Log2(Average depth of coverage)
0
1
2
3
4
0
1.20
2.42
3.61
4.81
6.01
7.21
8.41
9.62
10.82
12.02
Genome position (Mb)
*S. paradoxus*
*S. arboricola*

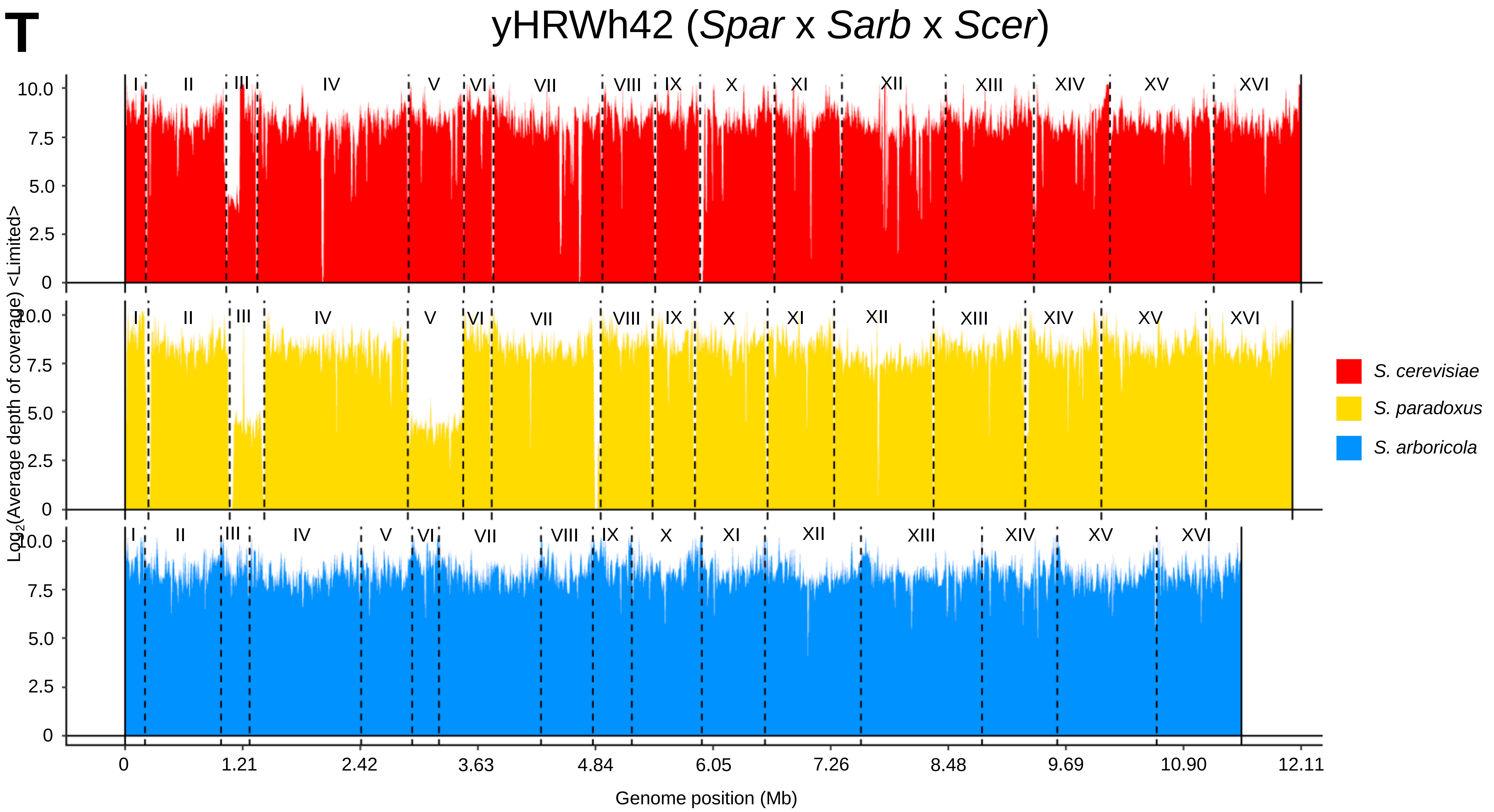
T
yHRWh42 (Spar x Sarb x Scer)
I
II
III
IV
V
VI
VII
VIII
IX
X
XI
XII
XIII
XIV
XV
XVI
10.0
7.5
5.0
2.5
0
Log2(Average depth of coverage) <Limited>
S. cerevisiae
S. paradoxus
S. arboricola
0
1.21
2.42
3.63
4.84
6.05
7.26
8.48
9.69
10.90
12.11
Genome position (Mb)

**U** yHRWh56 (*Scer* x *Suva* x *Smik* x *Skud* x *Spar* x *Sarb*)

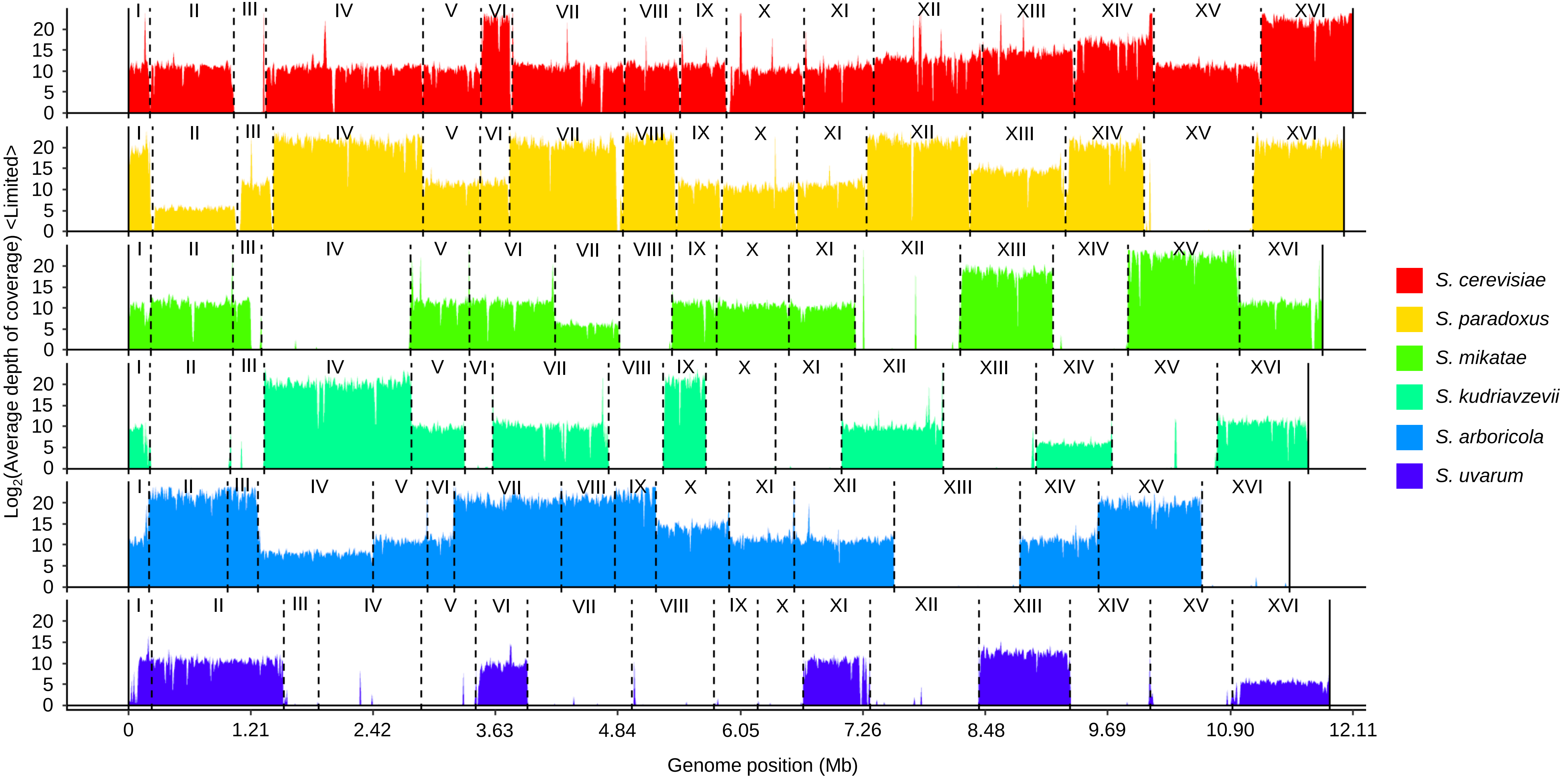

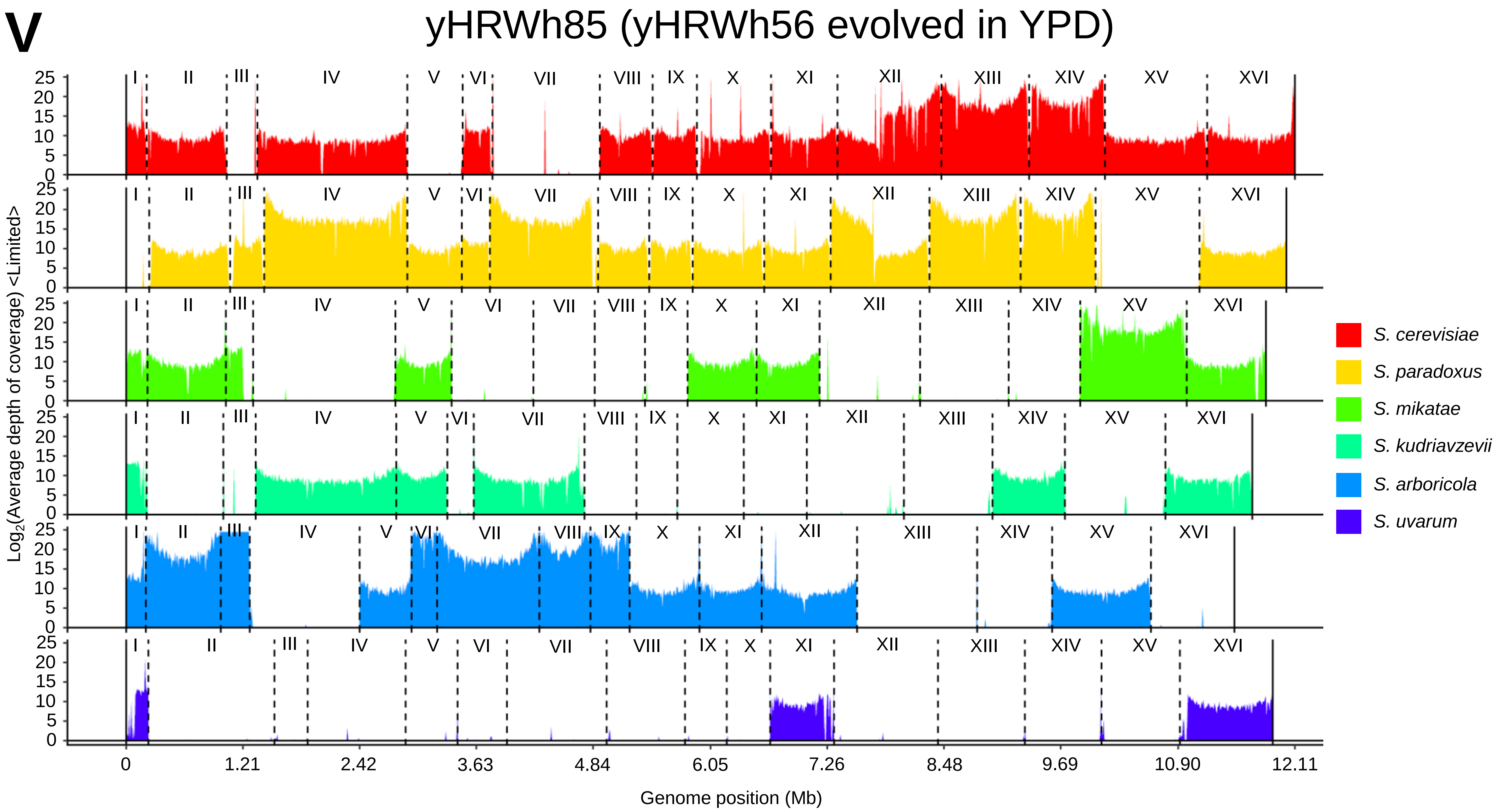
V
yHRWh85 (yHRWh56 evolved in YPD)
Log2(Average depth of coverage) <Limited>
I II III IV V VI VII VIII IX X XI XII XIII XIV XV XVI
S. cerevisiae
S. paradoxus
S. mikatae
S. kudriavzevii
S. arboricola
S. uvarum
0 1.21 2.42 3.63 4.84 6.05 7.26 8.48 9.69 10.90 12.11
Genome position (Mb)

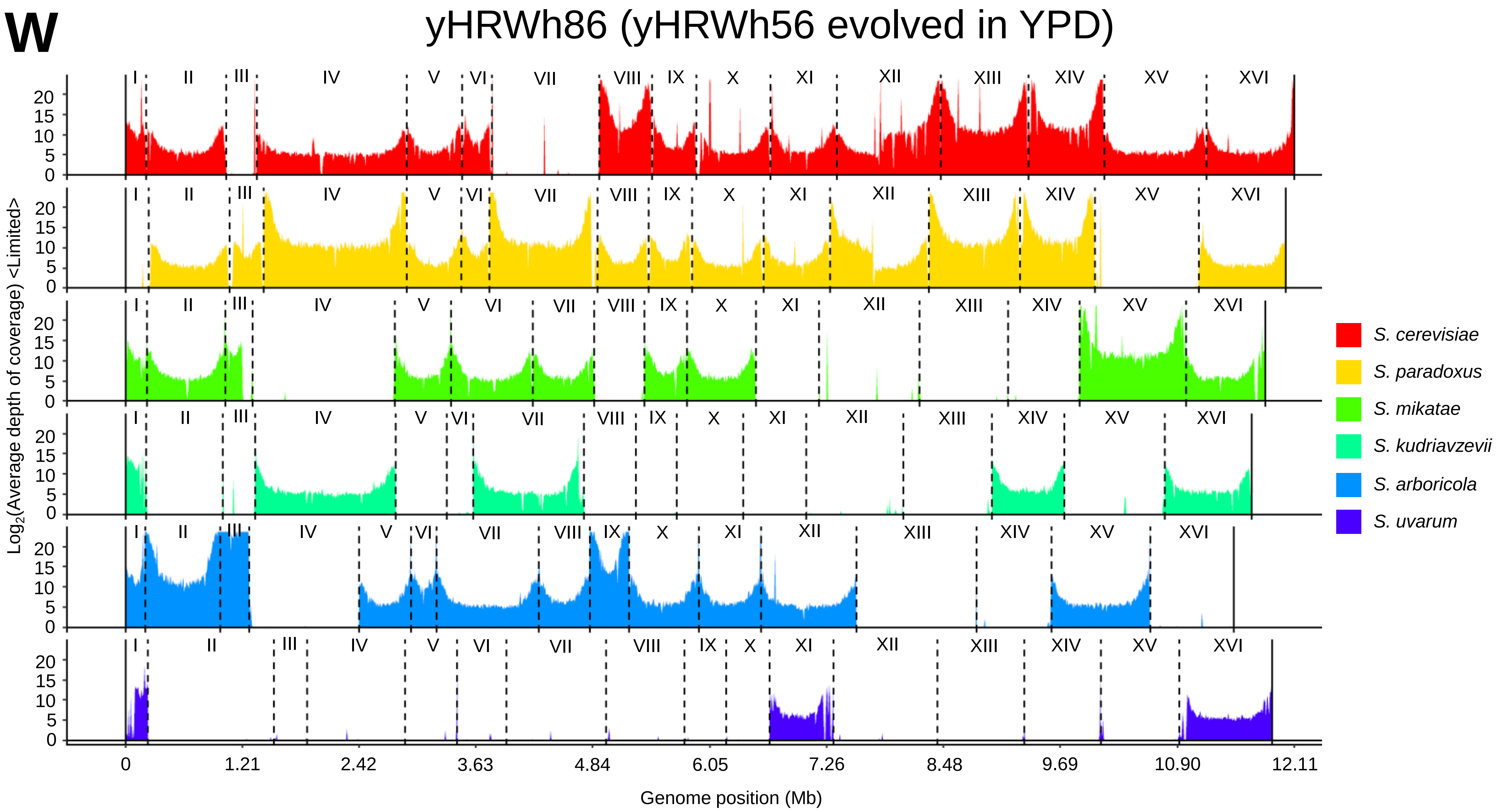
W
yHRWh86 (yHRWh56 evolved in YPD)
Log2(Average depth of coverage) <Limited>
I
II
III
IV
V
VI
VII
VIII
IX
X
XI
XII
XIII
XIV
XV
XVI
0
5
10
15
20
0
1.21
2.42
3.63
4.84
6.05
7.26
8.48
9.69
10.90
12.11
Genome position (Mb)
S. cerevisiae
S. paradoxus
S. mikatae
S. kudriavzevii
S. arboricola
S. uvarum

**X**

yHRWh87 (yHRWh56 evolved in YPD)

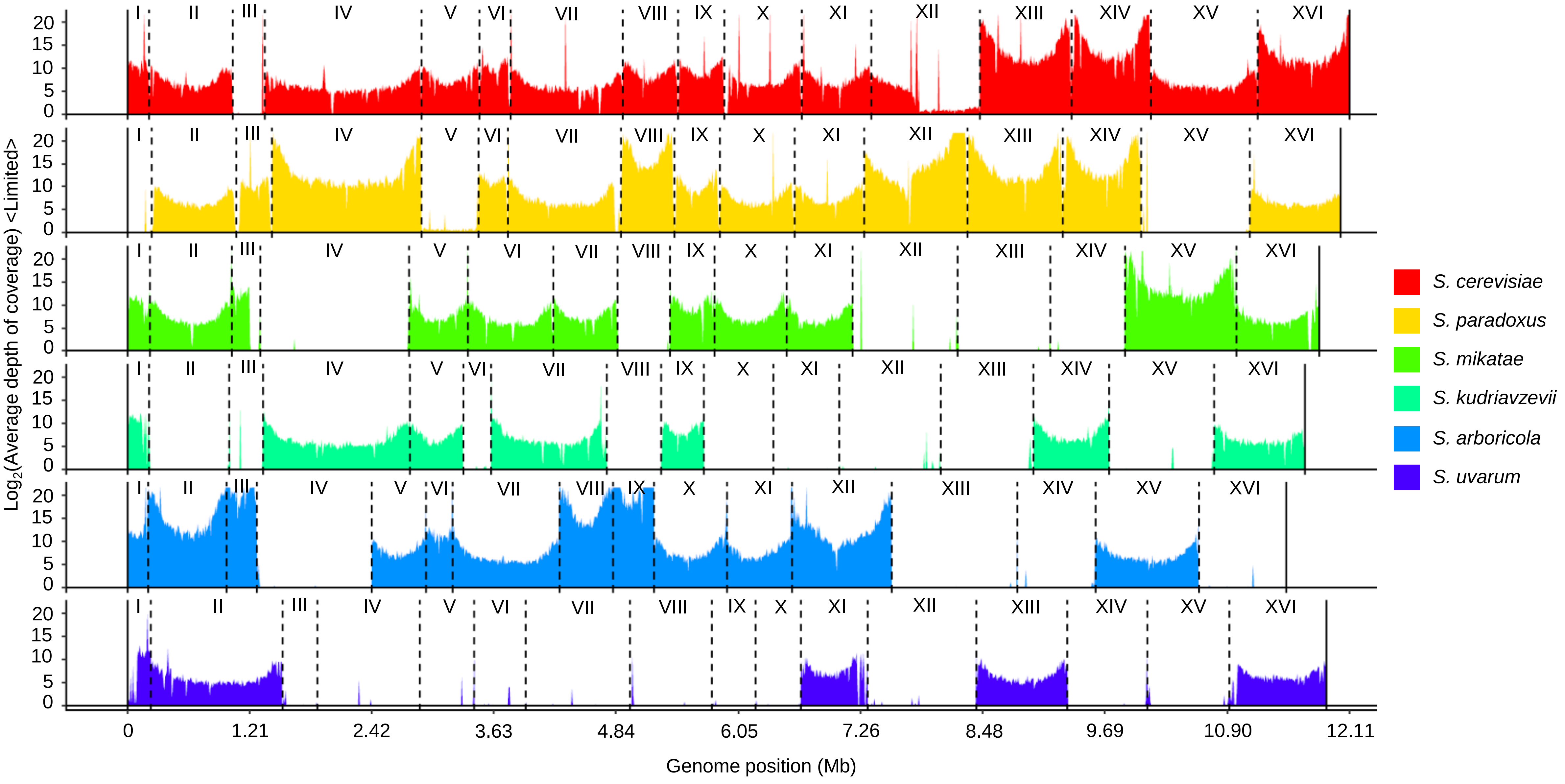

**Y**

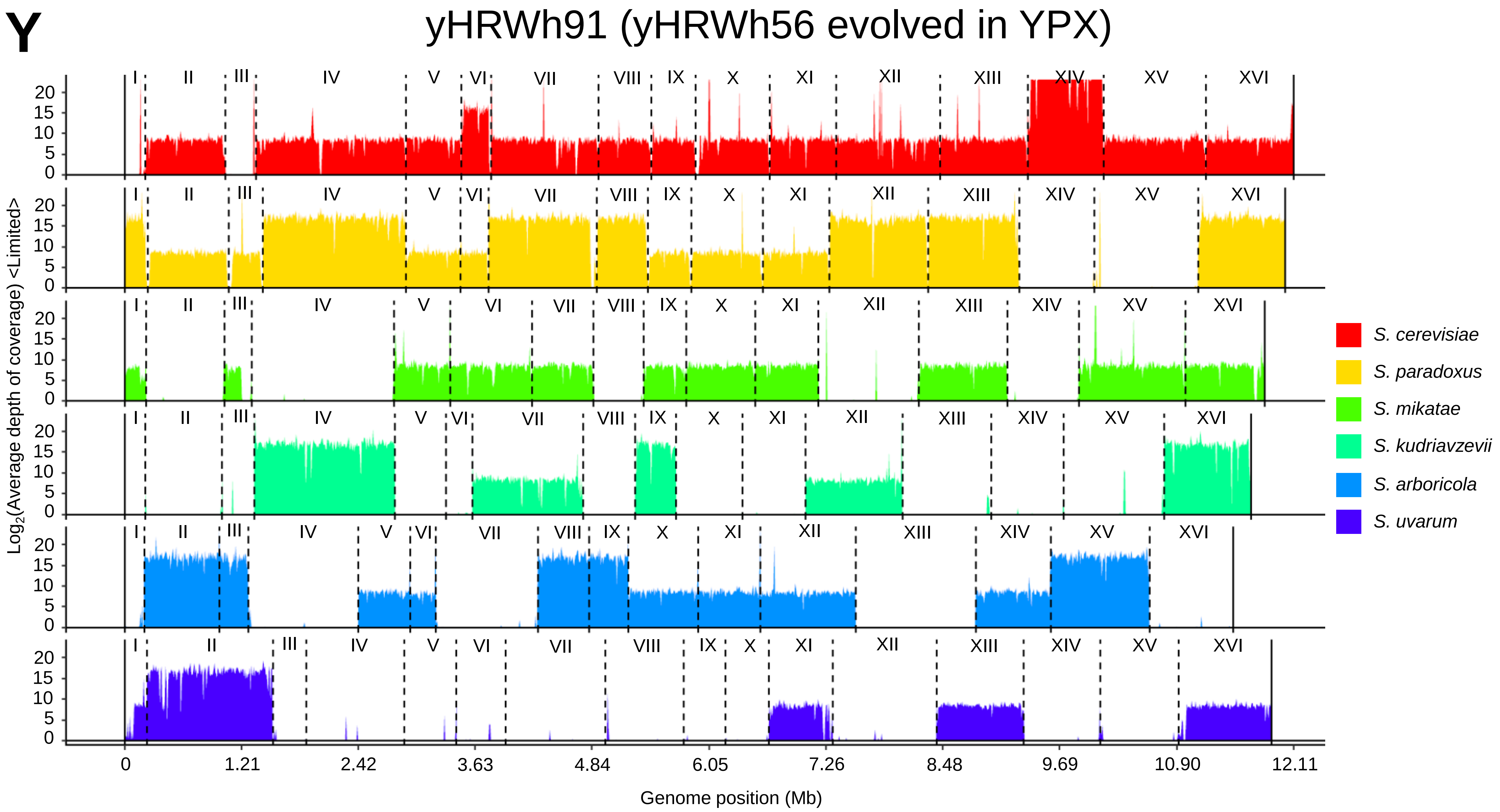

yHRWh91 (yHRWh56 evolved in YPX)
Log2(Average depth of coverage) <Limited>
I
II
III
IV
V
VI
VII
VIII
IX
X
XI
XII
XIII
XIV
XV
XVI
0
5
10
15
20
0
1.21
2.42
3.63
4.84
6.05
7.26
8.48
9.69
10.90
12.11
Genome position (Mb)
S. cerevisiae
S. paradoxus
S. mikatae
S. kudriavzevii
S. arboricola
S. uvarum

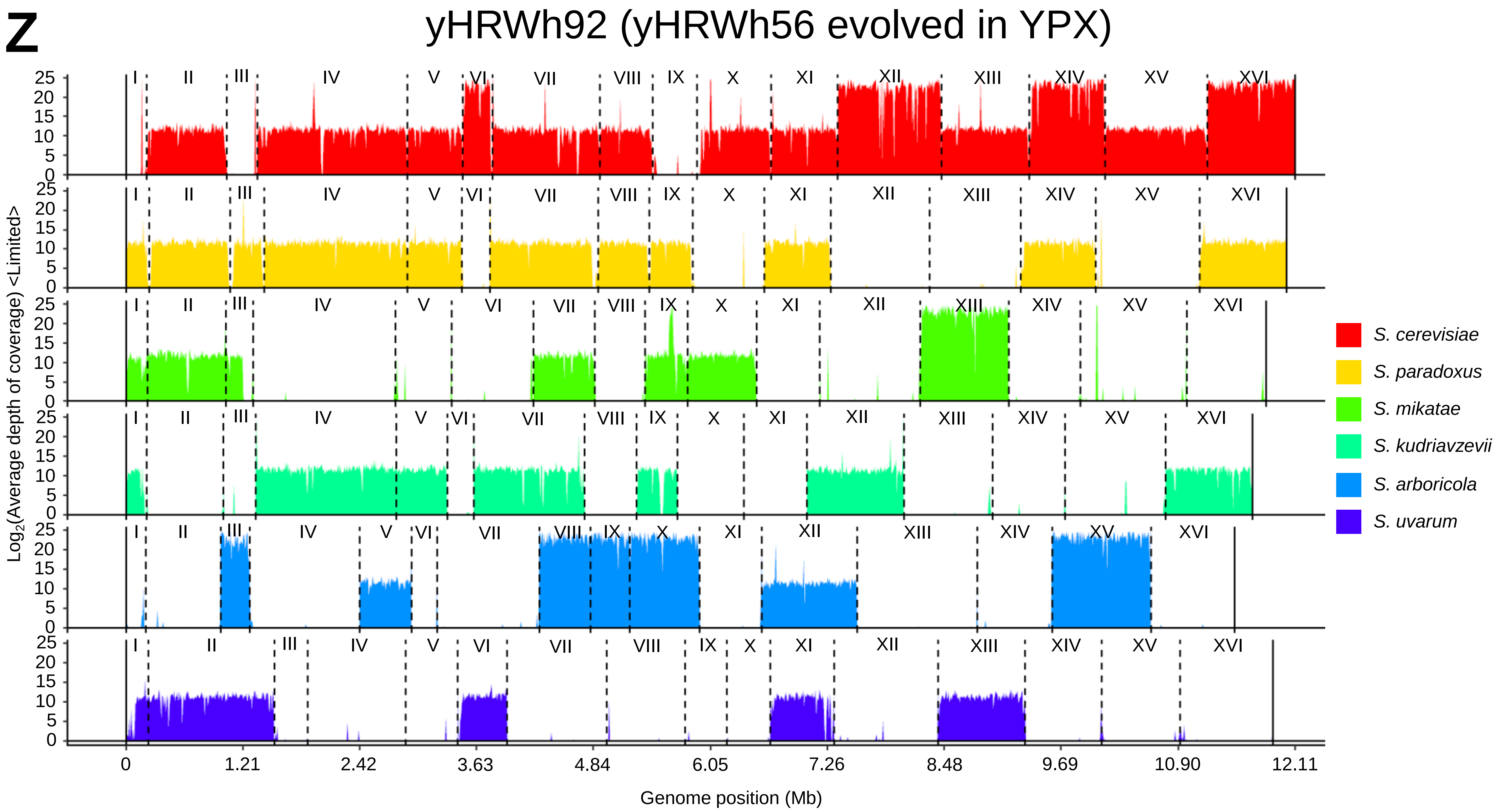
Z
yHRWh92 (yHRWh56 evolved in YPX)
Log2(Average depth of coverage) <Limited>
Genome position (Mb)
0
1.21
2.42
3.63
4.84
6.05
7.26
8.48
9.69
10.90
12.11
S. cerevisiae
S. paradoxus
S. mikatae
S. kudriavzevii
S. arboricola
S. uvarum

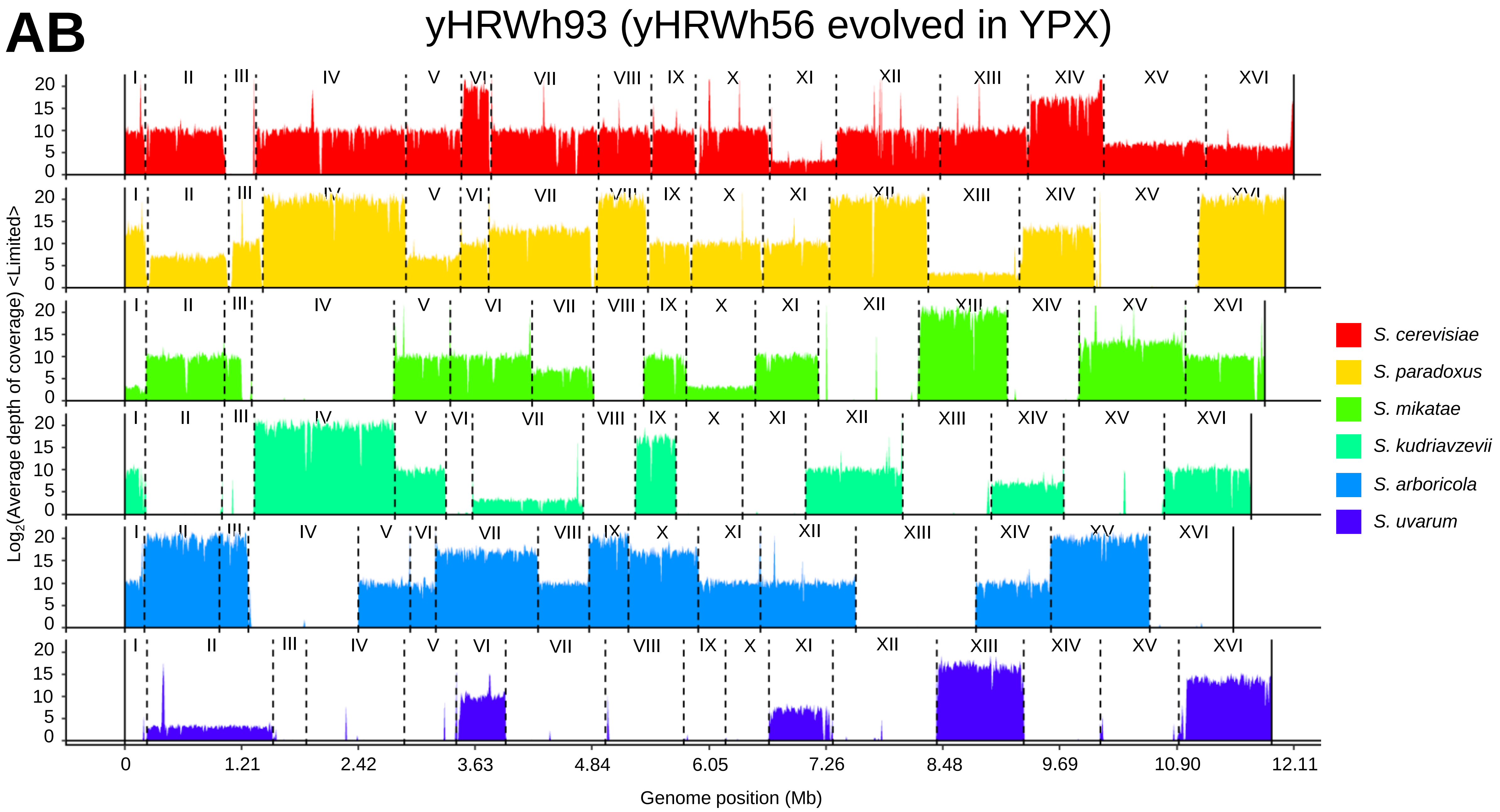
AB
yHRWh93 (yHRWh56 evolved in YPX)
Log2(Average depth of coverage) <Limited>
Genome position (Mb)
0
1.21
2.42
3.63
4.84
6.05
7.26
8.48
9.69
10.90
12.11
S. cerevisiae
S. paradoxus
S. mikatae
S. kudriavzevii
S. arboricola
S. uvarum

### Supplementary figure 4

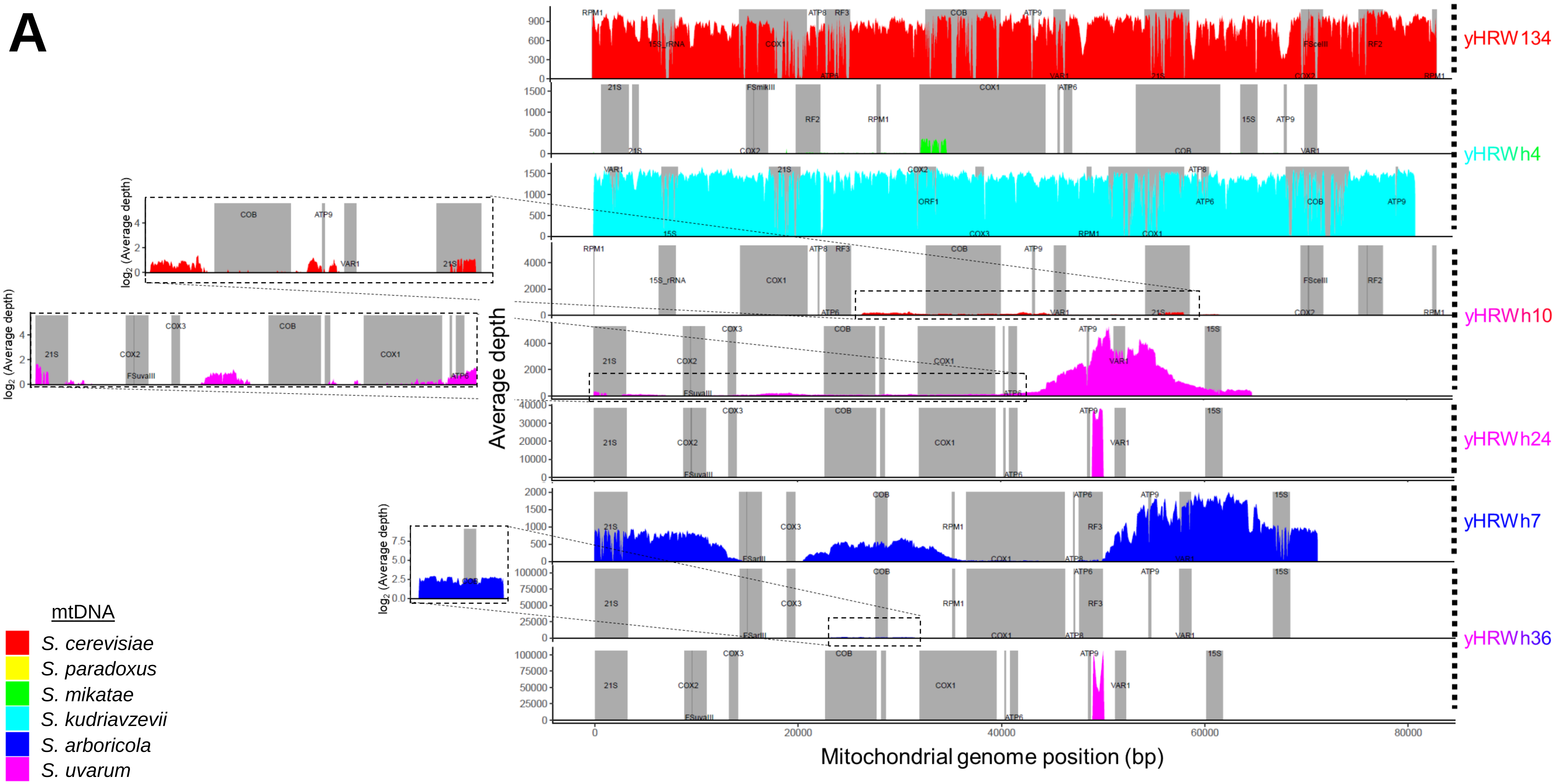

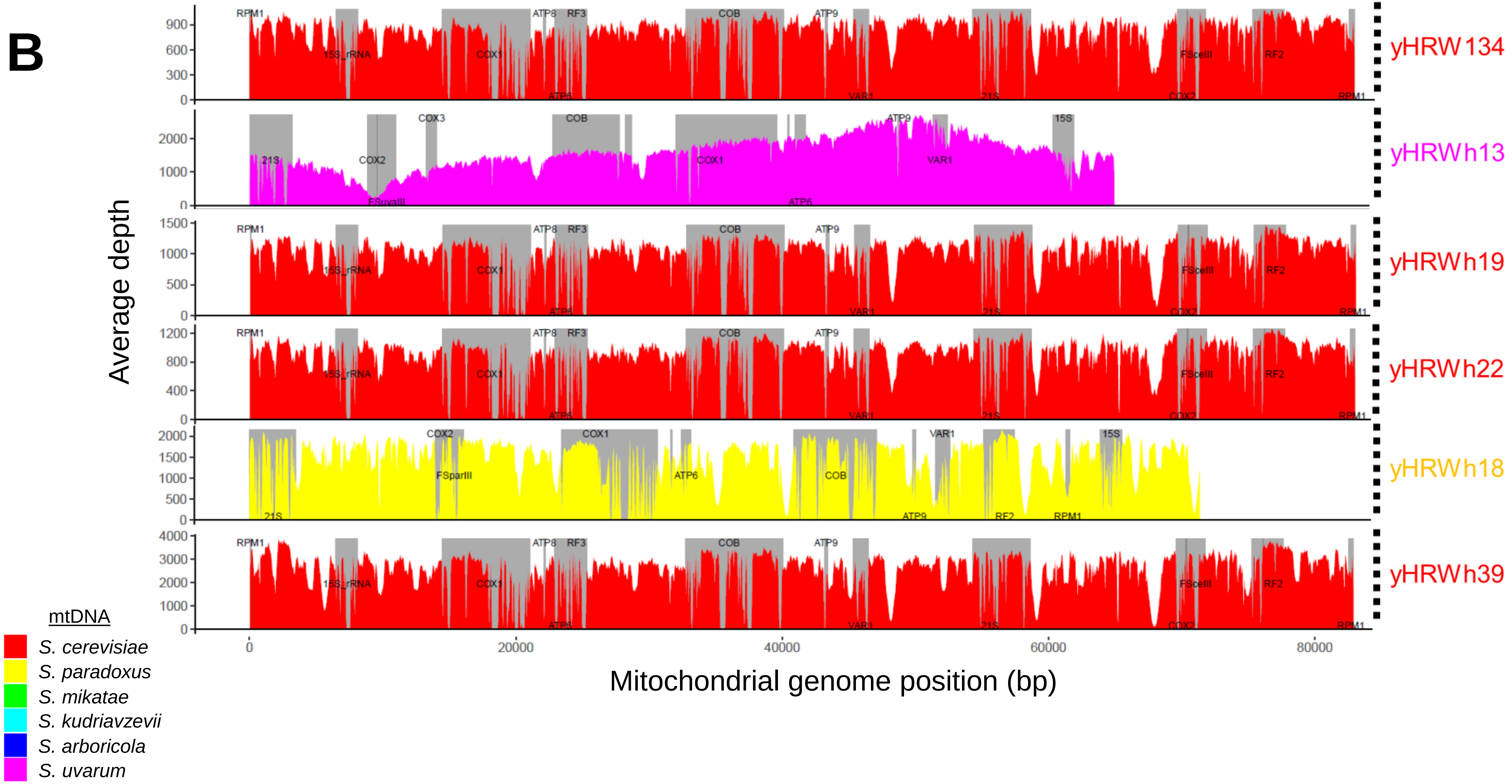
B
Average depth
Mitochondrial genome position (bp)
0
20000
40000
60000
80000
yHRW134
yHRWh13
yHRWh19
yHRWh22
yHRWh18
yHRWh39
RPM1
15S_rRNA
COX1
ATP8
RF3
ATP6
COB
ATP9
VAR1
21S
FScelII
COX2
RF2
21S
COX2
FSuvaIII
COX3
COB
COX1
ATP6
ATP9
VAR1
15S
FSparIII
ATP6
COB
RPM1
mtDNA
S. cerevisiae
S. paradoxus
S. mikatae
S. kudriavzevii
S. arboricola
S. uvarum

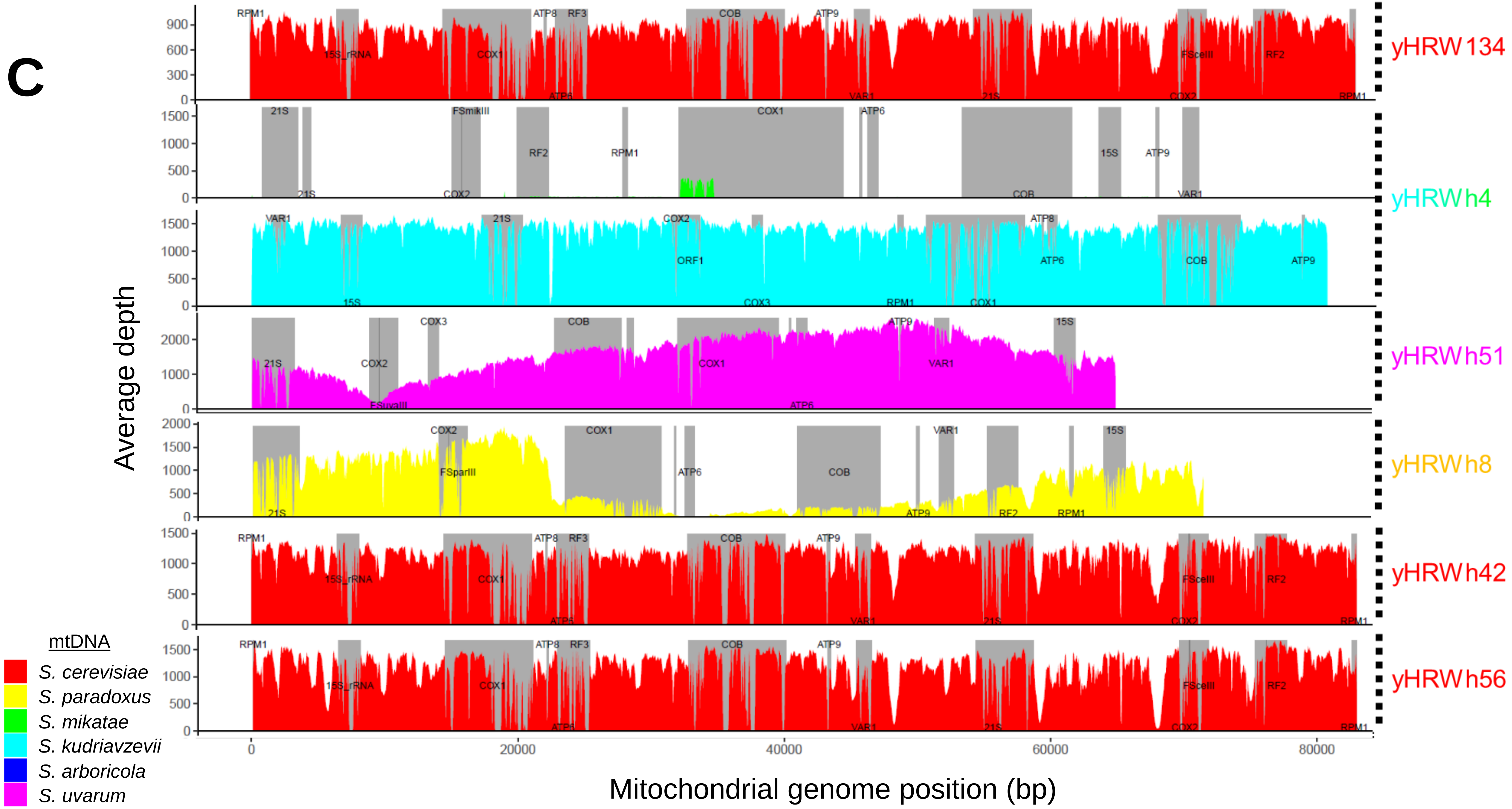
C
Average depth
Mitochondrial genome position (bp)
yHRW134
yHRWh4
yHRWh51
yHRWh8
yHRWh42
yHRWh56
mtDNA
S. cerevisiae
S. paradoxus
S. mikatae
S. kudriavzevii
S. arboricola
S. uvarum
0
20000
40000
60000
80000

### Supplementary figure 5

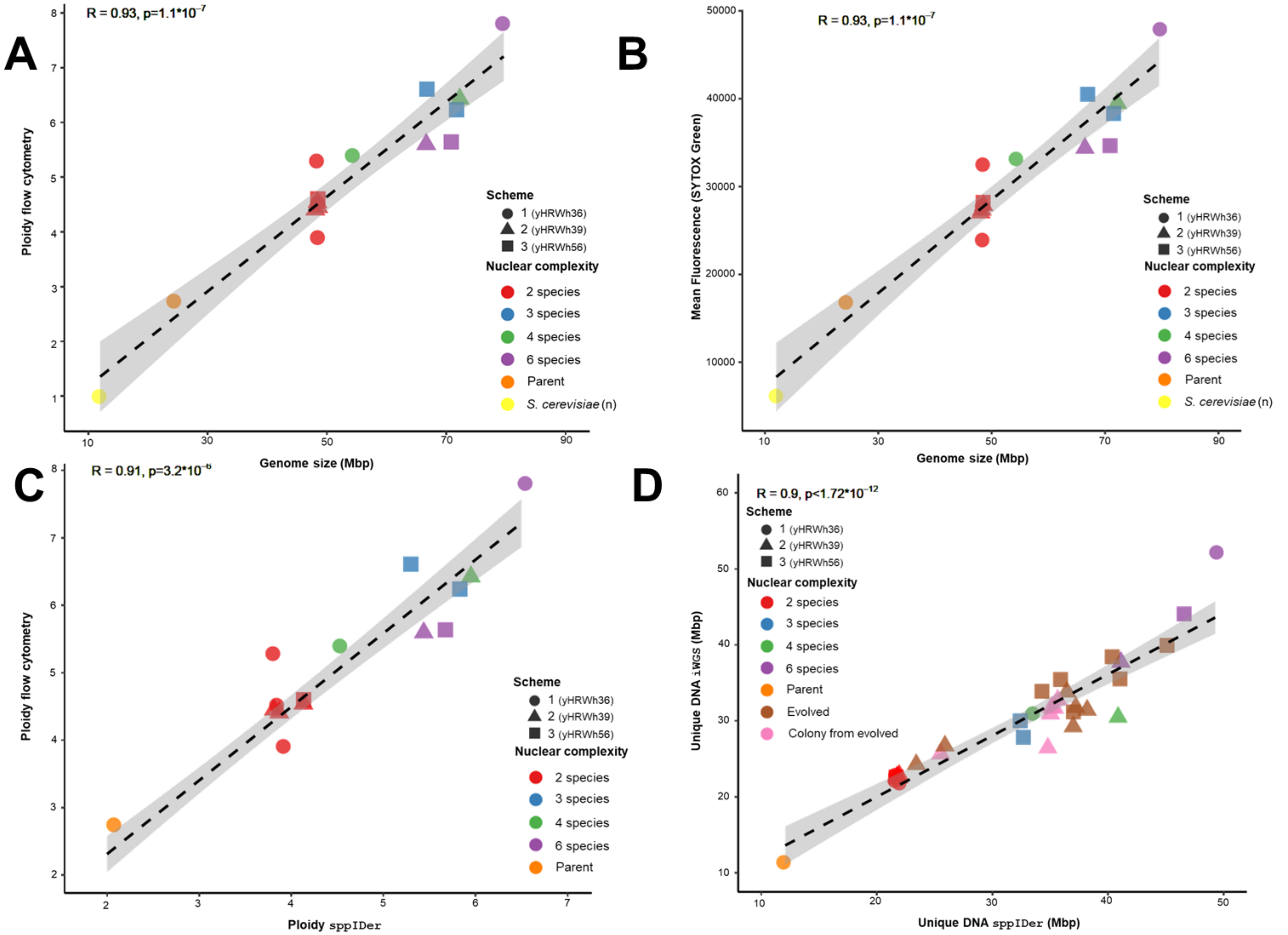

**Supplementary figure 6**

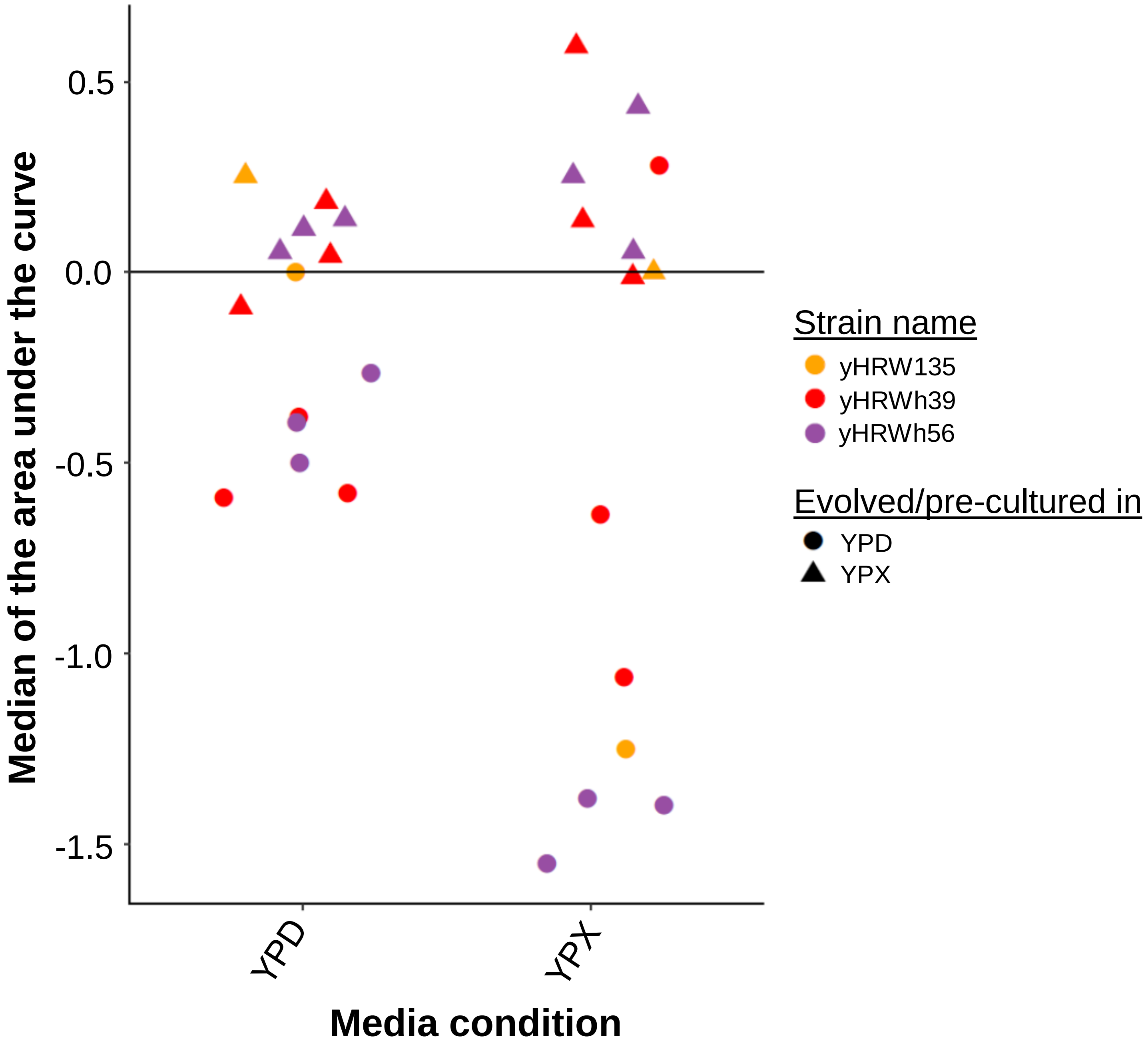

#### Supplementary figure 7

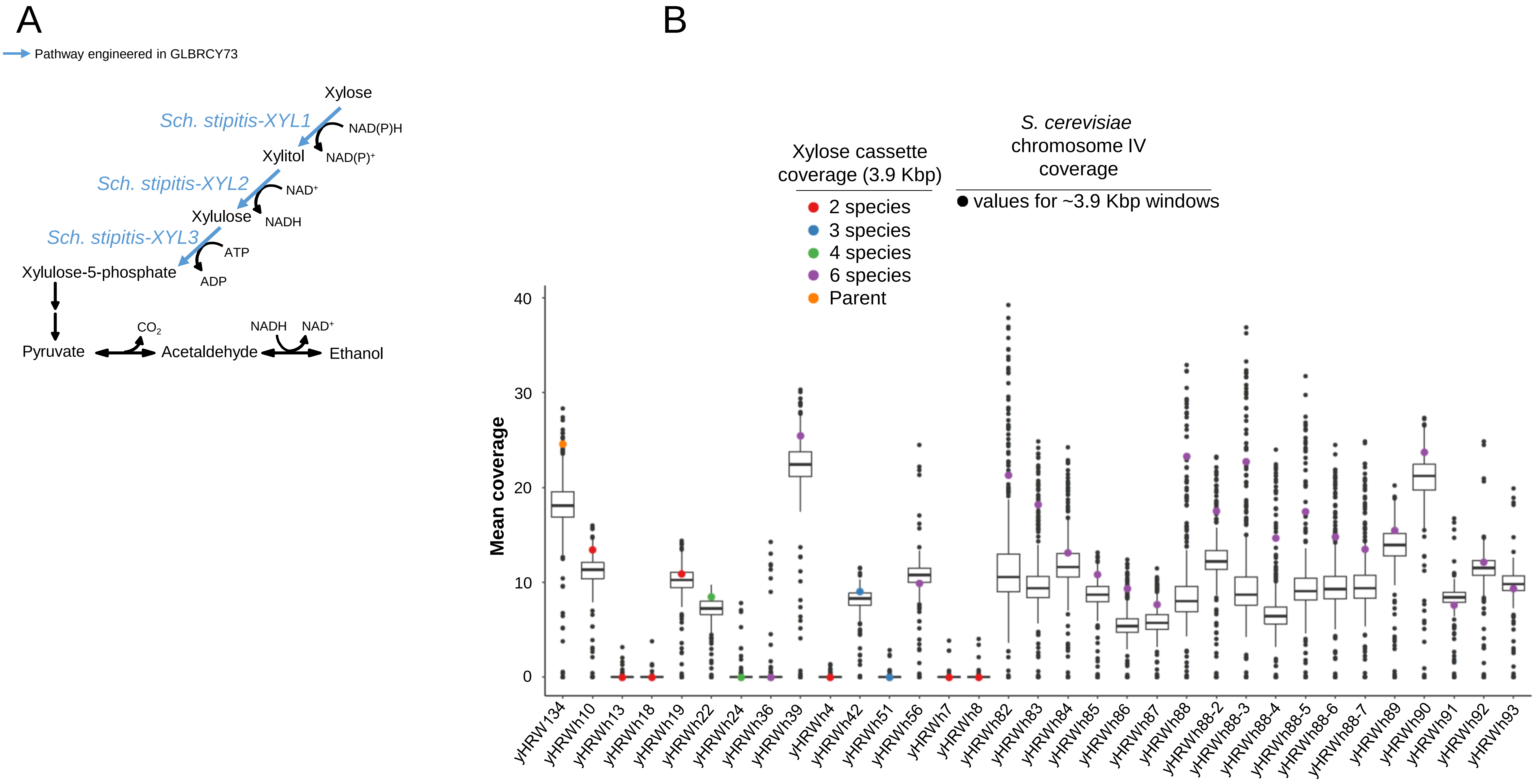

**Supplementary figure 8**

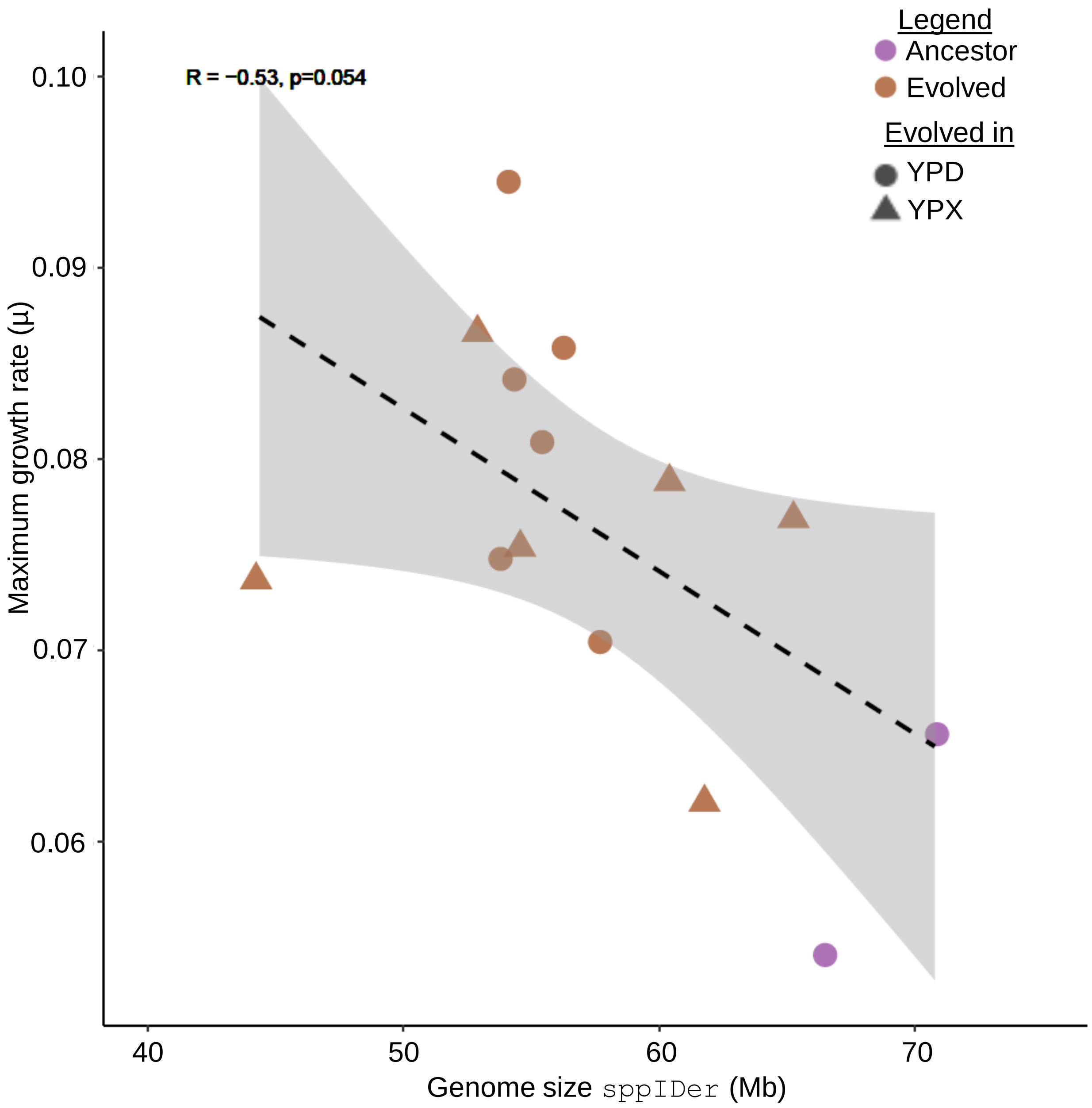
